# Spontaneous thought transitions follow a foraging-like policy on high-order memory patches

**DOI:** 10.64898/2026.09.07.749780

**Authors:** Jiaqi Zhou, Huiting Lei, Ziang Wang, Xia Liang

**Author notes:** **Corresponding author:** Xia Liang Research Center for Social Computing and Interactive Robotics, Harbin Institute of Technology, Harbin 150001, China.

## Abstract

When people engage in spontaneous thought under low external constraint, thought content changes continuously and can shift across topics. However, the cognitive computational principles governing these transitions remain unclear, as does whether similar dynamics emerge in artificial language generation. Here, we formalized spontaneous narrative dynamics using a foraging-inspired framework, constructing an agent based on the marginal value theorem (MVT) that balances exploration and exploitation while selecting among informational resources represented in a participant-specific, high-order patch network. Patches were defined as partially overlapping composite structures containing second-order paths and third-order interactions, with their available resources quantified using multilevel structure values. In a multi-theme free spoken narrative task, the model predicted the temporal organization of human patch activation substantially better than random-shuffle baselines, although the strength of model conformity varied across participant–theme samples. Model-predicted patch transitions corresponded to systematic semantic segmentation and reorganization across multiple representational levels. Transition rates were positively associated with the perceived importance and likelihood of the narrated events. These findings were replicated when the modeled memory environment was enriched with additional hub-related narrative resources in a supplemental experiment. Control-model analyses further showed that patch-network topology, resource depletion history, or either order of structure value alone could explain only part of the observed dynamics; the most stable human prediction required foraging-related decisions guided jointly by multilevel structure values. Narratives generated by large language models (LLMs) also showed detectable conformity with the same framework, but the full model captured LLM content less consistently, while simpler control models accounted for a larger proportion of their patch-level dynamics. Together, these findings support a computational account in which patch-level narrative dynamics are organized by value-guided foraging over structured memory resources, while revealing graded differences in the strength and stability of these dynamics across human and artificial language generation.

## 1. Introduction

Humans spend almost half of their waking lives in an “offline” mode of spontaneous thought. Such self-generated, low-constrained thought, which typically arises when external task demands are low is highly dynamic^1^. Its contents can switch spontaneously among topics. On the one hand, this dynamic flow helps integrate past experiences and simulate or plan possible future scenarios^2,3^. On the other hand, the temporal structure and transition patterns of spontaneous thought also reflect individual differences in personality and mental health^4,5^. At the neural level, spontaneous thought is accompanied by activation of the default mode network (DMN), particularly a subsystem centered on the medial temporal lobe (MTL)^6,7^. Supported by these systems, individuals may continuously retrieve potentially useful information or ideas from long-term memory to construct the content of their thoughts^8,9^. Recent work further shows that abrupt semantic shifts during spontaneous thought elicit neural activity similar to those evoked by external event boundaries^10^. Based on such findings, it has been proposed that the dynamic transitions of spontaneous thought can be understood as a search process over a memory network: thought wanders among nodes that represent memory fragments, with edges reflecting latent associations between them^2,11^. However, it remains unclear what decision processes govern movement through such a memory space and whether the resulting transitions correspond to meaningful boundaries in the organization of internally generated content.

Research on semantic search provides a possible framework for addressing this question. Human search in semantic memory often exhibits foraging-like dynamics, with individuals retrieving several related items within a local “semantic patch” before switching to a different region^12^. Search efficiency in such semantic-foraging tasks is also associated with creative ability^13^, and spatial foraging policies can modulate the temporal organization of creative thought^14^. Foraging behavior is classically framed as an explore–exploit trade-off. According to the marginal value theorem (MVT), a forager should leave the current resource patch when the local rate of gain falls below the average rate available in the environment, thereby optimizing long-term reward returns. This principle has been applied to human decision-making across a range of cognitive tasks^15–18^. In light of these previous work, a critical question arises: whether transitions of spontaneous thought among different memory elements also follow such value-guided stay–leave decisions.

Memory representations are unlikely to consist only of isolated elements or pairwise connections. Multiple memory elements can also participate in high-order relationships, in which three or more elements are jointly related rather than represented only through pairwise connections. Humans can represent such relational organization in knowledge^19^, temporal structure^20^, and social relations^21^, and can infer latent relationships even when they must be reconstructed from observed pairwise relations^22,23^. High-order relationships may also have functional consequences, shaping global network organization and activity propagation and accounting for neural and behavioral variability beyond traditional pairwise connectivity measures^24–26^. Together, these findings suggest that memory representations may contain relational organization extending beyond pairwise associations. We propose that such relational organization can form local memory-resource regions required by a foraging account of spontaneous thought, which we refer to as patches. A patch is not an individual memory element, but a local composite unit that integrates pairwise connections through high-order relational organization. Because patches differ in the amount and organization of the relational content they contain, memory resources are unevenly distributed across the resulting environment. Thought trajectories may therefore be preferentially drawn toward regions rich in memory resources, making their temporal organization difficult to capture using a simple random-walk account.

Such preferential movement implies that different memory resources may carry different values for thought. In human foraging contexts, neuromodulatory systems, including dopaminergic, cholinergic, and noradrenergic systems, help regulate value-based decisions by balancing reward and search costs^27,28^. Decision making is also constrained by the costs of acquiring and processing information^29,30^. Based on these ideas, we propose that spontaneous thought can be viewed as a foraging process guided by informational reward. within a memory environment composed of patches, structural information itself constitutes an internal resource. We quantify the value provided by the composite organization of each patch as its structure value, which serves as the informational reward guiding thought foraging. Within this framework, individuals preferentially enter patches with greater expected structural value and decide whether to stay in or leave the current patch by comparing its remaining local value with the expected value of the broader environment.

Beyond providing a computational account of human spontaneous thought, this foraging framework also offers a principled benchmark for evaluating whether artificial language systems exhibit similar cognitive dynamics. Large language models (LLMs) have demonstrated striking fluency in generating coherent, contextually appropriate text, and their outputs are increasingly used as proxies for human behavior in psychological research^31–37^, raising the question of whether the processes underlying this output also mirror the dynamics of human thought. However, there are theoretical reasons to expect fundamental divergences. The transitions of human spontaneous thought, as characterized above, is driven by internally generated rewards and the structural organization of episodic and semantic memory^2,38^. LLMs, by contrast, generate text through next-token prediction over learned statistical regularities. Whether this process gives rise to transition dynamics functionally analogous to foraging in memory space, and comparable to those observed in humans, remains an open empirical question.

To investigate these questions, we developed an MVT-based model agent operating within a theme-specific patch network reconstructed from free spoken narratives. Participants narrated personally relevant themes across three trials, allowing us to identify recurring memory elements, temporally connected hub-to-hub narrative paths, and high-order patches distributed across trials. The model agent evaluated the structure values of patches and implemented foraging-like patch-entry and patch-leaving behavior according to two strategies: (a) maximizing information acquisition, whereby the agent preferentially entered patches with lower expected information entropy and greater informational complexity; and (b) optimizing the explore–exploit trade-off, whereby the agent decided whether to remain in or leave the current patch according to an MVT-based rule. We evaluated model predictions against the observed temporal organization of patch activation and tested whether predicted patch-level transitions were accompanied by distinctive semantic changes in thought content. We also examined whether transition rates varied with subjective evaluations of the narrated themes. A follow-up two-stage experiment further assessed whether these predictions persisted when the patch network was enriched with additional hub-related narrative content beyond the original free narratives. Finally, we applied the same modeling and semantic analyses to narratives generated by two large language models. Together, these analyses test whether the patch-level dynamics of spontaneous thought conform to foraging-related decisions guided by multilevel structure values.

## 2. Materials and methods

### 2.1 Experimental procedure

Fifty healthy adults participated in a free spoken narrative task (average age 23.3, range from 18 to 30, 33 males and 17 females). Before the narration session, participants completed a “current concerns” questionnaire in which they listed 6–8 ongoing personally relevant events and provided ratings for each event. From the self-reported events, we selected four themes per participant such that they spanned opposite affective valence (positive vs. negative) and opposite subjective importance (high vs. low). During the narration, each trial required participants to speak freely for 15 minutes about anything within the scope of the four selected themes. Participants were instructed to avoid extended silences as much as possible. Each participant completed three trials, separated by 5-minute breaks. Informed consent was obtained from all participants before the experiment, after the experiment, they were compensated with money. The study was approved by the Ethics Committee of Harbin Institute of Technology.

### 2.2 Pre-task questionnaires

All participants completed a set of self-report questionnaires to quantify their appraisals of the 6–8 self-reported current concerns. For each concern, participants rated nine items capturing complementary dimensions of evaluation, including emotional valence toward the event; perceived importance, urgency, controllability, knowledge, and likelihood of event realization; commitment to pursuing the event; expected time to event realization. Ratings were provided on an 11-point integer scale ranging from 0 to 10, with higher values indicating greater intensity or a shift along a bipolar continuum depending on the item (e.g., from “not at all important” to “extremely important,” or from “highly worried, anxious” to “highly happy, pleased”). Participants completed the full nine-item questionnaire for each listed concern. The complete questionnaire is provided in Supplementary Table 1.

### 2.3 Data preprocessing and construction of theme-memory networks

Seven participants were excluded due to low speech volume, strong accent-related transcription failures, or insufficient narrative content, leaving 43 participants for analysis. Audio recordings were transcribed into text using aTrain, which segments speech into text units based on pauses (silent intervals). We organized each participant’s transcripts by theme (the four selected themes) and trial, and retained those themes for which the participant produced content in all three trials. The raw transcripts contained substantial filler words, transcription errors, and semantically uninformative expressions, thus we preprocessed all text segments using the large language model deepseek-V3-0324 with the following fixed user prompt: remove repetitive fillers (e.g., “um,” “uh,” “then”), retain technical/proper terms, preserve narrative details, keep the original meaning while making the narration more concise, maintain the temporal order, and correct typos/mistranscriptions.

We treated each preprocessed text segment as a thought unit, and each thought unit corresponded to a node in the theme-memory network for a given theme. For each theme, nodes were connected in their temporal order of occurrence. We then merged semantically nodes with highly similar semantics using the embedding model moka-ai/m3e-base, within a strict threshold (cosine similarity in the range 0.86–0.90). We constructed one theme-memory network per participant per theme (as shown in Figure 1A).

**Figure 1.**
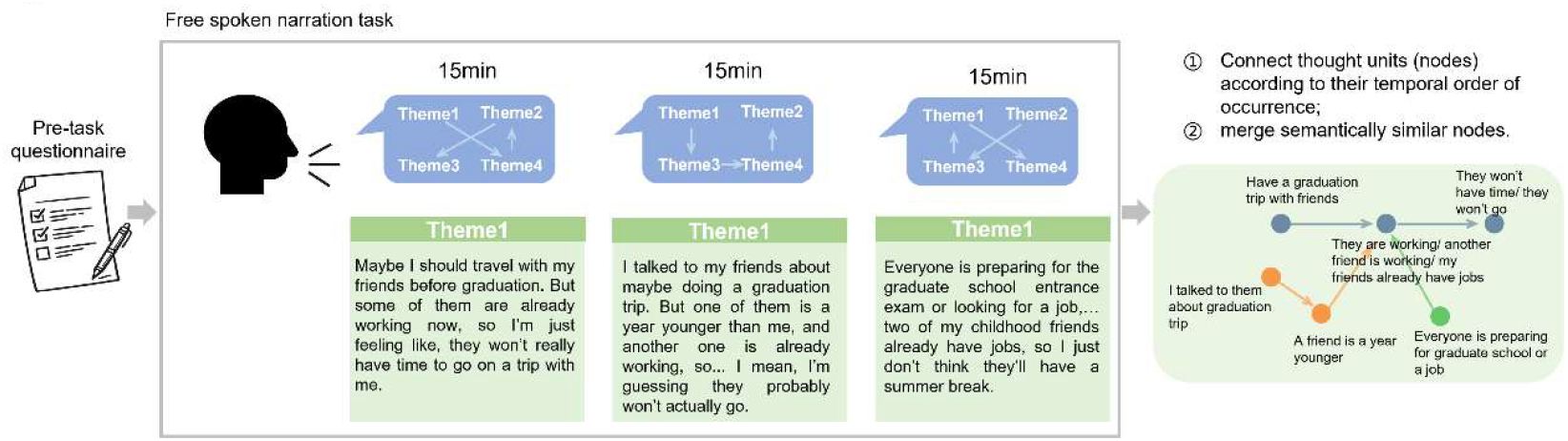
Experimental overview. Before the free spoken narration task, participants completed self-reported questionnaires about their current concerns. During the task, they spoke freely for 15 minutes in each trial while staying within four themes selected from their listed concerns, Part of the narrative samples are shown in the green “Theme1” boxes. Preprocessed text segments were treated as thought units, we connected thought units according to their temporal order and merged semantically similar units, to construct a theme-memory network (the network in the green box of the right column is constructed from “Theme1”).

### 2.4 The foraging environment on the human theme-memory network

#### The human theme-memory network for a particular theme

We used each participant’s narrative content for a given theme to construct an individual-level memory network, intended to capture that participant’s theme-specific pool of thinking resources. Across repeated narrations, participants tend to return to a small set of stem sentences that convey the core content of the theme, while progressively enriching the narrative by adding additional details from the resource pool^39^. We treated all preprocessed text segments across trials as thought units and modeled them as nodes in the theme-memory network (Figure 1, right column), each node could reflect a mixture of semantic and episodic content^40^. Nodes were connected according to their temporal order of occurrence in speech, and highly similar nodes were merged based on their semantic similarity, yielding an individual-level theme-memory network as illustrated in Figure 2A.

**Figure 2.**
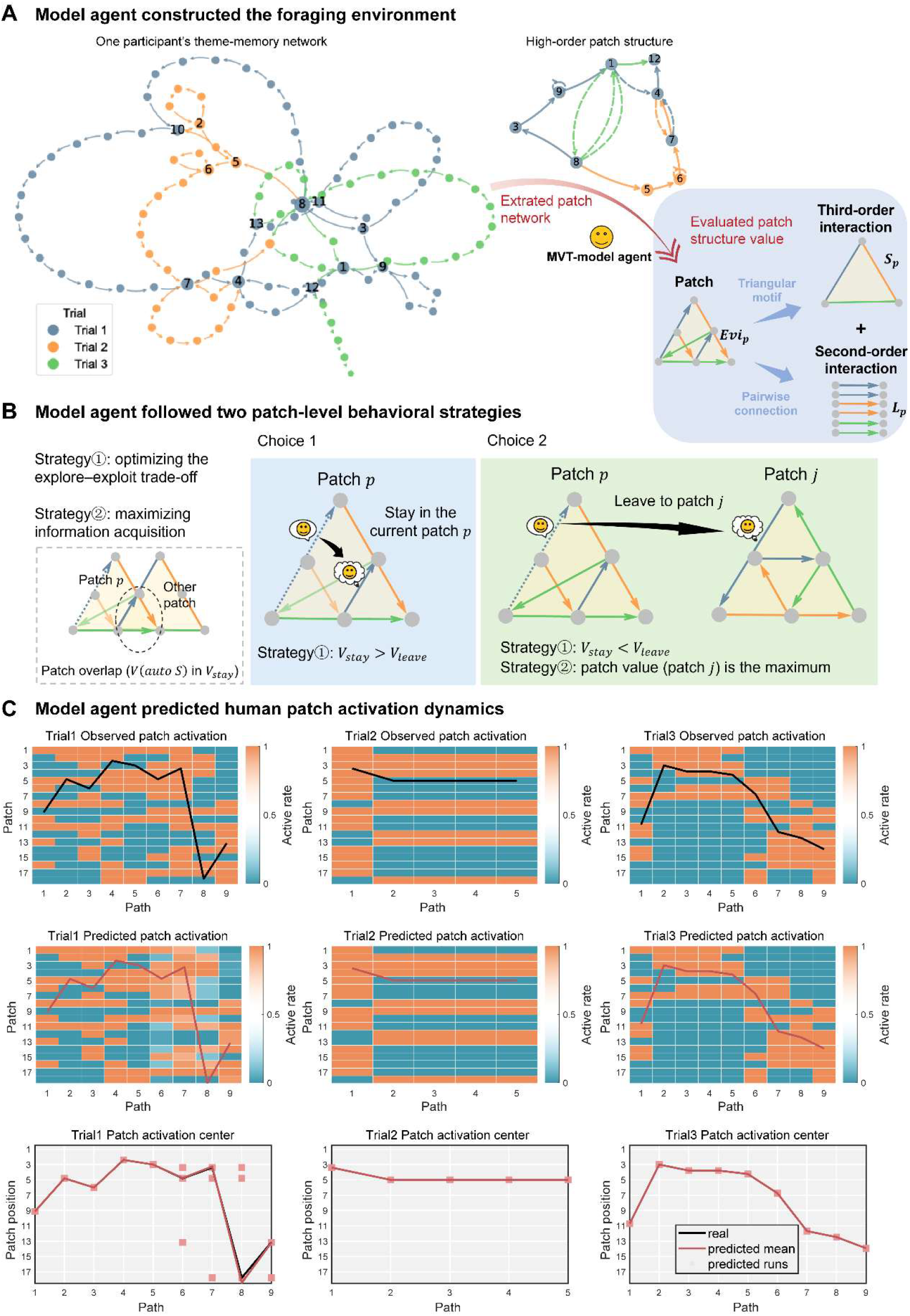
Working pipeline of the MVT-based model agent and its prediction for an example participant–theme sample. The MVT-based model agent constructed a foraging environment, evaluated the multilevel structure values of patches, and selected its behavior within the patch network (A–B). The resulting model prediction captured the patch-level activation pattern observed in the human narrative (C). (A) Construction and evaluation of the patch-based foraging environment. The agent constructed patches composed of multiple second-order interactions and one third-order interaction. It then evaluated the informational resources provided by these structures, with the value of each patch reflecting the weighted benefits associated with foraging over structures of different orders. (B) Patch-entry and patch-leaving strategies. The agent’s behavior was governed by two principles: maximizing information acquisition and optimizing the explore–exploit trade-off. At each decision point, the agent compared the value of staying in the current patch, *V*_stay_, with the value of leaving, *V*_leave_. When *V*_stay_ > *V*_leave_, the agent remained in the current patch and proceeded to another hub-to-hub path within that patch. Otherwise, it left the current patch, evaluated the candidate patches, and proceeded along a hub-to-hub path belonging to the patch with the highest expected value. Dashed lines in Choices 1 and 2 indicate the current hub-to-hub path. Because patches could overlap, traversal of a single hub-to-hub path could simultaneously activate multiple patches (left column), contributing to the co-activation patterns shown in (C) and affecting the calculation of *V*_stay_. (C) Observed and model-predicted patch activation and patch position centers. Top: observed patch activation matrix for the example human sample; the black trace denotes the observed patch position center. Middle: patch activation matrix predicted by the MVT-based model agent operating on the same theme-memory network; the red trace denotes the predicted patch position center. Because one path could belong to multiple overlapping patches multiple patches could be co-active within the same matrix column. Bottom: comparison between the observed and predicted patch position center curves. Their close correspondence produced a small distance. Because value-maximizing patch selection required the agent to retrieve the remaining values of all candidate patches, retrieval noise was introduced at the relevant decision points. Red squares indicate the possible predicted patch position centers at each hub-to-hub path across 2000 simulations.

To characterize positional change in thought transition, we did not use the original nodes in the theme-memory network as position anchors. Instead, we used repeatedly revisited hub nodes as reference points for tracking positional change. This was motivated by the fact that the text segments of original nodes more often reflect local narrative details or fluctuations in linguistic organization, and are therefore more susceptible to individual differences in speaking style and narrative habit. By contrast, hub nodes are more likely to represent relatively stable semantic attractors within the theme. Using hub nodes as positional anchors both reduces local noise and maps the narrative process onto a more cognitively meaningful representation of changes in thought position. In a theme-memory network constructed by temporal order of occurrence, a non-terminal node that is visited only once typically has a degree of 2. Exceeding this baseline (i.e., degree ≥ 3) usually indicates that the node was revisited and linked to multiple narrative contexts. To ensure that the extracted hub nodes more stably captured the important informational structure of the network, we defined nodes with degree ≥ 4 as hub nodes (numbered in Figure 2A), and edges connecting pairs of hub nodes were treated as paths along which thought traverses the theme-memory network.

#### The patch structure in the human theme-memory network

To model how human thought switches between local regions of memory resources within a theme-memory network, we first defined the patch structure of the network. In natural environments, resource-rich areas are separated by physical distance, and travelling between them incurs costs for the forager. Analogously, we assumed that memory resources within a participant’s theme-memory network were not uniformly distributed but were concentrated within a set of local structures, with transitions between these structures incurring a cognitive cost. We refer to these local resource-rich regions of the memory network as patches. Corresponding to physical distance in natural environments, patches in the memory network are separated by structural distance, defined as the extent to which they are structurally distinct from one another.

From the perspective of high-order interaction networks, relationships among nodes can be organized into a hierarchical system of inclusion. When two high-order structures are not directly related through structural inclusion, moving from one to the other requires traversal through additional intermediate relational levels^41,42^. This traversal can be regarded as the structural distance between the two structures. Motivated by this principle, we defined each patch as comprising one third-order interaction and the second-order interactions contained within it. Under this definition, moving from one patch to another requires a hierarchical transition from a third-order interaction to its constituent second-order interactions and then into another third-order interaction, thereby creating a relatively large structural distance between patches.

More specifically, let *N*_1_, *N*_2_, *N*_3_ denote the sets of hub nodes in the three narration trials. Within each trial, a pairwise connection between two hub nodes was defined as a second-order interaction. In the context of continuous human narration, such an interaction was also referred to as a ***hub-to-hub path***. The sets of second-order interactions obtained from the three trials were denoted *E*_1_, *E*_2_, *E*_3_, respectively.

Suppose that interactions *e*_1_, *e*_2_, *e*_3_ are selected from *E*_1_, *E*_2_, *E*_3_, respectively. If these three interactions jointly form a closed triangular motif, they are defined as a third-order interaction (see Methods 2.5 for details). Consistent with the three-trial design, we required each third-order interaction to be formed by second-order interactions drawn from all three narration trials rather than from a single trial. This requirement allowed the third-order interaction to represent relationships preserved across repeated narrations. Local connections observed within only one trial may primarily reflect trial-specific narrative details. By contrast, a closed motif that can be completed only through the combination of *e*_1_, *e*_2_, *e*_3_ is more likely to reflect a relatively stable organizational scaffold of theme-related memory.

Accordingly, we defined a patch in the memory network as a composite structure consisting of one third-order interaction formed by temporally connected interactions drawn from *E*_1_, *E*_2_, *E*_3_, together with all second-order interactions contained within that third-order interaction, as illustrated in the right column of Figure 2A.

As free narration unfolded, the thought content expressed along each hub-to-hub path could activate multiple patches. As illustrated in Figure 2C, whenever a participant narrated along a hub-to-hub path, several patches could be co-activated because the same path could be contained in multiple overlapping patches. To characterize the temporal movement of patch activation and identify patch-level entry and leaving dynamics associated with the model’s foraging-related strategies, we calculated the patch position center for each narrated path (see Methods 2.6). This measure summarized the location of the activated patches within the ordered patch space (Figure 2C, black trace).

### 2.5 A behavioral model of optimal foraging strategies on the theme-memory network

#### Patch selection strategy for maximizing information acquisition

To test whether human thought traversed theme-memory networks in a manner consistent with optimal foraging theory, we constructed a model agent whose decisions and movements followed MVT-based foraging strategies. We refer to this agent as the ***MVT-based model agent***. The agent first constructed a foraging environment composed of the patches defined above. It then encountered multiple candidate resource-rich patches and determined which patch to enter.

Patch-entry decisions were guided by the structure value of each patch, which quantified the amount of harvestable informational resources embedded in the high-order patch network. We denoted the structure value of patch *p* as *Evi*_*p*_. To maximize its total informational gain, the agent preferentially entered the patch with the highest *Evi*_*p*_. At each decision time *t* in narration trial *j*, when the agent was not currently located within any patch, it selected the patch with the greatest expected value. The patch choice 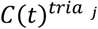 was defined as:

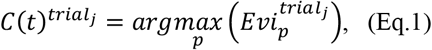

We further decompose the patch value 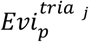 into contributions from high-order interaction structures. Specifically, the expected structure value of patch *p* in trial *j* is modeled as a combination of (1) the third-order interaction resource 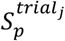 and (2) the sum of the second-order interaction resources across the three trials 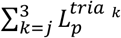,

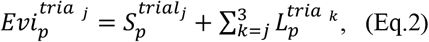

To quantify the resources 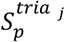 and 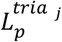, we treated the semantic contribution of isolated nodes as equivalent and focus on the additional structural benefits introduced by high-order structures^43^. Let *b*_0_ denote the baseline contribution of a single hub node in the theme-memory network. When hub nodes were combined in pairs to form a second-order interaction, the benefit that an agent obtained by visiting such a second-order interaction is *l*_0_ = *b*_0_ ∗ (1 + *δ*_2_), where *δ*_2_ represented the rate of diminishing marginal effect of the node contribution within second-order interactions.

The third-order interactions are the closed loops constructed by temporally connected by second-order interactions from all three narration trials. For simplicity, we quantified the structure value of a third-order interaction as the additional structural benefits of a triangular motif formed by three second-order interaction structures (as depicted on the right of Figure 1B). When an agent accessed a hub node, the benefit it obtained is 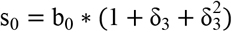, where δ_3_ represented the rate of diminishing marginal effect of the node contribution within third-order interaction structure.

In addition, remaining narration trials may influence how the agent evaluates patch value. We therefore allowed the node contribution to vary across trials, denoted 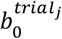. We considered the three narrative trials as three vertices. Each pair of trials formed a second-order trial interaction, and all three trials together formed a third-order trial interaction. Under this formulation, for the beginning of trial 1, the remaining three narrative trials consisted of one third-order trial interaction and three second-order trial interactions. For the beginning of trial 2, the remaining two narrative trials were composed of two second-order trial interactions. For the beginning of trial 3, the remaining one narrative trial was made up of one second-order trial interaction. The node contributions under the three trials are as follows,

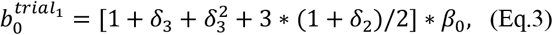

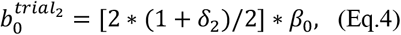

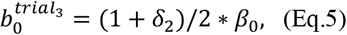

Where the *β*_0_ represented the node contribution constant.

Accordingly, the third-order and second-order resources within patch *p*, 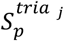 and 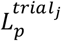, are defined as,

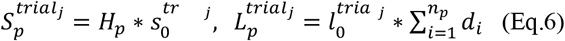

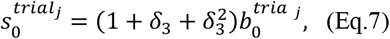

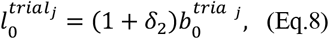

where *H*_*p*_ and *d*_*i*_ represented the weights assigned to different structures when the agent estimates the structure value in patch *p*, respectively. *n*_*p*_ denoted the remaining number of second-order interactions within patch *p. H*_*p*_ reflected the structural complexity of the third-order interaction structure. It was measured by the spectral entropy of the patch *p* and exhibited a negative-exponential decay profile. *d*_*i*_ is the normalized degree of key node *i*, which follows a normal distribution. Together, Equations (1), (2), (6) suggested that when the forager treats multi-order structural information as patch value and uses this value as decision evidence for patch entry.

#### Patch transition strategy for optimal explore-exploit trade-off

During foraging, under a finite energy budget, the agent must trade off exploitation of the current patch (continued harvesting) against exploration of alternative patches, which incurs energetic costs and yields no reward during transition. The MVT formalizes the optimality of this trade-off: the forager should continue exploiting the current patch as long as the expected reward rate within the patch exceeds the background reward rate. Otherwise, it should leave the patch and explore other patches.

We considered that during each foraging step, the agent obtains information resource associated with a second-order interaction. At foraging step *t*, the agent decides whether to transition by comparing the value of staying in the current patch versus leaving it^44^. The value of staying in the current patch, denoted as *V*_stay_, is defined as the agent’s expected structure values within the current patch at the next foraging step (t+1):

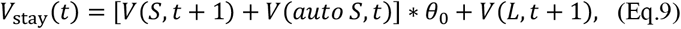

Where *V*(*S, t* + 1) denoted the expected decrement of the third-order interaction *S* during the (*t* + 1)-th foraging step. Here, *θ*_0_ representd the foraging proportion parameter. The variable *V*(*S, t* + 1) follows a negative exponential decay, given by the formula −*a* ∗ exp[(*t* + 1)/*b*], where *a* and *b* control the decay profile of *S*. In addition, *V*(*L, t* + 1) representd the denotes the expected resource available from a second-order interaction *L* during the (*t* + 1)-th foraging step. That is, it is the expected decrement of *L* in this foraging step, expressed as 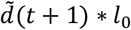. The term 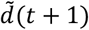 represented the expected weight induced by the degree of the hub nodes and followed a normal distribution 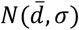. Since the theme-memory network is constructed based on the content of individual narratives, we assumed that the individual has full knowledge of the decay dynamics of *S* and the distribution properties of *L*.

Notably, different patches can overlap, that is, they shared same second-order interaction structures. Consequently, at each foraging step the agent may also obtain additional benefits attributable to the decay of third-order interactions in other patches that overlap with the current one. The This overlap-driven benefits influenced the agent’s estimate of *V*_stay_(*t*), corresponding to the *V*(*auto S, t*) component in Equation (9).

The value of leaving the current patch, *V*_leave_(*t*), is defined as the background reward rate, estimated by the average resource acquisition rate across all previous foraging steps,

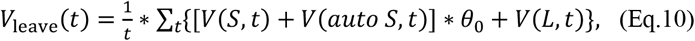

When *V*_stay_ < *V*_leave_, the agent would decide to leave the current patch and initiates a new patch-entry decision process.

### 2.6 Computation of model predicted performance

For each observed hub-to-hub path in a participant’s narrative, the corresponding MVT-based model agent completed one foraging step. The observed sequence of hub-to-hub paths determined the number and temporal order of the simulation steps, whereas the model’s patch-level decisions determined which patches were activated at each step. Because patches could overlap, a single hub-to-hub path could be contained in multiple patches, allowing several patches to be co-activated during the same foraging step.

For each participant-theme sample and narration trial, we constructed an observed patch activation matrix, ***A***^obs^, in which rows represented patches and columns represented the temporally ordered hub-to-hub paths. An element 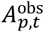 was assigned a value of 1 if the observed path at step *t* belonged to patch *p*, and 0 otherwise. Similarly, for each model simulation *s*, we constructed a predicted patch activation matrix, ***A***^pred,*s*^, in which 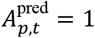 indicated that patch *p* was activated at step *t* in that simulation. Multiple elements within the same column could therefore equal 1 when overlapping patches were co-activated. Under value-maximizing patch selection, the agent must retrieve the remaining value of all patches from memory. We therefore introduced retrieval noise. At the *t*-th decision step, the agent may mistakenly recall a patch’s value as its remaining value at an earlier time *t* − *Δt*, where *Δt*~*Boltzmann*(*β, t*). Such misrecall can propagate and produce cascading effects on subsequent patch transitions^45^. To account for this, we simulated the model agent 2,000 times, and averaged the resulting patch activation matrices across simulations, leading to the activation patterns shown in Figure 2C.

For visualization, the predicted patch activation matrix was obtained by averaging the binary activation matrices across the 2000 simulations:

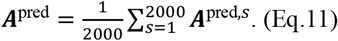

Each element of ***A***^pred^ therefore represented the proportion of simulations in which a given patch was activated at a given hub-to-hub path. Importantly, model performance was not calculated by directly comparing this averaged matrix with the observed matrix. Instead, cosine similarity and patch position distance were calculated separately for each simulation and were subsequently averaged across the 2000 simulations.

#### 2.6.1 Cosine similarity between observed and predicted patch activation matrices

For each simulation *s*, we quantified the similarity between the observed and predicted patch activation patterns by vectorizing the two matrices and calculating their cosine similarity,

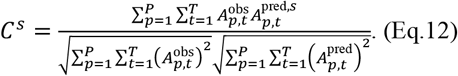

This measure simultaneously captured whether the model activated the appropriate patches and whether those activations occurred at the corresponding positions in the observed path sequence. Higher cosine similarity indicated greater agreement between the observed and model predicted patch activation matrices. The cosine-similarity performance of a participant-theme sample in a given trial was defined as the arithmetic mean across the 2000 simulations: 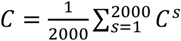.

#### 2.6.2 Patch position center and circular distance

To characterize the movement of human narratives and model simulations through the global patch space, we further calculated a patch position center for each hub-to-hub path. Because patches do not have an intrinsic one-dimensional order, all *P* patches were assigned equally spaced positions on a circular ring. For a given patch order, the angular position of patch *p* was denoted by θ_*p*_. The position center ϕ_*t*_ at path step *t* was decoded as the weighted circular mean of the activated patch positions^46^,

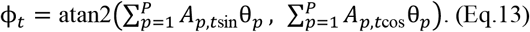

This procedure was applied separately to the observed activation matrix and to the predicted activation matrix from single-theme simulation, leading to an observed patch position trajectory and a predicted patch position trajectory across the observed hub-to-hub paths.

Because the resulting position centers depend on the arbitrary assignment of patches to circular positions, we repeated the calculation using 1000 random permutations of patch order. Within each permutation, the same patch order was applied to both the observed and predicted activation matrices. For simulation *s*, random patch order *k*, and path step *t*, the circular distance between the observed and predicted position centers was calculated as the shortest angular separation,

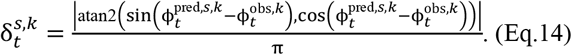

Dividing by π normalized the circular distance to the range from 0 to 1. For each simulation and patch-order permutation, distances were first averaged across all hub-to-hub paths in the trial. The resulting values were then averaged across the 1000 random patch orders to obtain one patch position distance for each simulation. Finally, the sample-level distance was calculated by averaging these distances across the 2000 simulations. Lower values indicated closer correspondence between the observed and model predicted patch-position trajectories.

#### Random shuffle performance

To determine whether the MVT-based model predicted the correspondence between patch activations and specific positions in the narrative path sequence beyond chance, we constructed a random shuffle baseline. For each participant-theme sample and trial, we randomly permuted the columns of the observed patch activation matrix. This procedure preserved the complete set of patches co-activated within each column, as well as the overall patch activation frequencies, but disrupted the correspondence between each group of activated patches and its original hub-to-hub path position.

For each shuffled matrix, cosine similarity and patch position distance were calculated relative to the original observed activation matrix using the same procedures as those applied to the MVT-based model. In particular, patch position distance was again averaged over 1000 random permutations of patch order. The random-shuffle procedure was repeated 2000 times, and the resulting cosine similarities and distances were averaged to obtain the random shuffle performance for each participant-theme sample and trial. Thus, the random shuffle baseline tested whether the model captured the temporal assignment of patch-activation patterns to the observed paths, rather than merely reproducing the overall distribution of patch activations.

#### Group-level model performance

Group-level performance was calculated from the scalar performance values obtained for individual participant-theme samples, rather than by pooling or averaging their patch activation matrices. For each trial, the group-level cosine similarity and patch position distance were calculated as the arithmetic means of the corresponding sample-level values across all included participant-theme samples. Random-shuffle performance was aggregated in the same manner.

For group-level visualization, observed and predicted patch activation matrices with different dimensions were projected onto a common normalized coordinate space and resampled before averaging across samples (Supplementary Methods S1).

### 2.7 Human and Qwen grouping based on model-prediction performance

#### Human H1/H2 grouping

Human participant–theme samples were characterized by a six-dimensional model-performance profile consisting of random-shuffle-standardized cosine similarity and patch position distance across the three trials. Samples were first clustered according to these profiles, and the resulting clusters were subsequently grouped according to their similarity to the cluster showing the strongest model-conformity profile, producing the higher-conformity H1 and lower-conformity H2 groups.

#### Qwen Q1/Q2 assignment

Qwen-generated samples were represented in the same six-dimensional performance space but were not independently clustered. Instead, each sample was assigned to its nearest original human cluster and labeled Q1 or Q2 according to whether that cluster belonged to H1 or H2. Full clustering and classification procedures are provided in the Supplementary Methods S2.

### 2.8 Identification of model-predicted patch transitions

Predicted patch activations included both patches actively selected by the model agent and patches automatically co-activated because of overlap among their constituent paths. To isolate transitions generated by the model’s foraging-related decisions, we retained only the actively selected patch at each path and represented these selections in an active patch selection matrix.

For both the model-predicted and empirical trajectories, we tracked the evidence value 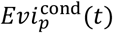 of each patch *p* at each path *t*, where cond ∈ {pred, emp}. The two conditions used the same structure-value and depletion rules but differed in the activation history used to update the remaining patch values: 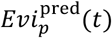 was updated according to the model-predicted patch activations, whereas 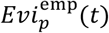 was updated according to the observed patch activation matrix, including both observed activations and overlap-driven co-activations. For either condition, the choice outcome was defined as the patch with the greatest current evidence, 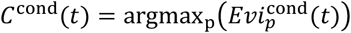. A choice update was denoted by *ΔC*^cond^(*t*), which equaled 1 when *C*^cond^ (*t* + 1) *≠ C*^cond^ (*t*) and 0 when the same patch remained favored across the two consecutive paths.

Model-predicted transition steps were identified using two criteria. **(i) Leave condition**. At path *t*, the currently selected patch entered the leave region, such that *V*_stay_(*t*) < *V*_leave_(*t*), while still containing available resources. **(ii) Choice-update condition**. The predicted choice outcome changed across the corresponding path boundary, such that *ΔC*^pred^(*t*) = 1. A path boundary satisfying both conditions was classified as a predicted transition step. Boundaries at which the current patch exhausted all resources available in the trial were also classified as transitions when the subsequent path involved a different patch, even though the leave condition was not satisfied. Conversely, a choice update that occurred before the leave condition emerged was not classified as an MVT-generated transition. All remaining boundaries were defined as predicted non-transition steps.

Because the actively selected patch cannot be directly observed in human narratives, the empirical evidence provided a corresponding measure based on the observed activation history. Specifically, *ΔC*^emp^(*t*) = 1 indicated that the patch with the greatest remaining structure-value evidence changed between two consecutive paths in the observed evidence *C*^emp^(*t*). We therefore compared the empirical choice-update rate between model-predicted transition and non-transition steps to test whether boundaries generated by the model’s active patch-transition process were also associated with changes in the patch favored by the empirical patch evidence.

### 2.9 Semantic dynamics associated with model-predicted patch transitions

Using the model-predicted transition steps identified above, we next examined whether these decision boundaries were associated with systematic changes in the semantic content of the narratives. We first defined transition-related and patch-internal comparison sets and then performed four complementary analyses that characterized semantic relationships at different levels of narrative organization, ranging from the overall organization of hub nodes to the local relationship between consecutively narrated paths.

#### 2.9.1 Definition of patch-internal and transition-related comparison sets

We defined two sets of inner-patch paths. ***Transition-leaving paths*** were defined as the paths immediately preceding model-predicted patch transition steps identified according to the criteria described above. ***Inner-patch paths*** included all hub-to-hub paths contained within patches after the transition-leaving paths had been excluded. Because overlapping patches could contain the same path, duplicate semantic values could occur in this set. The ***inner-patch paths control*** removed these duplicated contributions, such that each unique inner-patch path was included only once. The principal analyses reported in the main text compared ***transition-leaving paths*** with both the ***inner-patch paths*** and the ***inner-patch paths control***, these analyses were conducted at the hub-node level.

#### 2.9.2 Analysis 1: Order-independent hub-node similarity

Analysis 1 examined the overall semantic organization of hub nodes without preserving their temporal adjacency. Within each comparison condition, we calculated the cosine similarity between all eligible pairs of hub nodes associated with the corresponding paths and averaged these pairwise values for Transition-leaving paths and Inner-patch paths/Inner patch paths control. This analysis therefore tested whether the overall pairwise similarity among hub nodes differed between the interior of model-defined patches and the path steps at which the model agent left the current patch.

#### 2.9.3 Analysis 2: Sequential hub-node similarity

Analysis 2 retained the temporal connections between hub nodes. For each hub-to-hub path, we calculated the semantic similarity between the two temporally connected hub nodes defining that path. These sequential hub-node similarities were then averaged separately for the inner-patch and transition-leaving conditions. Thus, whereas Analysis 1 characterized the overall semantic similarity among hub nodes after temporal adjacency was disregarded, Analysis 2 specifically assessed the local semantic continuity between consecutively connected hub nodes in the narrative sequence.

#### 2.9.4 Analysis 3: Sequential path-to-path similarity at patch transitions

Analysis 3 examined semantic changes between two consecutive paths. For each temporally adjacent path pair (*t, t* + 1), we removed the hub nodes shared by the two paths and calculated the mean cross-path similarity between all remaining hub nodes in the two paths, which prevented the similarity estimate from being artificially increased by nodes that were present in both paths.

Consecutive path pairs were classified as transition pairs when the boundary between them had been identified as a model-predicted transition step; the remaining eligible consecutive pairs were classified as non-transition pairs. Analysis 3 therefore tested whether the semantic relationship between consecutively narrated paths differed across model-predicted transition and non-transition boundaries.

#### 2.9.5 Analysis 4: Sequential versus non-sequential path-pair similarity

The MVT-based model predicted whether the agent stayed in or left the current patch, but it did not predict the specific identity of the next narrated path. We therefore conducted an additional analysis to examine whether the observed selection of the next path exhibited a systematic semantic pattern independently of the model-predicted patch transition.

For each path *t*, the sequential path pair was defined as (*t, t* + 1). Non-sequential comparisons paired path *t* with other paths in the same trial that were neither path *t* itself nor its immediately subsequent path *t* + 1. Path-to-path similarity was calculated using the same shared-node removal and cross-path averaging procedure as in Analysis 3. We then compared the semantic similarity of sequential path pairs with the average similarity of the corresponding non-sequential path pairs. Analysis 4 therefore tested whether the path that was actually narrated next was more or less semantically similar to the current path than were other paths occurring elsewhere in the same narrative trial.

### 2.10 Follow-up two-stage experiment

To test whether the memory resources captured by the original free narratives represented only a subset of those potentially available within each theme, we recruited an additional 13 healthy participants to complete a follow-up two-stage experiment. Phase 1 was identical to the free spoken narrative task used in the main experiment. In Phase 2, participants were presented with individualized cues extracted from their Phase 1 theme-memory networks and were asked to freely narrate any thoughts elicited by each cue while remaining within the scope of the corresponding theme. After preprocessing the two-stage narrative data, the final analysis included 11 participants and 25 eligible participant–theme datasets. Detailed procedures for cue selection and the two-stage experiment are provided in Supplementary Methods S3.

Phase 2 narrative content was used to identify additional hub-to-hub paths that could be mapped onto existing Phase 1 patches. These expanded paths enriched the candidate memory resources available within the existing patch environment but did not create new patches, were not incorporated into the observed Phase 1 narrative trajectory, and did not directly generate patch activations. The MVT-based model was then rerun on the unchanged Phase 1 path sequence using the enriched patch resources. Detailed procedures for expanded-path construction and model implementation are provided in Supplementary Methods S3.

### 2.11 Control models

To determine whether the predictive performance of the MVT-based model depended on its foraging-related decision rules and multilevel structure-value representations, we constructed four principal control and ablation models: a patch-semantic random walk model, a No S model, a No L model, and a Depletion history model. All models were implemented on the same theme-specific patch networks and used the same trial-specific patch-path membership structure and observed path sequences as the full model. Each model was simulated 2000 times per participant-theme sample and evaluated using the same cosine similarity and patch position distance measures described in Section 2.6. Additional random-walk variants and an Equal L control are described in the Supplementary Methods S4.

#### Patch-semantic random walk model

The Patch-semantic random walk model tested whether patch-level dynamics could be reproduced from relationships among patches without explicit foraging-related value evaluation or stay-leave decisions. Patch-to-patch transition probabilities were determined by the mean semantic similarity between the hub nodes contained in each pair of patches. At each path, the next primary patch was sampled from the transition probabilities over patches that still contained available paths. As in the full model, paths were depleted after being visited, and a path shared by multiple patches could co-activate all corresponding patches. The model did not compute patch structure value, *V*_stay_ and *V*_leave_.

#### No S model

The No S model tested the contribution of third-order structure value. The original patch definitions and overlap structure were retained, but the third-order component *S* and the overlap-related *V*(*auto S, t*) contribution were removed. Patch entry and stay–leave decisions therefore depended only on the second-order path-value component *L*, while paths continued to be processed and depleted as in the full model.

#### No L model

The No L model tested the contribution of second-order path value. All path-specific *L* values were set to zero, whereas the original patch definitions, overlap structure, third-order value *S*, overlap-related value, and foraging-related decision rules were retained. Patch entry and stay–leave decisions therefore depended only on the remaining *S*-related components.

#### Depletion history model

The Depletion history model tested whether cumulative patch activation and resulting value depletion were sufficient to explain the observed patch transition patterns in the absence of structure value. Both *S* and *L* were removed, and all patches were assigned an initial value of 1. The value of patch *p* at path *t* was defined as 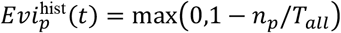, where *n*_*p*_(*t*) denotes its cumulative number of previous activations and *T*_all_ is the total number of paths across the three trials. Both primary activations and overlap-driven co-activations contributed to depletion, which accumulated across trials. When a new patch was entered, the model selected the eligible patch with the highest remaining history-dependent value. Because the model did not evaluate structure value, individual paths no longer differed in the structural reward they provided. Consequently, activation history could still guide which patch to enter, but structural information could no longer inform whether the model should remain in or leave the current patch.

## 3. Results

### 3.1 Working pipeline and prediction target of the MVT-based model agent

We recruited 50 participants to complete repeated free spoken narratives about personally relevant themes, providing temporally ordered sequences of thought content across three trials (Figure 1). After preprocessing, each participant–theme sample was represented as a theme-memory network in which recurring hub nodes and temporally connected hub-to-hub paths captured the observed narrative scaffold.

On this scaffold, we constructed an MVT-based model agent that operated within a participant-specific patch environment (Figure 2A). Patches represented local concentrations of memory resources organized through high-order relationships across repeated narrations. Each patch was assigned a dynamic structure value, which combined second-order path value and third-order structure value. As the observed hub-to-hub path sequence unfolded (Figure 2C), patches containing the current path were activated and their remaining structure values were gradually depleted. Because patches could overlap, a single narrated path could also co-activate multiple patches and consume their structure values.

The agent followed two patch-level behavioral strategies. When exploiting a patch, it applied an MVT-based stay-or-leave rule that balanced continued resource acquisition from the current patch against exploration of alternative patches. When it was not located within a patch, it entered the patch with the highest current structure value to maximize potential information acquisition. Crucially, the model was not designed to predict the identity of the next hub-to-hub path. Instead, the observed path sequence specified the temporal trajectory along which patch values were updated, whereas the model predicted which patch was actively selected and whether that selection changed across consecutive paths. The model therefore produced two related but distinct outputs: a predicted patch activation matrix that included overlap-driven co-activation and a decision-level sequence of active patch selections from which patch-level transitions could be identified.

### 3.2 Human patch activation and transition dynamics are consistent with optimal foraging strategies

#### Operational definitions and an illustrative sample

We first formalized how patch activations and transitions were defined and measured (see Methods for full details). For each observed hub-to-hub path, we constructed an observed patch activation matrix, indicating the patches engaged by participants’ narration, and a predicted matrix, indicating the patches engaged by the model agent when traversing the same network. Because one path could belong to multiple overlapping patches, several patches could be co-active within the same matrix column. To separate genuine patch selection from this overlap-driven co-activation, the predicted activations were decomposed into actively selected activations according to the model’s foraging strategies, and automatically induced co-activations arising solely from patch overlap (Figure 3B). A patch transition was defined as a change in the actively selected patch between two consecutive paths that satisfied the model’s MVT-based leave condition together with a subsequent choice update (Figure 3C); two exception rules, concerning resource exhaustion and premature choice changes, are described in Methods. Path boundaries meeting these criteria were labeled as predicted transition steps, and all remaining boundaries as predicted non-transition steps.

**Figure 3.**
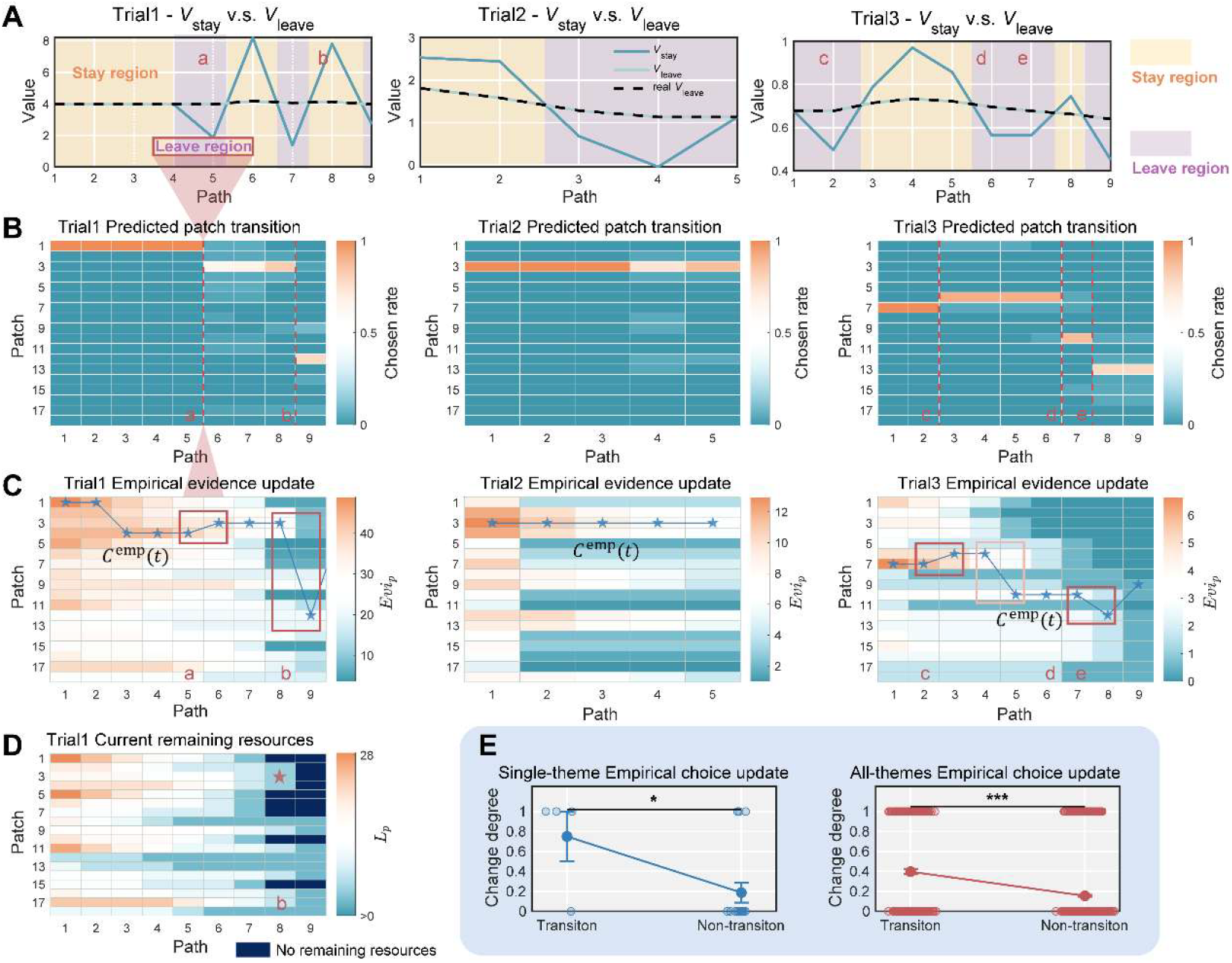
Two foraging strategies underlying patch transitions. (A) The values of staying in the current patch (*V*_*stay*_) and leaving to alternative patches (*V*_leave_) across thought transitions. Yellow and purple shaded regions denote the leave region (where leaving is permitted) and stay region (where staying is preferred), respectively, for the corresponding paths. Blue solid lines denote the model-evaluated value, and black dashed line is the human-evaluated *V*_leave_ (B) Model-predicted active patch selection matrices after removing activations attributable to overlapping of patches at each hub-to-hub path. Dashed lines and labeled points (a–e) mark the agent’s transition steps under the two strategies. An active transition occurs between paths t and t+1 when two conditions were satisfied: (i) Leave condition: the current hub-to-hub path lies in the leave region; and (ii) Choice-update condition: the patch selected for the subsequent path differed from the current patch before the transition. (C) Patch evidence *Evi*_*p*_ (remaining patch resources) computed from the human observed patch activation matrices. Blue line with stars indicates the empirical choice outcome *C*^*emp*^(*t*) under the maximizing information acquisition strategy, which determines which patch to enter when the step falls in the leave region. The hub-to-hub paths at which the activated patch changes in (B) are identified as predicted transition steps, all other paths are identified as predicted non-transition steps, with two expectations: points b and d. The choice outcome *C*^*emp*^(*t*) of point d changes before the leave region emerged (light red box). (D) A special case that being identified as predicted transition step: at steps such as point b, resources of the current patch are exhausted within the trial, forcing the agent to leave even if the step lies in the stay region. (E) Differences between predicted transition steps and predicted non-transition steps in empirical choice update Δ*C*^tem^(*t*). The mean choice update Δ*C*^tem^(*t*) in predicted transition steps show values closer to 1, whereas in predicted non-transition steps show values closer to 0, yielding a significant separation at both the individual (left) and group (right) levels. *: p < 0.05, ***: p < 0.001.

Figure 2C illustrates the resulting correspondence for one representative participant. As the human narrative proceeded along the observed sequence of hub-to-hub paths, the predicted patch activation matrices showed patterns similar to the observed matrices (Figure 2C). To further characterize the positions of the human narrative and model prediction within the space spanned by all patches, we calculated a patch position center for each narrated path (see Methods). Both the observed and predicted position-center curves exhibited frequent changes across successive paths and showed similar trajectories (black and red curves in Figure 2C). Descriptively, cosine similarity between the observed and predicted matrices was near ceiling in all three trials of this sample (0.999, 1.000, and 1.000), and the circular distance between the position-center curves remained close to zero (0.030 ± 0.013, 0.000 ± 0.000, and 0.000 ± 0.000; Figure 2D, lower panel).

The matrix correspondence described above, however, did not isolate the patch-level decisions generated by the model’s foraging strategies, because a narrated path could automatically co-activate several overlapping patches regardless of the agent’s choice. We therefore separated the predicted activations into actively selected patches, reflecting the agent’s patch-entry, stay, and leave decisions under its foraging strategies; and co-activations induced automatically by patch overlap. The actively selected patches were then represented in active patch selection matrices (Figure 3B; formal criteria in Methods). Within this matrix, a transition step is a path boundary at which the model both left the current patch under its MVT-based rule and switched to a different patch for the next path. Figure 3B illustrates this classification for the example sample: at points a, c, and e, the leave rule was triggered while the patch still contained resources and the agent’s subsequent choice changed, so these boundaries were labeled predicted transition steps; point b was also counted, because the agent was forced to leave once the current patch had exhausted its resources for that trial; and point d was excluded, because the choice changed before the leave rule was triggered and thus did not reflect an MVT-generated transition. All remaining boundaries were labeled predicted non-transition steps.

Because the actively selected patch cannot be directly observed in human narratives, we next tested whether the model-predicted transition steps corresponded to analogous choice changes in the empirical data. For each narrated path, we derived empirical patch evidence from the remaining structure values updated according to the observed patch activation history (see Methods). The patch with the greatest remaining evidence was defined as the empirical choice outcome, *C*^emp^(*t*), which formed the choice trajectory shown by the blue line in Figure 3C. We then defined a choice-update indicator, Δ*C*^emp^(*t*), making the boundaries at which the favored patch changed from one path to the next (Figure 3C, red box). If human thought followed decision principles similar to those of the model agent, such choice updates should occur more frequently at model-predicted transition steps than at predicted non-transition steps. Within this illustrative sample, this was the case: the mean choice-update rate was higher at predicted transition steps than at predicted non-transition steps (0.750 ± 0.250 vs. 0.188 ± 0.101; t = 2.081, p = 0.038). Thus, the boundaries at which the model generated an active patch transition were also the boundaries at which the empirical patch evidence most often shifted to favor a different patch.

Together, these findings show that the MVT-based model captured two complementary aspects of human thought dynamics. Its predicted patch activation matrix reproduced which patches were associated with each narrated path, including co-activation caused by patch overlap. After these overlap-driven activations were removed, the remaining active patch transitions identified boundaries at which the human evidence was also more likely to favor a different patch. Human thought trajectories were therefore consistent with a combination of value-maximizing patch selection and an MVT-based balance between staying and leaving the current patch.

### Group-level patch dynamics showed alignment with model predictions

We next examined whether this correspondence generalized across the 97 participant–theme samples. Because the numbers of patches and narrated paths varied across samples, the matrices could not be averaged directly. For group-level visualization, we therefore projected the path and patch dimensions of every matrix onto a common normalized space, resampled the matrices to a fixed resolution, and ordered patches according to their temporal activation patterns in the observed matrix, applying the same row order to the corresponding predicted matrix (see Supplementary Methods S1). As shown in Figure 4A, the model-predicted matrices captured the principal temporal organization of the observed patch activation patterns in all three trials.

**Figure 4.**
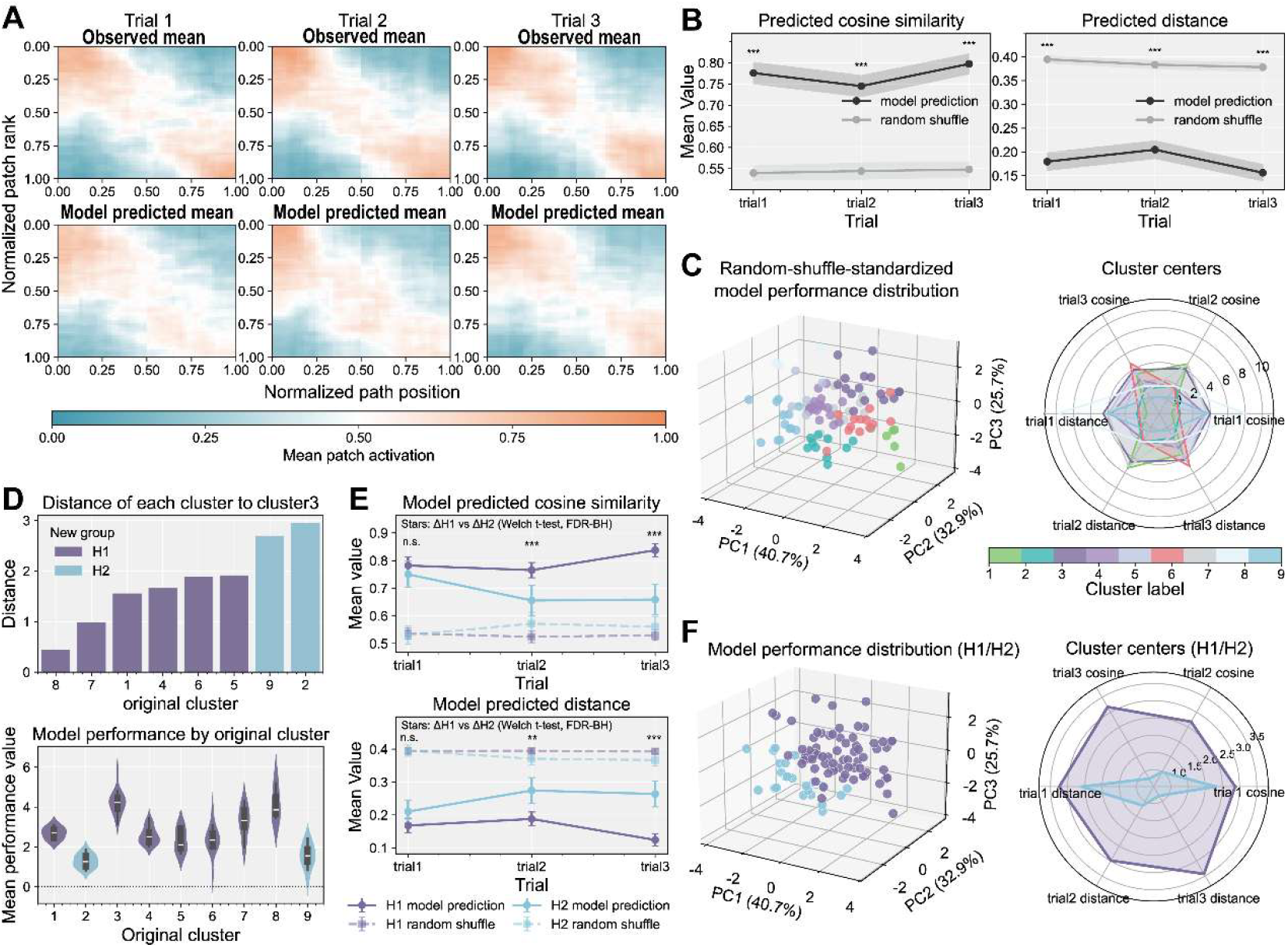
Group-level model performance and heterogeneous model conformity across participant–theme samples. (A) Group-averaged patch activation matrices for human narratives and model predictions. Top: the group-averaged observed patch activation matrices; bottom: the group-averaged predicted patch activation matrices, these matrices were resampled and projected onto a common normalized [0,1] × [0,1] space. Patches were reordered according to their temporal activation positions in the observed matrix, and the same row alignment was applied to the model-predicted matrix. The correspondence between the two matrices indicates that the model captured the principal temporal organization of human patch activation dynamics. (B) Group-level model performance. Cosine similarity and patch position distance were calculated for each participant–theme sample and then averaged across samples. Predictive performance was evaluated relative to a random shuffle baseline in which the temporal order of narrated paths was permuted, thereby removing information about which patches would be activated at the next path. The model significantly outperformed the random shuffle baseline in all three trials for both measures. Details of the random shuffle baseline are in Methods. (C) Clustering of human model-conformity profiles. Left: overall distribution of model performance across human participant–theme samples. Right: the nine clusters showed distinct trial-wise model-conformity profiles across trials 1–3, larger values and positions farther from the center consistently indicated better model performance. (D) Classification into the H1 and H2 groups. Cluster 3, which showed the strongest overall conformity with the model, was selected as the reference template. Distances between the remaining clusters and this template were then calculated. The template and clusters closer to it were combined into the H1 group (Clusters 1, 3, 4, 5, 6, 7, and 8), whereas the remaining clusters were combined into the H2 group (Clusters 2 and 9), representing samples with higher and lower model conformity, respectively. (E) Model performance in the H1 and H2 groups. Both groups performed significantly above the random shuffle baseline, but their predictive performance differed primarily in trials 2 and 3. (F) Distributions of model performance in the H1 and H2 groups. ***: p < 0.001.

Across all three trials, the cosine similarity of the MVT-based model was significantly higher than that of the random shuffle baseline. The mean ± SEM cosine similarities of the model were [0.776 ± 0.025, 0.745 ± 0.025, 0.797 ± 0.024] for trials 1–3, respectively, compared with [0.540 ± 0.017, 0.544 ± 0.019, 0.548 ± 0.018] for the random shuffle baseline (trial 1: t = 12.947, p < 0.001, Cohen’s d = 1.315; trial 2: t = 10.448, p < 0.001, Cohen’s d = 1.061; trial 3: t = 13.915, p < 0.001, Cohen’s d = 1.413; Figure 4B). Consistent with this result, the model-predicted patch position center trajectories were significantly closer to the observed trajectories than were those generated by the random shuffle baseline. The mean ± SEM distances of the model were [0.180 ± 0.018, 0.205 ± 0.019, 0.156 ± 0.017] across the three trials, whereas the corresponding random shuffle distances were [0.395 ± 0.008, 0.384 ± 0.008, 0.378 ± 0.009] (trial 1: t = −13.914, p < 0.001, Cohen’s d = −1.413; trial 2: t = −10.937, p < 0.001, Cohen’s d = −1.111; trial 3: t = −14.935, p < 0.001, Cohen’s d = −1.516; Figure 4B). These findings indicate that the model captured not only the overall distribution of patch activations but also their temporal organization along the narrated path sequence.

The model-predicted transition steps also corresponded to patch-level changes in the empirical human evidence at the group level. The mean choice-update rate was significantly higher at predicted transition steps than at predicted non-transition steps (mean ± SEM: transition, 0.397 ± 0.025; non-transition, 0.155 ± 0.010; t = 10.403, p < 0.001). Thus, the group-level results reproduced both components observed in the individual example: the model predicted which patches were activated along the narrative path sequence and identified transition boundaries at which the patch favored by the human evidence was more likely to change.

#### Robustness in an enriched memory environment

To examine whether this correspondence depended on using only the paths directly observed during the original narration sequence, we conducted an additional analysis based on the follow-up two-stage experiment. For the 25 participant-theme samples, we first constructed a Phase 1-only model using the originally observed paths. We then constructed a Phase 2 model that incorporated the additionally expanded paths identified from the Phase 2 narratives into the structural-value calculations. Both models reproduced the broad group-level pattern of the observed patch activation matrices (Supplementary Figure S1A). Relative to the random shuffle baseline, the Phase 1-only model showed significantly greater cosine similarity in all three trials (mean differences: 0.279, 0.218, and 0.284; t = 7.951, 7.463, and 8.363; all FDR-adjusted p < 0.001; Cohen’s d = 1.590, 1.493, and 1.673) and significantly smaller patch position distance (mean differences: −0.229, −0.183, and −0.228; t = −8.170, −6.986, and −8.076; all FDR-adjusted p < 0.001; Cohen’s d = −1.634, −1.397, and −1.615; Supplementary Figure S1B). The Phase 2 model also significantly outperformed the random shuffle baseline for both cosine similarity (differences: 0.155, 0.166, 0.213 across trials 1–3; t = 4.403, 4.119, and 6.074; all FDR-adjusted p < 0.001, Cohen’s d = 0.881, 0.824, and 1.215) and distance (differences: −0.123, −0.138, −0.184; t = −4.002, −4.120, and −6.117; all FDR-adjusted p < 0.001; Cohen’s d = −0.800, −0.824, and −1.223). Direct comparison of the two models showed that the Phase 1-only model performed better in trial 1 for cosine similarity (differences: 0.124, 0.051, 0.071; t = 3.007, 1.172, and t = 1.861; FDR-adjusted p = 0.008, 0.253, and 0.084; Cohen’s d = 0.601, 0.234, and 0.372), and distance (differences: −0.106, −0.045, and −0.044; t = −2.991, −1.288, and −1.616; FDR-adjusted p = 0.008, 0.210, and 0.134; Cohen’s d = −0.598, −0.258, and −0.323). No significant difference was found between the two models in trials 2 or 3. These results indicate that the model remained predictive when the foraging environment was enriched with additional memory resources associated with hub-related narrative content.

### 3.3 Group-level patch dynamics reveal heterogeneous conformity with optimal foraging strategies

Although the MVT-based model showed robust predictive performance at the group level, the degree of correspondence varied considerably across participant–theme samples. To characterize this heterogeneity, we represented each sample by a six-dimensional performance vector comprising the shuffle-standardized cosine similarity and position-center distance from the three trials, and applied k-means clustering to the 94 samples with complete values on all six dimensions (see Supplementary Methods S2 for parameter selection). This analysis identified nine performance clusters (Figure 4C), which we then collapsed into two groups based on their similarity to the best-performing cluster. Specifically, taking Cluster 3 — the cluster showing the strongest overall conformity with model predictions — as the reference template, we computed the Wasserstein distance between the joint performance distribution of each remaining cluster and that of the template. The two most distant clusters (Clusters 2 and 9) were combined into a low-conformity group (H2), and the remaining seven clusters into a high-conformity group (H1) (Figure 4D, E). Of the 94 samples, 71 (75.5%) fell into H1 and 23 (24.5%) into H2, indicating that the model captured the narrative dynamics of the majority of samples, while a substantial minority systematically deviated from its predictions.

H1 showed higher model predicted cosine similarity and lower patch position distance than H2, particularly in trials 2 and 3. For cosine similarity (mean ± SEM, trials 1–3; H1: 0.782 ± 0.031, 0.765 ± 0.028, 0.838 ± 0.024; H2: 0.750 ± 0.047, 0.656 ± 0.055, 0.659 ± 0.056), the group difference was nonsignificant in trial 1 (t = 0.589, FDR-adjusted p = 0.575, Cohen’s d = 0.142) but significant in trials 2 and 3 (t = 4.323 and 6.036, both FDR-adjusted p < 0.001; Cohen’s d = 0.878 and 1.395). The same pattern held for position-center distance (H1: 0.168 ± 0.022, 0.188 ± 0.021, 0.124 ± 0.017; H2: 0.210 ± 0.035, 0.274 ± 0.039, 0.264 ± 0.039; trial 1: t = 1.188, FDR-adjusted p = 0.276, d = 0.270; trial 2: t = 3.279, p = 0.004, d = 0.711; trial 3: t = 5.392, p < 0.001, d = 1.309).

Importantly, both groups nevertheless outperformed the random-shuffle baseline on both measures in all three trials. For H1, cosine similarity exceeded the baseline (t = 11.429, 10.653, and 16.987; all FDR-adjusted p < 0.001; Cohen’s d = 1.356, 1.264, and 2.016) and position-center distance fell below it (t = −12.223, −10.768, and −18.003; all p < 0.001; d = −1.451, −1.278, and −2.137). For H2, these advantages were smaller but remained significant (cosine similarity: t = 5.754, 2.931, and 3.322; FDR-adjusted p = 0.001, 0.010, and 0.005; Cohen’s d = 1.200, 0.611, and 0.693; distance: t = −6.326, −3.475, and −3.839; p < 0.001, 0.004, and 0.002; d = −1.319, −0.725, and −0.801).

Together, these findings indicate that human free narratives exhibited heterogeneous levels of correspondence with the MVT-based model. H1 showed stronger and more stable model conformity, particularly across later trials, whereas H2 showed comparatively weaker correspondence. Nevertheless, the performance of both groups remained significantly better than the random shuffle baseline, suggesting that foraging-like patch dynamics were present across the classified human samples but varied substantially in strength.

### 3.4 Patch transitions mark semantic boundaries and vary with theme-related evaluations

#### Semantic characteristics of model-predicted patch transitions

We next examined whether the patch transition steps predicted by the MVT-based model corresponded to systematic changes in the semantic organization of human narratives, using four complementary analyses of the semantic relationships among hub nodes and paths, both within patches and around predicted patch transitions (Figure 5B).

**Figure 5.**
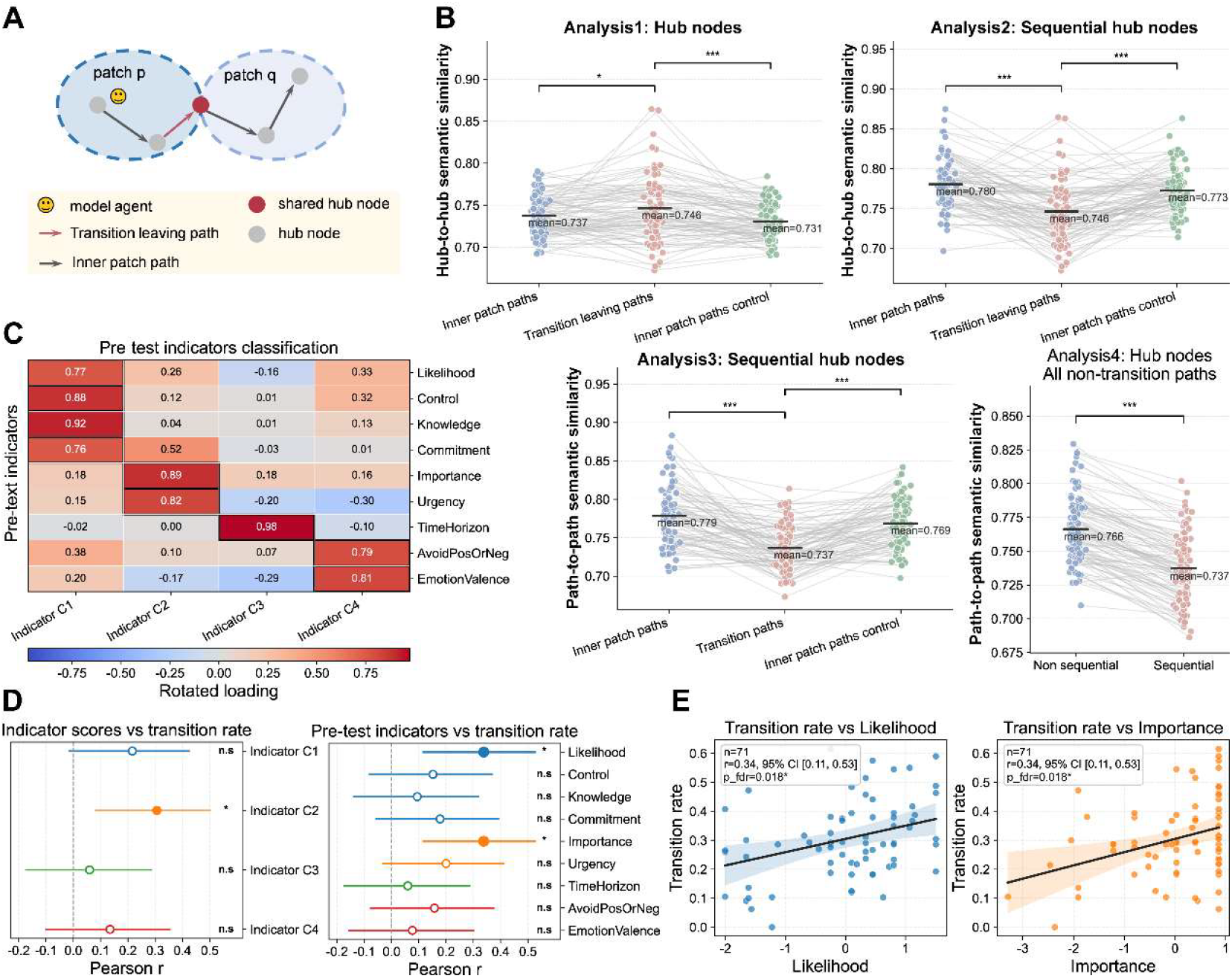
Model-predicted patch transitions marked semantic boundaries and varied with event importance and likelihood. (A) Illustration of inner-patch paths and transition-leaving paths. Transition-leaving paths were defined as the paths immediately preceding model-predicted transitions between actively selected patches. Inner-patch paths comprised paths located within patches after transition-leaving paths had been excluded. (B) Results of the four transition-related semantic analyses. In Analysis 1, transition-leaving paths showed greater semantic continuity than inner-patch paths when temporally connected information between hub nodes was omitted. In Analyses 2 and 3, transition-related paths showed greater semantic change when temporal connected information was retained, at the hub-node and path levels, respectively. In Analysis 4, the cross-semantic similarity between sequentially connected paths was lower than that between non-sequential paths, indicating additional semantic reorganization at the next-path selection stage. (C) Factor structure of the nine pre-experiment theme-rating indicators. The indicators were grouped into four factors, denoted C1–C4. (D) Associations between theme evaluations and model-predicted patch transition rate. Left: correlations between the four indicator factors and transition rate. C2 was significantly positively correlated with transition rate. Right: correlations between individual indicators and transition rate. Likelihood and Importance were positively correlated with transition rate, although the C1 factor containing Likelihood was not significant. (E) Scatterplots showing the associations of Likelihood and Importance with model-predicted patch transition rate. *: p < 0.05.

Analysis 1 asked whether patches and transition boundaries differed in overall semantic coverage, irrespective of the temporal order in which hub nodes were visited. We distinguished transition-leaving paths from inner-patch paths. Here, transition-leaving paths referred to the paths immediately preceding model-predicted patch transitions, whereas inner-patch paths referred to paths contained within patches after transition-leaving paths had been excluded (Figure 5A). Because patch overlap could contribute the same path to multiple patches, we additionally constructed an inner-patch control set by removing duplicate values caused by patch overlap from the inner-patch paths. Using the inner-patch paths, semantic similarity at transition-leaving paths exceeded that within patches by 0.009 (t = 2.600, FDR-adjusted p = 0.016, Cohen’s d = 0.271). When duplicate paths caused by patch overlap were removed (i.e. the inner-patch paths control), the mean difference increased to 0.016 (t = 4.798, FDR-adjusted p < 0.001, Cohen’s d = 0.500; Analysis 1). Thus, when temporal adjacency was disregarded, the hub nodes sampled within patches covered a broader semantic range than those occurring along the paths immediately preceding predicted patch transitions.

A different pattern emerged when the temporal order between hub nodes were taken into account (Analysis 2). The semantic similarity between sequentially connected hub nodes was significantly lower at transition-leaving paths than at inner-patch paths, both for the full path set (mean difference = −0.034; t = −7.907, FDR-adjusted p < 0.001, Cohen’s d = −0.824) and for the deduplicated control set (mean difference = −0.026; t = −6.510, FDR-adjusted p < 0.001, Cohen’s d = −0.679). These results indicate that, despite the greater overall semantic diversity of the hub nodes represented within a patch, sequentially connected hub nodes showed greater semantic continuity within patches; by contrast, the paths at which the model agent left the current patch were associated with larger local semantic changes. The same pattern was observed at the path level (Analysis 3). Consecutive path pairs spanning a model-predicted patch transition showed lower cross-path semantic similarity than consecutive path pairs for which the model agent stayed in the same patch, again both for the full path set (mean difference = −0.042; t = −9.380, FDR-adjusted p < 0.001, Cohen’s d = −0.989) and for the deduplicated control set (mean difference = −0.032; t = −8.615, FDR-adjusted p < 0.001, Cohen’s d = −0.908). Leaving the current patch and entering another was therefore accompanied by a more pronounced semantic change between consecutively narrated paths. Together, these findings are consistent with the possibility that the patch transition steps identified by the MVT-based model capture spontaneously emerging semantic segmentation boundaries during free narration within a personally relevant theme.

Moreover, the MVT-based model predicted patch-level transition behavior rather than the identity of the specific path narrated next. We therefore asked whether the observed next-path choices themselves exhibited a systematic semantic pattern (Analysis 4). Sequential path pairs showed significantly lower semantic similarity than non-sequential path pairs from the same narrative trial (mean difference = −0.029, t = −19.389, FDR-adjusted p < 0.001, Cohen’s d = −1.969). This finding suggests that next-path selection was not semantically arbitrary but was biased toward comparatively dissimilar semantic content.

The Phase 2 model largely reproduced these above semantic patterns. Analyses 2–4 remained significant with effect directions consistent with the primary results (Supplementary Figure S1C and Table 4).

### Theme-related variation in patch transition frequency

Having established the semantic characteristics of individual patch transitions, we next asked whether the frequency of these transitions varied with the characteristics of the narrated themes. This analysis was restricted to the group H1, whose patch dynamics showed stronger conformity with the MVT-based model. For each sample, we related the patch transition rate to nine ratings collected before the narration task, proceeding at three levels of resolution. First, because the nine ratings were substantially intercorrelated, we tested associations at the level of latent dimensions derived from principal component analysis (PCA), which pools correlated ratings into orthogonal factors. Second, to localize significant factor-level associations to specific theme attributes, we tested each rating individually. Finally, for ratings loading on the same factor, we used simultaneous regression to determine whether their associations reflected variance shared within the factor or information specific to each attribute.

The rating data were suitable for dimensionality reduction, as indicated by an overall Kaiser-Meyer-Olkin value of 0.784 and a significant Bartlett’s test of sphericity (*χ*^2^ (36) = 320.027, p < 0.001). PCA followed by varimax rotation identified four interpretable factors, *C1-C4*, jointly explaining 83.86% of the total variance (Figure 5C). C1 (44.28% of variance) included Knowledge (rotated loading: 0.916), Control (0.879), Likelihood (0.765), and Commitment (0.756), reflecting expectations and preparedness concerning the event’s realization. C2 (19.94%) included Importance (0.888) and Urgency (0.822), reflecting the motivational strength to act on the event. C3 (11.99%) was dominated by Time Horizon (loading = 0.975), representing the event’s expected temporal range. C4 (7.66%) included Emotion Valence (0.811) and Avoidance Orientation (0.793), reflecting emotional evaluations of the event.

Among the four factors, only *C2* was significantly associated with patch transition rate (r = 0.306, p = 0.009, FDR-adjusted p = 0.038; Figure 5D), which suggests that narratives concerning themes with greater motivational strength, jointly indexed by Importance and Urgency, tended to contain more frequent patch transitions. *C1* showed a positive but non-significant association with transition rate (r = 0.216, p = 0.070, FDR-adjusted p = 0.140). Neither *C3* (r = 0.060, p = 0.622, FDR-adjusted p = 0.622) nor *C4* (r = 0.134, p = 0.264, FDR-adjusted p = 0.351) was significantly related to transition rate.

To identify the specific theme attributes underlying these factor-level associations, we next tested each of the nine ratings separately. Likelihood was positively correlated with patch transition rate (r = 0.338, p = 0.0039, FDR-adjusted p = 0.0179), as was Importance (r = 0.338, p = 0.0040, FDR-adjusted p = 0.0179), as shown in Figure5E. None of the remaining ratings were significant under FDR correction (Control, r = 0.152, FDR-adjusted p = 0.309; Knowledge, r = 0.094, FDR-adjusted p = 0.559; Commitment, r = 0.178, FDR-adjusted p = 0.309; Urgency, r = 0.200, FDR-adjusted p = 0.284; TimeHorizon, r = 0.060, FDR-adjusted p = 0.622; AvoidPosOrNeg, r = 0.157, FDR-adjusted p = 0.309; and EmotionValence, r = 0.076, FDR-adjusted p = 0.592). The association observed for Factor *C2* therefore appeared to be driven primarily by Importance. Although Factor *C1* as a whole was not significantly related to transition rate, Likelihood showed a reliable item-level association.

Because the four ratings comprising *C1* were substantially correlated with one another (Supplementary Table 2, 3), the association between Likelihood and transition rate could reflect either variance shared with Control, Knowledge, and Commitment or information specific to Likelihood itself. To distinguish these possibilities, we fitted a linear mixed-effects model with participant as a random intercept and the standardized Likelihood, Control, Knowledge, and Commitment scores entered simultaneously as fixed effects. After controlling for the other Factor *C1* ratings, Likelihood remained a significant positive predictor of transition rate (*β* = 0.582, SE = 0.202, z = 2.874, p = 0.004). In contrast, Control (*β* = −0.239, p = 0.297), Knowledge (*β* = −0.139, p = 0.472), and Commitment (*β* = 0.055, p = 0.759) were not significant predictors. A likelihood-ratio test further showed that including Likelihood significantly improved model fit relative to a model from which it was removed, *χ*^2^ (1) = 9.372, p = 0.002. Thus, the relationship between patch transition rate and Likelihood could not be explained solely by the evaluations shared across the *C1* ratings.

Taken together, the semantic and theme-rating analyses characterize two complementary aspects of patch transitions. At the local level, model-predicted transitions marked boundaries associated with greater semantic change between consecutively narrated contents. At the theme level, these boundaries occurred more frequently for events that were considered more important or more likely to be realized. Patch transitions therefore appeared to reflect a theme-dependent process of segmenting and reorganizing event-related thought rather than arbitrary switches between narrative contents. Such segmentation and reorganization may provide an organizational basis for internal event planning.

### 3.5 LLM-generated narratives show weaker but detectable conformity with foraging-like patch dynamics

#### LLM narratives showed detectable but reduced model conformity

To examine whether foraging-like thought dynamics also emerged in LLM-generated narratives, we used Qwen-max (Qwen-max-2025-01-25) and Qwen-flash (Qwen-flash-2025-07-28), both of which support Chinese-language generation. The two models received the same task instructions used in the human free-narration experiment. Each participant’s self-reported themes were provided as fixed topics, and the LLMs were instructed to freely narrate within the scope of each theme. The generated texts were processed using the same pipeline applied to the human narratives. For each eligible LLM-generated sample in which a patch network could be constructed, we applied the same MVT-based simulation and evaluated how closely the predicted patch activation and transition dynamics matched those derived from the generated narrative.

At the group level, the predicted matrices for Qwen-max captured the broad temporal organization of the corresponding observed patch activation matrices, although the latter part of trial 2 showed weaker correspondence and approached the random-shuffle pattern. The predicted matrices for Qwen-flash showed less consistent correspondence overall. Its clearest match to the observed activation pattern occurred in trial 3, particularly during the earlier and later portions of the narrative sequence (Figure 6A).

**Figure 6.**
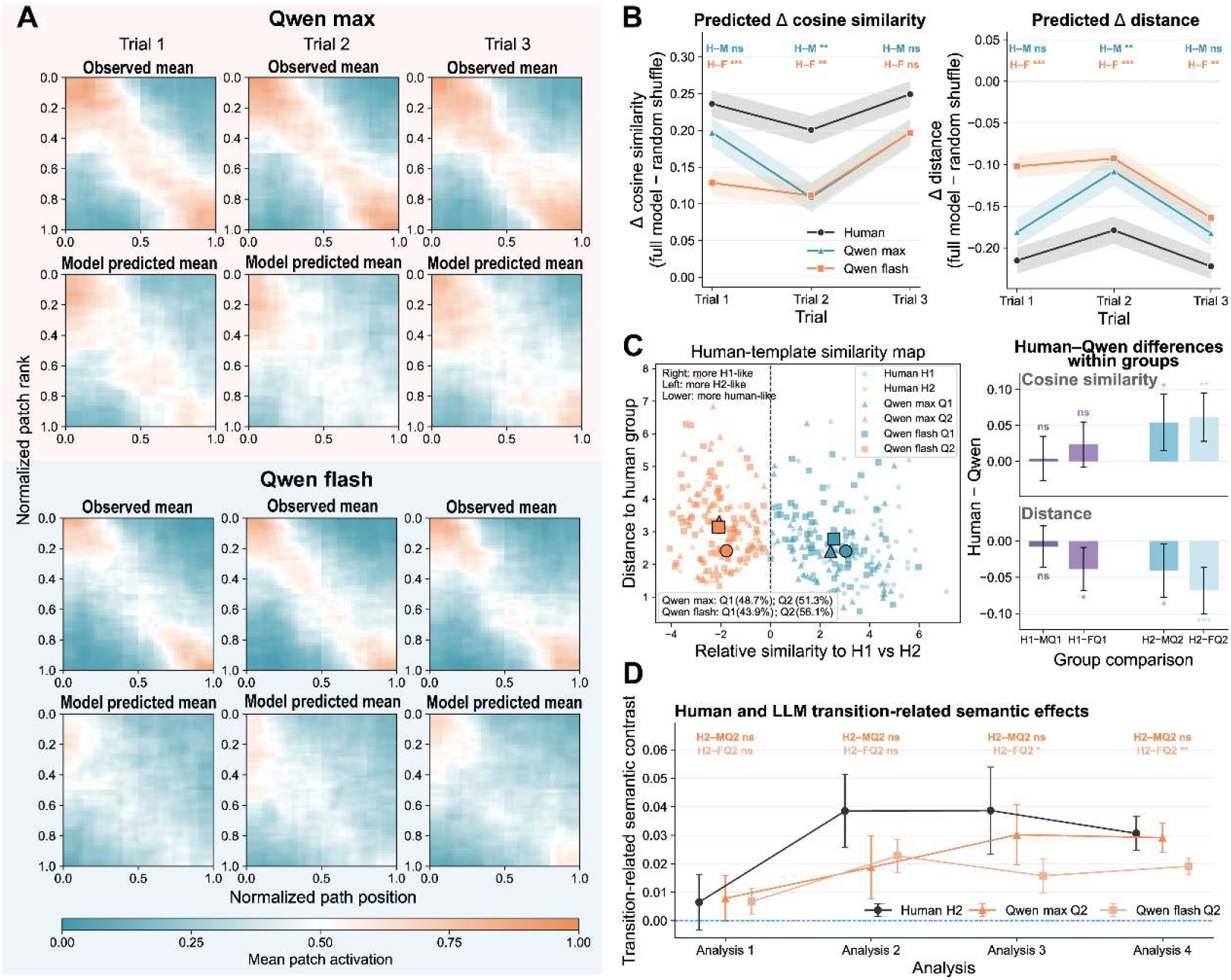
Human and Qwen narratives showed shared but heterogeneous conformity with the MVT-based model. (A) Normalized and resampled patch activation matrices for Qwen-max samples (top) and Qwen-flash samples (bottom). The Qwen-max predictions generally showed closer correspondence with the observed matrices, whereas Qwen-flash showed weaker correspondence in the earlier trials and clearer alignment in trial 3. (B) Comparison of model performance across human, Qwen-max, and Qwen-flash narratives. Performance was expressed relative to the random shuffle baseline using Δcosine similarity (left) and Δdistance (right). Qwen-max generally showed model performance closer to the human samples than Qwen-flash, although this pattern varied across trials. (C) Classification of Qwen samples according to the human model-conformity structure. Left: Qwen samples were assigned to Q1 when they were closer to the human H1 centroid and to Q2 when they were closer to the human H2 centroid. Right: comparison of performance among the corresponding human and Qwen groups. Q1 samples more closely approximated the H1 group, whereas Q2 samples showed lower model conformity than the human H2 group. (D) Comparison of transition-related semantic effect magnitudes across human and Qwen narratives. For Analysis 1, the effect was defined as the semantic measure for transition-leaving paths minus that for inner-patch paths. For Analyses 2 and 3, the effect was defined as the measure for inner-patch paths minus that for transition-leaving paths. For Analysis 4, the effect was defined as the measure for non-sequential paths minus that for sequential paths. *: p < 0.05, **: p < 0.01, ***: p < 0.001.

Model performance was expressed relative to the random shuffle baseline, with positive Δcosine similarity and negative Δdistance indicating better performance. For Qwen-max, Δcosine similarity was significantly greater than zero in all three trials (mean ± SEM = 0.197 ± 0.019, 0.109 ± 0.020, and 0.197 ± 0.019; t = 10.331, 5.390, and 10.354; all FDR-adjusted p < 0.001; Cohen’s d = 0.935, 0.488, and 0.937; Figure 6B), and Δdistance was significantly below zero (mean ± SEM = −0.181 ± 0.017, −0.108 ± 0.017, and −0.182 ± 0.015; t = −10.817, −6.260, and −11.778; all FDR-adjusted p < 0.001; Cohen’s d = −0.979, −0.567, and −1.066). Thus, the MVT-based model captured non-random temporal structure in the Qwen-max narratives across all three trials. The performance of Qwen-max was generally close to that observed in the human samples, the difference was significant only in trial 2 after FDR correction (trial 1: t = 1.475, FDR-adjusted p = 0.142, Cohen’s d = 0.197; trial 2: t = 3.297, FDR-adjusted p = 0.002, Cohen’s d = 0.440; trial 3: t = 1.987, FDR-adjusted p = 0.058, Cohen’s d = 0.265). Similarly, the differences between human and Qwen max in Δdistance survived FDR correction only in trial 2 (trial 1: t = −1.488, FDR-adjusted p = 0.138, Cohen’s d = −0.198; trial 2: t = −2.982, FDR-adjusted p = 0.006, Cohen’s d = −0.398; trial 3: t = −1.855, FDR-adjusted p = 0.078, Cohen’s d = −0.248).

The Qwen-flash narratives also showed performance significantly above the random shuffle baseline, although their correspondence with the model was weaker than that of the human narratives. Δcosine similarity was significantly greater than zero in all three trials (mean ± SEM = 0.129 ± 0.017, 0.111 ± 0.017, and 0.197 ± 0.017; t = 7.537, 6.467, and 11.692; all FDR-adjusted p < 0.001; Cohen’s d = 0.635, 0.545, and 0.985), and Δdistance was significantly below zero (mean ± SEM = −0.102 ± 0.014, −0.093 ± 0.014, and −0.164 ± 0.014; t = −7.470, −6.863, and −11.780; all FDR-adjusted p < 0.001; Cohen’s d = −0.629, −0.578, and −0.992). Human samples showed greater Δcosine similarity than Qwen-flash, these differences were significant in trials 1 and 2, but the trial 3 difference did not survive FDR correction (trial 1: t = 4.310, FDR-adjusted p < 0.001, Cohen’s d = 0.556; trial 2: t = 3.465, FDR-adjusted p = 0.002, Cohen’s d = 0.451; trial 3: t = 2.131, FDR-adjusted p = 0.051, Cohen’s d = 0.275). Human samples also showed more negative Δdistance than Qwen-flash in all three trials (trial 1: t = −5.484, FDR-adjusted p < 0.001, Cohen’s d = −0.715; trial 2: t = −4.063, FDR-adjusted p < 0.001, Cohen’s d = −0.536; trial 3: t = −2.861, FDR-adjusted p = 0.007, Cohen’s d = −0.369).

Direct comparison of the two LLMs showed that Qwen-max outperformed Qwen-flash primarily in trial 1 (Δcosine similarity, t = 2.683, FDR-adjusted p = 0.023, Cohen’s d = 0.333; Δdistance, t = −3.665, FDR-adjusted p < 0.001, Cohen’s d = −0.457), whereas no difference was detected in trial 2 or trial 3 (Δcosine similarity, FDR-adjusted p = 0.987 for both; Δdistance, FDR-adjusted p = 0.485). These results indicate that the relatively weak activation-pattern correspondence of Qwen-flash was most pronounced during the first trial.

#### LLM samples showed a more heterogeneous distribution of model conformity

We next examined how the LLM-generated samples were distributed relative to the heterogeneous human performance profiles. Each Qwen sample was assigned to the nearest of the nine original human clusters according to its Euclidean distance from the corresponding cluster centroids in the six-dimensional performance space (Figure 6C, left). A Qwen sample assigned to a human cluster belonging to H1 was classified as Q1, whereas a sample assigned to an H2 cluster was classified as Q2. Among the 117 eligible Qwen-max samples, 57 were assigned to Q1 (48.7%) and 60 to Q2 (51.3%). Among the 139 eligible Qwen-flash samples, 61 were assigned to Q1 (43.9%) and 78 to Q2 (56.1%). Thus, approximately half of the Qwen-max samples and slightly fewer than half of the Qwen-flash samples occupied regions of the performance space associated with the more strongly model-conforming human group.

The performance of the Q1 groups was generally similar to that of human H1 (Figure 6C, right). Human H1 and Qwen-max Q1 did not differ in either cosine similarity (mean difference = 0.004, t = 0.242, FDR-adjusted p = 0.809, Cohen’s d = 0.043) or distance (mean difference = −0.008, t = −0.536, FDR-adjusted p = 0.593, Cohen’s d = −0.094). Human H1 and Qwen-flash Q1 also did not differ significantly in cosine similarity (t = 1.492, FDR-adjusted p = 0.184, Cohen’s d = 0.262), although Qwen-flash Q1 showed a significantly greater distance (t = −2.567, FDR-adjusted p = 0.023, Cohen’s d = −0.449). In contrast, both Q2 groups performed significantly worse than human H2. Human H2 showed higher cosine similarity than Qwen-max Q2 (t = 2.737, FDR-adjusted p = 0.016, Cohen’s d = 0.524) and Qwen-flash Q2 (t = 3.654, FDR-adjusted p = 0.002, Cohen’s d = 0.700). Human H2 also showed lower distance than Qwen-max Q2 (t = −2.199, FDR-adjusted p = 0.042, Cohen’s d = −0.437) and Qwen-flash Q2 (t = −4.243, FDR-adjusted p < 0.001, Cohen’s d = −0.869). These results indicate that LLM-generated samples broadly followed the human pattern of occupying both more and less model-conforming regions of the performance space. However, the less model-conforming LLM samples showed weaker correspondence with the MVT-based model than the human H2 group.

We further examined whether patch transitions in the LLM-generated narratives were accompanied by the semantic changes observed in human narration. Both Qwen-max and Qwen-flash largely reproduced the human-like semantic patterns: Analyses 1–4 were significant and showed effect directions consistent with the human results (Supplementary Figure S2; Table 5, 6). Thus, LLM-predicted patch transitions were also associated with local semantic variation, and sequential path selection showed the same bias toward comparatively dissimilar content.

We further compared the magnitude of these semantic effects among the less model-conforming groups. Effect magnitudes were the differences in semantic change between transition-related paths and inner-patch paths in Analyses 1–3, and between non-sequential and sequential paths in Analysis 4 (Figure 6D). No comparison between human H2 and Qwen-max Q2 survived FDR correction across the four analyses (FDR-adjusted p = 0.954, 0.084, 0.411, and 0.824, respectively). By contrast, Qwen-flash Q2 showed weaker effects than human H2 in Analyses 3 and 4 (Analysis 3, t = 2.768, FDR-adjusted p = 0.045, Cohen’s d = 0.730; Analysis 4, t = 3.334, FDR-adjusted p = 0.002, Cohen’s d = 0.859), the differences in Analyses 1 and 2 were not significant after FDR correction. Qwen-max Q2 also showed a larger Analysis 4 effect than Qwen-flash Q2 (t = 3.324, FDR-adjusted p < 0.001, Cohen’s d = 0.646), their difference in Analysis 3 did not survive FDR correction (FDR-adjusted p = 0.054).

Taken together, both LLMs exhibited non-random, foraging-like patch dynamics and transition-related semantic segmentation. However, their overall correspondence with the MVT-based model was weaker and more heterogeneous than that observed in human narratives. Qwen-max showed the more human-like profile, particularly in its patch activation dynamics and in the magnitude of the semantic changes associated with patch transitions. Qwen-flash retained the general direction of the human semantic effects but showed weaker transition-related semantic differentiation among its Q2 samples.

### 3.6 Foraging decisions and multilevel structure values jointly supported model prediction

We next examined whether the patch activation patterns predicted by the MVT-based model could be explained by the topology of the patch network alone or by simpler forms of foraging-related control. We compared the full model mainly with three types of control models: a patch semantic random walk model, which omitted foraging decisions and value evaluation, while used the semantic similarity between patches to determine patch-to-patch transition probabilities; two structure-value control models, the No S and No L models, which retained the foraging rules but removed the third-order or second-order structure value component, respectively; and a Depletion history model in which patch values depended only on their cumulative activation histories. All models predicted patch activation matrices and were evaluated using the same cosine-similarity and patch-position-distance measures.

#### Control model performance in human narratives

In the human narratives, the patch semantic random walk reproduced part of the observed patch activation pattern, particularly during the earlier portion of each trial. Its predictions became less differentiated later in the path sequence and approached the random shuffle pattern (Figure 7A). Nevertheless, the random walk performed significantly better than random shuffle baseline in all three trials for Δcosine similarity and Δdistance (all FDR-adjusted p < 0.001). However, the random walk performed substantially worse than the full model in all three trials, with lower cosine similarity (t = 8.751, 6.099, and 8.941; all FDR-adjusted p < 0.001; Cohen’s d = 0.889, 0.619, and 0.908), and higher patch position distance (t = −9.422, −6.276, and −10.172; all FDR-adjusted p < 0.001; Cohen’s d = −0.957, −0.637, and −1.033; Figure 7B). We also tested alternative patch-to-patch transition rules based on patch overlap, patch size, or uniform transition probabilities rather than semantic similarity. These variants performed similarly to the patch semantic random walk used in the main analysis (Supplementary Figure S3; Supplementary Tables 7 and 8). Thus, patch-to-patch relationships alone, regardless of how transition probabilities were defined, were insufficient to reproduce the full temporal organization of human patch activation.

**Figure 7.**
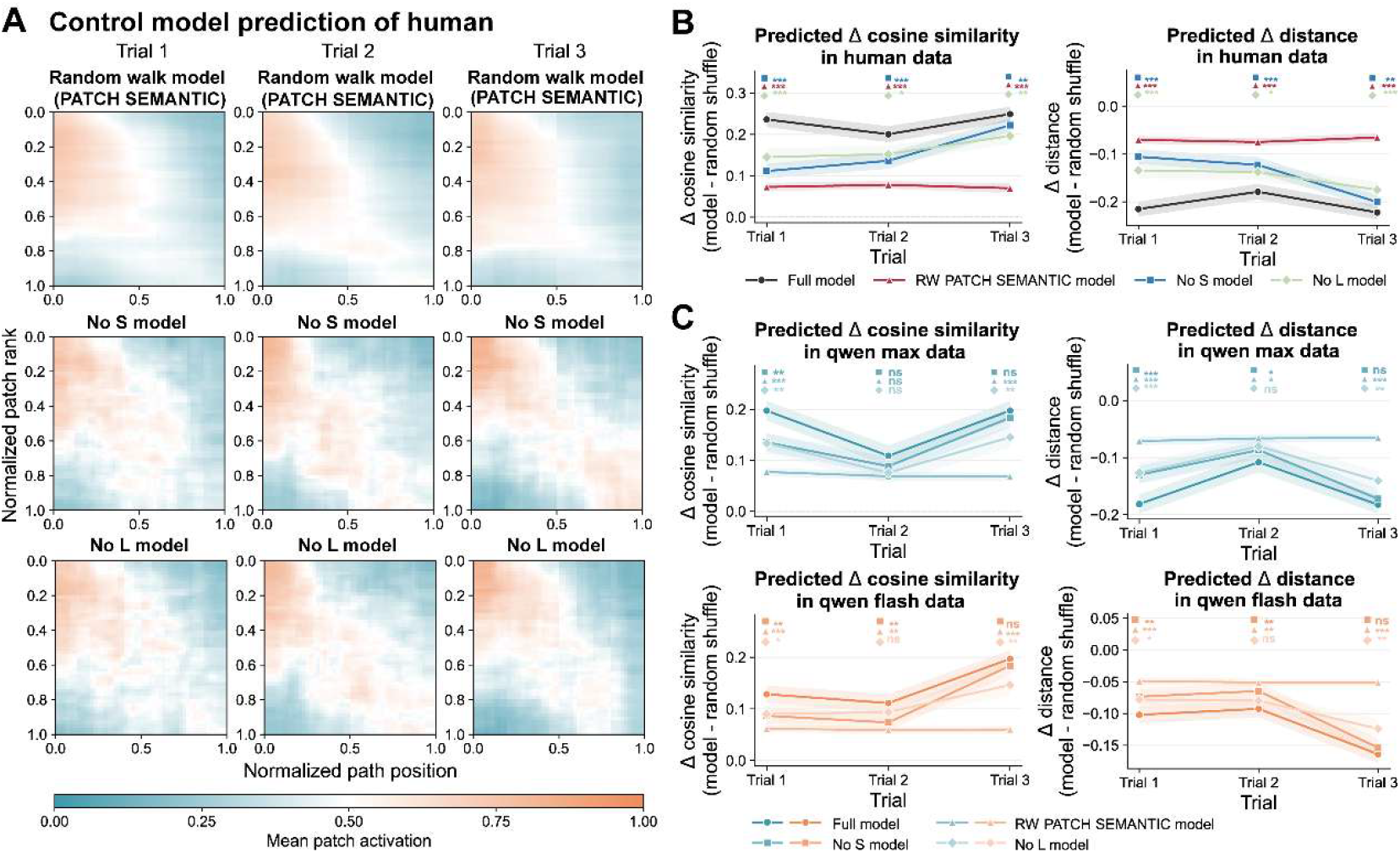
Control-model analyses of foraging decisions and multilevel structure values. (A) Normalized and resampled patch activation matrices predicted by the three principal control models. The No S and No L models retained the foraging-related decision rules while preserving only one order of structure-value evaluation. Their predictions showed closer correspondence with the observed human activation dynamics than those of the patch semantic random walk. By contrast, the random-walk predictions became less differentiated during the later portions of the path sequences in all three trials. (B) Comparison of control-model and full-model performance for human narratives. Although all three control models performed above the random shuffle baseline, each performed significantly worse than the full model in both cosine similarity (left) and patch position distance (right). The No S and No L models also generally outperformed the random walk, indicating that foraging-related decisions combined with either order of structure value provided additional predictive information. (C) Comparison of control-model and full-model performance for Qwen narratives. Top: for Qwen-max, differences between the control models and the full model were smaller and varied across trials, particularly in trials 2 and 3. Bottom: for Qwen-flash, the No S and No L models also approached the performance of the full model in some later trials, whereas the random walk showed weaker performance. *: p < 0.05, **: p < 0.01, ***: p < 0.001.

We next examined the two structure-value control models. The No S model retained the second-order structure value and foraging-related decision rules but removed the third-order structure value. Conversely, the No L model retained the third-order structure value and the same patch entry and leaving rules but removed the second-order structure value. Both models reproduced the broad organization of the observed activation matrices, particularly in trials 2 and 3, although their predictions became less differentiated during the later portion of trial 1 (Figure 7A). Both models performed significantly above the random shuffle baseline in all three trials: for the No S model, Δcosine similarity was higher than the shuffle baseline (t = 6.467, 7.119, and 13.338; Cohen’s d = 0.657, 0.723, and 1.354), and Δdistance was lower (t = −7.094, −7.490, and −14.019; Cohen’s d = −0.720, −0.760, and −1.423); The No L model showed the same pattern for Δcosine similarity (t = 6.956, 7.690, and 9.890; Cohen’s d = 0.706, 0.781, and 1.004), and Δdistance (t = −7.530, −8.186, and −11.059; Cohen’s d = −0.765, −0.831, and −1.123). All FDR-adjusted p values were < 0.001.

Despite this predictive ability, both structure-value controls performed significantly worse than the full model in every trial for both cosine similarity (for the No S model, t = 6.854, 4.660, and 3.213; FDR-adjusted p < 0.001, < 0.001, and 0.003; Cohen’s d = 0.696, 0.473, and 0.326. For the No L model, t = 4.849, 2.491, and 3.232; FDR-adjusted p < 0.001, 0.020, and 0.003; Cohen’s d = 0.492, 0.253, and 0.328), and patch position distance (for the No S model, t = −6.855, −4.787, and −3.396; FDR-adjusted p < 0.001, < 0.001, and 0.002; Cohen’s d = −0.696, −0.486, and −0.345. For the No L model, t = −4.838, −2.327, and −3.612; FDR-adjusted p < 0.001, 0.028, and < 0.001; Cohen’s d = −0.491, −0.236, and −0.367). Thus, either structure-value component combined with the foraging rules provided partial predictive information, but neither reproduced the performance of the full model.

In comparisons among the control models, both the No S and No L models outperformed the random walk in all three human trials for both cosine similarity and distance (for the No S model, all FDR-adjusted p ≤ 0.020; for the No L model, all FDR-adjusted p ≤ 0.001). No consistent performance difference was observed between the No S and No L models. The No L model showed a small cosine-similarity advantage in trial 1, whereas the corresponding distance difference did not survive FDR correction, and neither measure differed significantly in trials 2 or 3. These results indicate that, compared with patch-to-patch relationships alone, the foraging-related decision process provided greater predictive information about human narrative dynamics even when only one order of structure value was available.

We further tested whether the structure-value terms could be replaced by a simpler history-dependent depletion mechanism. In the Depletion history model, patch values were determined only by cumulative activations across trials. This model performed significantly worse than the full model in all three trials for both cosine similarity (t = 10.641, 7.650, and 7.030; all FDR-adjusted p < 0.001; Cohen’s d = 1.080, 0.777, and 0.714), and patch position distance (t = −11.190, −7.695, and −7.477; all FDR-adjusted p < 0.001; Cohen’s d = −1.136, −0.781, and −0.759), see Supplementary Figure S3 and Table 8. Accumulated patch activation history alone was therefore insufficient to account for the predictive performance of the full model.

Overall, the human control analyses revealed a consistent hierarchy of predictive performance. The No S and No L models more closely approximated the full model, indicating that either the second-order or third-order structure value could support partial prediction when combined with foraging decisions. By contrast, the patch semantic random walk performed substantially worse than models that retained the foraging-related decision rules. The Depletion history model retained a basic foraging process but discarded the evaluation of structure values was also inferior to the full model. These results support the joint contribution of foraging-related decisions and multilevel structure values to the prediction of human patch activation dynamics.

#### Control-model performance in LLM-generated narratives

We conducted the same control analyses for the Qwen-max and Qwen-flash narratives (Figure 7C). For both LLMs, all four control models performed significantly above the random shuffle baseline in each trial (details of statistical results in Supplementary Table 9, 11). These results indicate that patch relationships, individual structure-value components, and activation history each captured some non-random organization in the LLM-generated narratives.

For Qwen-max, the full model generally outperformed all control models, although some controls differed only slightly from the full model in trials 2 and 3 (upper panel of Figure 7C, and Supplementary Figure S4). In trial 2, all control models except the Depletion history model showed cosine similarity that did not differ significantly from that of the full model (No L model: FDR-adjusted p = 0.079, Cohen’s d = 0.178; No S model: FDR-adjusted p = 0.130, Cohen’s d = 0.153; random walk model: FDR-adjusted p = 0.068, Cohen’s d = 0.187; Depletion history model: FDR-adjusted p = 0.002, Cohen’s d = 0.293); differences in distance were also relatively small in this trial (No L model: FDR-adjusted p = 0.070, Cohen’s d = −0.178; No S model: FDR-adjusted p = 0.037, Cohen’s d = −0.207; random walk model: FDR-adjusted p = 0.019, Cohen’s d = −0.234; Depletion history model: FDR-adjusted p < 0.001, Cohen’s d = −0.332). The No S model showed the closest overall performance to the full model and also did not differ significantly from it in trial 3 (cosine similarity: FDR-adjusted p = 0.180, Cohen’s d = 0.135; distance: FDR-adjusted p = 0.225, Cohen’s d = −0.120).

In comparisons among the control models of Qwen-max, both structure-value controls generally outperformed the random walk in trials 1 and 3 but not trial 2. For the No S model, this pattern was observed for cosine similarity (FDR-adjusted p = 0.003, 0.321, and < 0.001; Cohen’s d = 0.303, 0.105, and 0.545) and distance (FDR-adjusted p < 0.001, 0.202, and < 0.001; Cohen’s d = −0.362, −0.127, and −0.631). The No L model showed the same trial-dependent pattern for cosine similarity (FDR-adjusted p = 0.008, 0.785, and < 0.001; Cohen’s d = 0.268, 0.029, and 0.356) and distance (FDR-adjusted p = 0.002, 0.487, and < 0.001; Cohen’s d = −0.313, −0.069, and −0.440). Direct comparison of the two structure-value controls showed that the No S model performed better than the No L model in trial 3, although the corresponding effects were small (cosine similarity: FDR-adjusted p = 0.044, Cohen’s d = 0.206; distance: FDR-adjusted p = 0.024, Cohen’s d = −0.228). Detailed statistical results are provided in Supplementary Table 10. The reduced separation among models in trial 2 and trial 3 suggests that the incremental contribution of the full combination of structure values was less pronounced for Qwen-max.

A similar pattern was observed for Qwen-flash. The full model generally outperformed all control models (lower panel of Figure 7C, and Supplementary Figure S5), although some controls showed performance comparable to that of the full model in trials 2 and 3. These included the No L model in trial 2 (cosine similarity: FDR-adjusted p = 0.292, Cohen’s d = 0.103; distance: FDR-adjusted p = 0.302, Cohen’s d = −0.099) and the No S model in trial 3 (cosine similarity: FDR-adjusted p = 0.207, Cohen’s d = 0.122; distance: FDR-adjusted p = 0.163, Cohen’s d = −0.132).

Differences among the Qwen-flash control models were concentrated mainly in trial 3. In this trial, the No S model outperformed the random walk in both cosine similarity (FDR-adjusted p < 0.001, Cohen’s d = 0.627) and distance (FDR-adjusted p < 0.001, Cohen’s d = −0.647). The No L model similarly outperformed the random walk in cosine similarity (FDR-adjusted p < 0.001, Cohen’s d = 0.375) and distance (FDR-adjusted p < 0.001, Cohen’s d = −0.439). The No S model also showed a lower patch position distance than the No L model (FDR-adjusted p = 0.021, Cohen’s d = −0.221). Their cosine-similarity difference was in the same direction but narrowly missed significance after FDR correction (FDR-adjusted p = 0.051, Cohen’s d = 0.191). Detailed statistical results are provided in Supplementary Table 12. These results indicate that the incremental contribution of the full combination of structure values was also less pronounced for Qwen-flash.

The LLM control analyses broadly reproduced the performance pattern observed in the human narratives. For both Qwen-max and Qwen-flash, the random walk and Depletion history models captured some non-random organization but remained inferior to the full model, whereas the No S and No L models generally provided closer approximations. However, the differences between the full model and the structure-value controls were smaller and more dependent on trial in the LLM-generated narratives, whereas the human full model consistently outperformed both structure-value controls across all three trials. The complete combination of second-order and third-order structure values therefore provided a more stable additional contribution to the prediction of human narratives, while its incremental contribution varied more across the LLM trials. Despite this difference, the shared pattern across human and LLM samples indicates that patch-level narrative dynamics were better explained by foraging decisions informed by structure-value information than by patch-to-patch relationships or depletion history alone.

## 4. Discussion

In the present study, we developed a computational account of thought transitions in participant-specific, high-order theme-memory networks and instantiated this account in an MVT-based model agent. The model combined two principles to predict patch-level dynamics during free spoken narration: maximizing the acquisition of structurally organized information and optimizing the explore-exploit trade-off. Several findings support this account. At the group level, the model reproduced the principal temporal organization of human patch activation and consistently outperformed random-shuffle predictions, although the degree of model conformity varied substantially across participant–theme samples. Model-predicted patch transitions also corresponded to systematic semantic changes in the narrated content, indicating that the transitions marked meaningful boundaries rather than arbitrary changes in model state. Furthermore, transition frequency varied with the perceived importance and likelihood of the narrated events. These patterns remained broadly detectable when the foraging environment was enriched with additional hub-related memory resources in the follow-up two-stage experiment. Finally, LLM-generated narratives showed detectable but more heterogeneous and less stable conformity with the same computational framework. Together, these findings support a graded account in which patch-level narrative dynamics are shaped by foraging-related decisions over multilevel structural resources, while the strength of this organization differs across themes, individuals, and generative systems.

A central assumption of our framework is that structurally organized memory content can provide an intrinsic form of informational value. Within the model, structure value refers to the amount of potentially harvestable information associated with the available second-order paths and third-order structures. When the agent is outside a patch, this value guides the selection of a resource-rich region; once inside a patch, its remaining value contributes to the decision to continue local exploitation or leave for an alternative region. This formulation is consistent with evidence that information can function as an internal reward independently of instrumental outcomes^47^, that expected information and novelty can motivate exploration^48^, and that information seeking engages neural systems associated with reward and dopaminergic valuation^49–51^. It is also compatible with accounts proposing that mind wandering reflects the reprioritization of internal goals when the currently available source of reward becomes less valuable^52^. Structure value should be understood here as a computational hypothesis about what guides narrative search. The contribution of the present model is to specify how such informational value can be derived from a structured memory environment and dynamically updated as narration proceeds.

The model further suggests that an explore-exploit trade-off governs when thought remains locally focused and when it moves toward another region of memory. Previous work on semantic fluency has established that memory retrieval often alternates between clusters of related items and switches to new clusters^12,53^. Hills and colleagues showed that these local-to-global transitions tend to occur after a semantic patch has been depleted and at a time broadly consistent with the marginal value theorem^53^. Their findings further suggested that patch switching may reflect an internally regulated search process, because patch-leaving occurred systematically as local retrieval returns declined rather than at arbitrary points in the retrieval sequence. More recent neuroimaging evidence has strengthened the strategic-search account: during semantic and phonetic fluency, hippocampal and cerebellar activity increased across within-patch retrieval and reset at switches, suggesting that transitions may be supported by a monitoring process rather than by entirely undirected movement^12^. Our findings extend these literatures from the rapid retrieval of discrete items within a predefined semantic domain to prolonged narration about personally relevant events. We also extend the representation of memory patches from local semantic clusters to participant-specific, partially overlapping structures reconstructed from repeated narratives. More importantly, rather than only asking whether observed switch timing is consistent with MVT, our framework implements an explicit value-based agent in which dynamically updated structure values guide patch-entry and stay-leave decisions, allowing prediction of active patch selection and transition dynamics along an extended narrative trajectory. In this setting, exploitation corresponds not merely to generating another member of a lexical category, but to continuing to develop content supported by an overlapping structural region of a participant-specific memory network. Exploration, in turn, involves redirecting narration toward another composite region when the expected benefits of remaining locally no longer exceed the broader environmental value. The correspondence between human and model trajectories therefore provides a computational account of how locally stable and more flexible modes of spontaneous thought may alternate over an extended narrative sequence. This interpretation is also consistent with evidence that shifts in spontaneous thought involve dynamic interactions between default-mode and control systems^10,54^ and that personally relevant goals and current concerns influence the direction of mind wandering^10,55,56^.

The control-model analyses help clarify why neither the patch representation nor the basic foraging rule alone is sufficient for this account. This distinction is important because foraging-like behavioral signatures can sometimes arise from the structure of a semantic representation without an explicit strategic switching mechanism. Abbott and colleagues showed that a random walk on a clustered semantic network derived from free-association data could reproduce several patterns previously interpreted as evidence of optimal foraging^57^. Their result initiated a question about whether apparent strategy resides in the search process or is already embedded in the representational structure traversed by the model. Hills and colleagues subsequently noted that networks derived from human association behavior may themselves contain traces of the retrieval processes that they are used to explain^58^. A recent work has extended the representational account by reporting that random walks over modern semantic embeddings enriched with descriptive information can reproduce summary-level MVT-like patterns in an animal-fluency task, whereas a more elaborate Metropolis-Hastings procedure did not provide a closer qualitative match^59^. Our results showed that the patch semantic random walk performed above the random-shuffle baseline, demonstrating that semantic relationships among patches already constrained the likely organization of narration. Thus, the topology of the theme-memory network was not merely a passive container for a separately defined decision process. Nevertheless, the random walk was substantially less predictive of human path-to-path activation dynamics than models retaining foraging-related decisions. The Depletion history model provided a complementary test: although it retained a basic process in which repeated patch activation progressively reduced availability, it did not evaluate the structural resources contained in the patches and remained markedly inferior to the full model. Generic depletion was therefore not sufficient to determine when the narrative should remain within or leave a patch. The intermediate performance of the No S and No L models further indicated that either second-order or third-order structure value could support partial prediction when combined with foraging decisions, but that neither component alone reproduced the full model’s stable advantage in human narratives. No consistent superiority of one component over the other was observed, suggesting that the two orders provide complementary hierarchical sources of predictive information. These findings point to an interaction between representation and process: patch topology defines the possible routes and contributes meaningful transition biases, whereas human narrative trajectories are better characterized when movement through this environment is guided by dynamically updated multilevel structure values.

The high-order patch representation provides an important part for human narration dynamics. Episodic experiences and narratives are segmented and integrated into structured event units^60–63^, including during internally generated spoken narratives^64^. Transitions between such units have been associated with distance-like changes in neural state space^65,66^. Hierarchical accounts further propose that narrative memory is organized across multiple levels, ranging from broad themes and major event branches to more specific event details^39^. Our framework is compatible with these accounts but adds a dynamic value-based interpretation. A patch integrates multiple pairwise narrative paths into a recurring third-order structure, allowing locally narrated details to be evaluated in relation to a broader cross-trial scaffold. This provides a possible mechanism through which thought may alternate between elaborating specific details and reorganizing higher-level meanings^67^. Importantly, patches in our framework are not simply semantic categories. Their overlap allows the same narrated path to contribute resources to several structural contexts, and the value of a patch changes as these shared resources are activated. The resulting environment is therefore both heterogeneous and dynamically coupled, creating decision conditions that are not present in a flat collection of independent semantic clusters.

The transition-related semantic analyses provide convergent evidence that the patch transitions predicted by the model correspond to boundaries in the organization of narrative content. Previous event-segmentation research has shown that continuous experiences are represented as nested event units and that transitions between events are associated with changes in neural and mnemonic states^60–66,68^. Neural meta-state transitions also align with perceived narrative events and have been proposed as general markers of changes in thought^66^. More recently, studies of spontaneous thought have shown that thought streams can be divided into semantically coherent episodes^69^ and that large semantic shifts during spontaneous memory recall and future thinking recruit default and control networks in a manner resembling externally induced event boundaries^10^. The present results add an important qualification to the idea that a thought boundary is the descent in semantic similarity. When the temporally connections of narrative content were omitted, transition-leaving paths showed relatively strong continuity. However, when the temporal unfolding of the narrative was retained, transition-related paths exhibited greater semantic change at both the hub-node and complete-path levels. These patterns suggest that semantic segmentation depends on whether the temporal organization of narrative content is preserved. A narrator can move between related contents while substantially changing the route through which those contents are developed. Patch transitions may therefore mark a reorganization of the local narrative trajectory. This extends the similarity descent signature observed at category boundaries in semantic fluency by showing that, in continuous narration, a structural boundary can preserve high-level thematic continuity while reorganizing the intermediate semantic sequence.

The association between transition rate and theme evaluations further suggests that this segmentation process varied with the functional significance of the narrated event. Themes rated as more important and more likely to be realized showed more frequent patch transitions. Importance may index the motivational priority of an event and the strength of the need to act on, prepare for, or respond to it, regardless of whether its anticipated outcome is positive or negative. Likelihood, may reflect the extent to which an imagined event is perceived as realizable and integrated with the person’s existing knowledge. Research on episodic future thinking has shown that constructing future events supports adaptive functions including planning and decision-making^70^, and that increasing episodic specificity can improve the generation of relevant means–end steps for personally concerning future events^71^. The belief that an imagined event will occur is also shaped by how coherently the event can be contextualized in relation to autobiographical knowledge, personal characteristics, and goals^72,73^. In parallel, hippocampal representations during planning are organized by context, prospective decision points, and future goals rather than by overlapping associations alone^74^. Against this background, one possibility is that important or plausible events recruit repeated movement among distinct sets of memories, anticipated outcomes, constraints, and possible actions. More frequent patch transitions could support the comparison and reorganization of these resources into alternative event trajectories or action pathways^75^. Our results support a link between patch reorganization and motivationally or prospectively significant event representations, but targeted manipulations of event importance, likelihood, and planning demands will be needed to determine the causal direction.

The supplemental cue experiment provided an additional test of whether the central findings depended exclusively on the paths used to construct the original theme-memory networks. When hub-related narration introduced additional candidate memory resources, the model continued to predict patch activation above the random shuffle baseline, and the transition-related semantic patterns were broadly reproduced. These findings suggest that the model’s predictive performance did not arise merely from replaying the finite set of paths observed in the original free narratives, although predictivity showed a modest reduction in the first trial. One possible explanation is that the additional candidate resources expanded the set of theoretically available routes without ensuring that those routes would be recruited during the actual narrative trajectory. In addition, some candidate resources may have been relevant to the general thematic scope but only weakly related to the specific unfolding of the imagined event. Nevertheless, the supplemental design provided only a partial expansion of the latent memory environment, because the additional paths were referenced to hub nodes identified in the original narratives and did not define entirely new patches. Despite this limitation, the model continued to outperform the random shuffle baseline after the candidate memory resources were incorporated, supporting the robustness of its predictive performance to an expanded set of possible paths.

Applying the same framework to human and LLM-generated narratives revealed a graded combination of similarities and differences. Both Qwen-max and Qwen-flash produced narratives with identifiable patch structure, and their trajectories showed detectable and heterogeneous correspondence with the MVT-based model. Several transition-related semantic effects were also shared across human and LLM samples. These results revise a simple dichotomy between human and LLMs: next-token generation over learned distributional regularities can produce local clustering, structured transitions, and some output-level signatures compatible with foraging. This interpretation is consistent with reports that repeated LLM generation can converge on attractor-like expressions^76^ and with the broader finding that richly structured semantic representations can allow relatively simple sampling processes to exhibit foraging-like behavior. However, the LLM samples showed weaker or less stable model conformity, particularly among the lower-conformity groups, and the full model’s advantage over the single-value and random-walk controls was more dependent on trial. The complete integration of second-order and third-order structure values therefore provided a more stable additional contribution to human than to LLM narrative prediction. Several nonexclusive mechanisms could account for this difference. Human narration about current concerns is constrained by autobiographical memory, subjective importance, anticipated consequences, and internally maintained goals, all of which may provide relatively persistent value signals as thought develops. LLM generation can reproduce statistical and semantic structure without being organized by the same participant-specific values. Explicit or internally constructed reward signals can alter LLM reasoning behavior^77^, suggesting that reward-like guidance may change generated trajectories. Most current studies of extended LLM reasoning also involve explicit problems and externally specified objectives^76–78^, whereas the present task imposed relatively weak constraints on how a theme should be developed. It remains possible that different prompting, sampling parameters, memory mechanisms, or reward-guided training would increase conformity with the human pattern. More importantly, similarity or difference in generated trajectories cannot by itself identify the internal computation that produced them.

Overall, the present work specifies a computational strategy through which spontaneous thought may search internally available memory resources under relatively low external constraint. Research on spontaneous thought has shown that self-generated concepts vary along semantic, affective, and self-relevant dimensions and that their transition dynamics relate to individual psychological characteristics^11^. Complementary neural work has shown that large shifts in thought resemble event boundaries and involve interactions between default and control networks^10,68,69^. Building on these approaches, our framework describes how successive thought units may be organized through patch selection and stay–leave decisions over a participant-specific, high-order memory environment. The findings suggest that semantic topology supplies important constraints, but that human narrative dynamics are more accurately captured when foraging-related decisions are guided by dynamically changing multilevel structure values. Patch transitions, in turn, mark a form of semantic segmentation and reorganization that may support the internal exploration of personally significant events and possible action pathways. Future work should determine how these computations are implemented neurally, how they change under different goals and affective states, and when strategic internal foraging facilitates or impairs planning, problem solving, and adaptive behavior.

## Acknowledgments

This work was supported by the National Natural Science Foundation of China (www.nsfc.gov.cn/; grant numbers 82072000 and 81671769), the Natural Science Foundation of Heilongjiang Province, China (http://kjt.hlj.gov.cn/, grant number YQ2023H016), and the State Key Laboratory of Space Environment Interaction with Matters.

## Supplementary Materials

### Supplementary Methods

#### S1. Normalization and group-level averaging of observed and model predicted patch activation matrices

The observed and model predicted patch activation matrices differed in size across participant-theme samples because both the number of patches and the number of hub-to-hub paths varied across samples. To visualize observed and model-predicted patch activation patterns at the group level, we projected each matrix onto a common two-dimensional normalized coordinate system, resampled it to a fixed spatial resolution, and then averaged the resampled matrices across participant-theme samples.

For participant-theme sample *k* in trial *i*, the observed patch activation matrix was denoted by 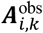, and the model predicted matrix averaged across the 2000 simulations described in Section 2.6 was denoted by 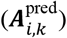. Both matrices had dimensions *P*_*k*_ × *T*_*i,k*_, where *P*_*k*_ was the number of patches and *T*_*i,k*_ was the number of temporally ordered hub-to-hub paths in that trial. Rows represented patches, whereas columns represented paths in their original narrative order. Entries in 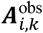 indicated whether a given path activated a given patch. Entries in 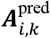 represented the proportion of simulations in which the corresponding patch was activated at that path position.

Before resampling, patches were reordered according to their temporal activation patterns in the observed matrix. Patches activated during the trial were ordered primarily by their mean activation position across the narrative sequence. When two or more patches had the same mean activation position, they were further ordered according to their first activation position and, when these were also identical, according to their last activation position. Patches that were not activated during the trial were placed after the activated patches. The row permutation derived exclusively from the observed matrix was then applied identically to both the observed and predicted matrices. Thus, the paired matrices shared the same patch order, and the predicted matrix was not reordered according to the model predictions themselves.

The path and patch dimensions were subsequently mapped onto the interval [0,1]. For sample *k* in trial *i*, the normalized center of path *t* was defined as

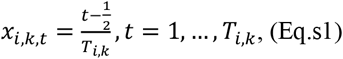

Similarly, after patch reordering, the normalized center of patch rank *p* was defined as

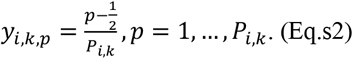

Accordingly, the horizontal axis represented normalized position within the narrative path sequence, whereas the vertical axis represented normalized patch rank based on the temporal activation order observed in the observed matrix. The normalized patch axis therefore represented the relative temporal organization of patches within each sample.

Each reordered matrix was then resampled from its original *P*_*k*_ *× T*_*i,k*_ resolution to a common 50 × 50 grid, corresponding to 50 normalized patch-rank bins and 50 normalized path-position bins. We used area-overlap resampling, in which each original matrix entry was treated as a rectangular bin in the normalized [0,1] × [0,1] space. The value assigned to each target bin was calculated as the weighted average of the original bins overlapping that target bin, with weights determined by their proportional spatial overlap.

Let 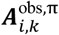 and 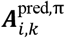 denote the reordered observed and predicted matrices, respectively. Their resampled forms were calculated as

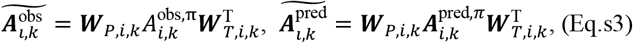

where ***W***_*P,i,k*_ was the 50 × *P*_*k*_ overlap-weight matrix for the patch dimension and ***W***_*T,i,k*_ was the 50 × *T*_*i,k*_ overlap-weight matrix for the path dimension. Each element of an overlap-weight matrix represented the normalized spatial overlap between an original bin and a target bin along the corresponding dimension. The same transformation was applied separately to the paired observed and predicted matrices. This procedure converted matrices with different numbers of patches and paths to a common resolution while preserving their overall mean activation levels.

The resampled observed and predicted matrices were then averaged separately within each participant group and trial. For group *G*, containing *N* valid participant-theme samples in trial *i*, the group-level observed matrix and predicted matrix were calculated as

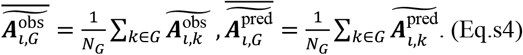

Each participant-theme sample contributed equally to the group average, irrespective of its original number of patches or hub-to-hub paths. Group-level matrices were calculated separately for each trial and for the full set of included participant-theme samples, Group H1, and Group H2. The resulting matrices represented the average observed and model predicted patch activation patterns as functions of normalized narrative position and normalized patch rank.

#### S2. Classification of human and Qwen samples based on model-prediction performance

To characterize individual differences in the extent to which human thought dynamics were captured by the MVT-based model, we classified human participant-theme samples according to the model’s predictive advantage over the random-shuffle baseline. We then transferred the resulting human cluster structure to the Qwen max and Qwen flash samples to determine whether each Qwen-generated sample was more similar to the human group showing higher or lower conformity with the model-predicted foraging-like dynamics.

##### Classification of human samples into H1 and H2

For each human participant-theme sample, we obtained the cosine similarity and patch position distance of the MVT-based model for each of the three narration trials. Both performance measures were standardized relative to the corresponding random-shuffle baseline. The direction of the distance score was reversed so that, for both measures, a larger standardized value indicated a greater predictive advantage of the MVT-based model over the random-shuffle baseline.

Specifically, the standardized cosine-similarity and distance scores were defined as

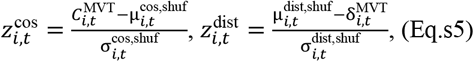

where 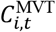 and 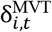 denote the cosine similarity and patch position distance, respectively, of human sample *i* in trial *t*. The terms 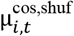 and 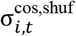 denote the mean and standard deviation of the corresponding random-shuffle cosine similarity distribution, and 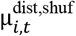 and 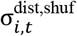 denote those of the random-shuffle distance distribution. The reversed numerator of the distance score reflected the fact that lower distance indicated better model prediction.

Each human sample was therefore represented by a six-dimensional model-performance vector 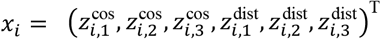. Samples with a missing or nonfinite value in any of the six dimensions were excluded from clustering. Of the 97 human participant-theme samples, 94 had complete six-dimensional features and were included, whereas three were excluded because at least one feature could not be calculated.

Candidate numbers of clusters ranged from *k* = 2 to *k* = 9. For each value of *k*, k-means was fitted using 100 different initial centroid configurations. Solutions containing any cluster with fewer than three samples were discarded. Among the remaining valid solutions, the number of clusters with the highest silhouette score was selected. This procedure selected a nine-cluster solution.

We subsequently consolidated the nine original clusters into two interpretable human groups. First, we identified the cluster exhibiting the strongest overall model-conformity profile. For each original cluster *C*_*c*_, the mean of each of the six random-shuffle-standardized features was calculated as 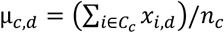, where *n*_*c*_ denotes the number of samples in cluster *C*_*c*_. A cluster was considered a candidate template if its mean exceeded the corresponding across-sample mean in all six feature dimensions. For each cluster, the joint model-conformity score was defined as 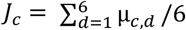.

Among the clusters satisfying the six-dimensional criterion, the cluster with the highest *J*_*c*_ was selected as the reference template cluster *C*_*temp*_. In the present data, original cluster 3 was selected as the template. To quantify the similarity of the remaining clusters to the template, we first calculated a joint performance score for each human sample as 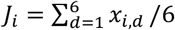. The scores of all samples assigned to cluster *C*_*c*_ formed its empirical joint-score distribution: *F*_*c*_ *=* {*J*_*i*_: *i* ∈ *C*_*c*_}.

For every non-template cluster, we calculated the first-order Wasserstein distance (*W*_1_) between its joint-score distribution and that of the template cluster: *D*_*c*_ = *W*_1_(*F*_*c*_, *F*_*r*_). The resulting Wasserstein distances of the eight non-template clusters were then divided into two sets using one-dimensional k-means clustering with *k* = 2, 50 initial centroid configurations. The set with the smaller distance centroid was considered more similar to the template cluster, whereas the set with the larger distance centroid was considered less similar. The template cluster and the clusters in the nearer set were combined to form Group H1, whereas the clusters in the farther set formed Group H2:

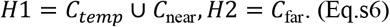

Of the 94 human samples included in the clustering, 71 were assigned to H1 and 23 to H2. H1 therefore comprised samples whose model-performance profiles were more similar to the strongest model-conformity template and was interpreted as showing greater correspondence with the model predicted foraging-like strategies. H2 comprised samples whose performance profiles were more distant from the template and was interpreted as showing comparatively lower correspondence with these strategies.

##### Classification of Qwen max and Qwen flash samples into Q1 and Q2

To establish a direct correspondence between the Qwen-generated samples and the human H1, H2 classification, we did not independently cluster the Qwen max and Qwen flash samples. Instead, the original nine-cluster human structure and its subsequent H1, H2 grouping were retained as a fixed reference.

As for the human samples, each Qwen-generated participant-theme analogue was represented by a six-dimensional vector containing the random-shuffle-standardized cosine-similarity and distance-performance scores from the three trials, 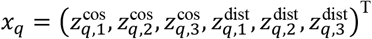. Only samples with finite values in all six dimensions were retained. The same classification procedure was applied separately to Qwen max and Qwen flash.

Using the 94 human samples with complete features, we calculated the centroid of each original human cluster in the six-dimensional random-shuffle-standardized performance space as 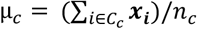. For each Qwen sample *q*, its Euclidean distance from every human cluster centroid was calculated as *d*_*q,c*_ = ∥ *x*_*q*_ − μ_*c*_ ∥_2_. The Qwen sample was assigned to the nearest original human cluster: 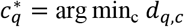. Its final group label was then determined by whether the nearest human cluster belonged to H1 or H2,

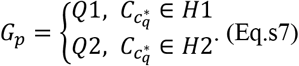

Thus, Q1 comprised Qwen samples whose six-dimensional model-performance profiles were closest to an original human cluster assigned to H1, whereas Q2 comprised samples closest to a human cluster assigned to H2. This procedure preserved heterogeneity among the original human clusters rather than representing H1 and H2 using only one centroid per group.

As a continuous measure of relative group proximity, we additionally calculated the difference between each Qwen sample’s distance to the nearest H2 cluster and its distance to the nearest H1 cluster:

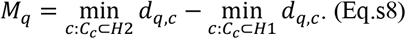

A positive *M*_*q*_ indicated that the sample was closer to an H1 cluster than to any H2 cluster, whereas a negative value indicated greater proximity to an H2 cluster. Of the 122 Qwen max samples, 117 had complete six-dimensional features and were included in the classification, five were excluded because of incomplete features. Among the included Qwen max samples, 57 were classified as Q1 and 60 as Q2. Of the 141 Qwen flash samples, 139 had complete features and were included, whereas two were excluded. Among the included Qwen flash samples, 61 were classified as Q1 and 78 as Q2.

#### S3. Follow-up two-stage experiment: detailed procedures and analyses

Because the content produced during unconstrained free narration may not exhaust the memories and thoughts potentially available within a given theme, we recruited additional 13 healthy adults (average age 25.7, range from 18 to 30, 6 males and 7 females) to complete a two-stage narrative experiment. Phase 1 was identical to the original free spoken narrative task. Participants first completed the pre-test questionnaire, from which four personally relevant themes spanning affective valence and subjective importance were selected. They subsequently completed three 15-min free-narration trials separated by 5-min breaks.

Participants returned 2–3 days later for an individualized hub-cued narrative elaboration session. Before Phase 2, each participant’s Phase 1 narratives were preprocessed using the same procedure as in the main experiment, including expression simplification, temporal connection of successive textual nodes, and semantic merging of similar nodes. A participant- and theme-specific memory network was then constructed. Themes were eligible for Phase 2 if they contained narrative content in all three Phase 1 trials and yielded a valid patch network in which the extracted patches collectively covered all observed hub-to-hub paths. Of the 13 recruited participants, one was excluded because one Phase 2 trial contained no usable narrative response, and another was excluded because none of their Phase 1 themes yielded a valid patch network. The final analysis therefore included 11 participants and 25 eligible participant-theme datasets.

For each eligible theme, four hub nodes were selected as individualized cues. Candidate hubs were primarily selected to maximize their occurrence across the three Phase 1 trials while minimizing semantic similarity among the selected hub nodes. When more than four hub nodes satisfied these criteria, the candidates were ranked by network degree, and the four highest-degree hub nodes were retained. Before Phase 2, participants were informed that they would be shown brief statements taken from their previous narratives and would be asked to elaborate on any content that naturally came to mind within the corresponding theme. They were not informed that the selected statements represented hub nodes in their narrative networks.

During each Phase 2 trial, the theme label and one of its four cues remained on the screen for 60s, during which the participant spoke freely about thoughts naturally elicited by the cue while remaining within the scope of the displayed theme. Each eligible theme–cue combination was presented once per trial, and presentation order was randomized independently within each trial. Phase 2 comprised three trials separated by 5-min breaks, and each theme–cue combination was presented once per trial, resulting in three presentations in total. Spoken responses were audio-recorded, transcribed, and preprocessed using the same text-processing procedure as for the Phase 1 narratives.

##### Construction of cue-elicited expanded paths

The Phase 2 responses were used to identify cue-elicited expanded paths that were not expressed in the original Phase 1 narrative sequence. For each eligible theme, the four cues presented across three Phase 2 trials produced 12 cue-by-trial response sequences. For each cue-by-trial response sequence, we preserved the original temporal order of the text segments and divided the sequence into one or more consecutive expanded paths according to the sequence in which matches to Phase 1 hub nodes occurred. Each preprocessed Phase 2 text node was embedded using the same semantic-embedding model as in Phase 1 and compared with all hub nodes in the corresponding Phase 1 theme-memory network. The same theme-specific semantic-similarity threshold used in the Phase 1 network construction was applied, these thresholds ranged from 0.86 to 0.90 across themes.

The cue corresponded to a Phase 1 hub node and was treated as the starting hub node of the first expanded path. For each subsequent Phase 2 text node, we identified the most semantically similar Phase 1 hub node. When the highest semantic similarity exceeded the predefined threshold and the matched hub differed from the current starting hub, a directed expanded path was defined between the two hubs. The newly matched hub then served as the starting hub for the next expanded path. Accordingly, a cue-by-trial response sequence could yield temporally ordered series of expanded hub-to-hub paths.

Each expanded path was subsequently mapped to all existing Phase 1 patches containing both of its starting and ending hubs. Expanded paths that could not be mapped to any existing patch did not contribute to the augmented model. Importantly, these paths were treated as cue-elicited expanded structural paths rather than as additional observations in the empirical Phase 1 narrative trajectory.

##### MVT-model simulation with expanded paths

To examine whether additional cue-elicited paths altered the model’s account of Phase 1 patch transitions, we reran the MVT-based model after augmenting the internal path structure of the existing Phase 1 patches. Expanded paths did not create new patches, were not incorporated into the observed Phase 1 path trajectory, and did not directly produce patch activations. The number and temporal order of experienced Phase 1 hub-to-hub paths therefore remained unchanged.

For each patch and trial context, an augmented local path graph was constructed from the original observed paths and the expanded paths mapped to that patch. An observed–expanded or expanded–expanded connection was assigned when the two paths shared at least one hub endpoint. The adjacency and degree matrices of each augmented local graph were then recomputed. Consequently, the expanded paths changed both their own degrees and the degrees of the original paths with which they shared hub endpoints.

The third-order structural contribution was recalculated from the normalized Laplacian spectral entropy of the augmented local path graph,

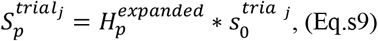

where 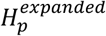 denotes the spectral entropy of patch *p* in Phase 1. As the original Phase 1 paths were experienced, they were removed from the remaining-path graph used to calculate entropy, whereas the expanded paths remained structurally available. Expanded paths could therefore alter both the initial value of 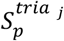 and its subsequent change as observed paths were depleted.

The second-order contribution of each path was recalculated from its degree in the augmented local graph:

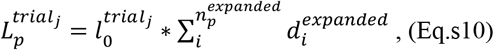

where 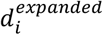 is the augmented degree of path *i* in patch *p*, j denotes the trial associated with that path. The values of the remaining observed paths were dynamically depleted as in the original model. In contrast, the expanded-path contributions were summed into a patch-specific, nondepleting structural bonus because these paths were not experienced in the Phase 1 trajectory.

To maintain a common comparison coordinate, the patch ordering used for the patch activation matrices, patch position trajectories, and distance calculation was fixed to the ordering derived from the original Phase 1 network rather than. Model performance was then evaluated using the same patch-activation similarity and position-distance measures as in the main analysis.

#### S4. Details of the control models

To determine whether the predictive performance of the MVT-based model depended on its foraging-related decision rules and multilevel structural value representations, we constructed a set of control and ablation models. These included four random walk models using different patch-transition rules, a No S model that removed the third-order structural value, a No L model that removed the second-order path value, an Equal L model that removed path-specific variation in second-order value, and a Depletion history model in which patch values depended only on their cumulative activation histories.

All models were implemented on the same participant-specific, theme-specific patch networks and used the same trial-specific patch–path membership structure as the full MVT-based model. They were evaluated over the same trial lengths and against the same temporally ordered human path sequences. Each model was simulated 2000 times for each participant-theme sample. Cosine similarity and patch-position distance were calculated separately for each simulation and then averaged across simulations, as described in Section 2.6. Within each simulation, patch-position distance was additionally averaged over 1000 random permutations of patch order.

##### Random-walk models

The random walk models tested whether the observed patch activation dynamics could be reproduced without the foraging-related decision process of the full model, using only the patch network and predefined patch-to-patch transition probabilities. We implemented four variants in which transition weights were determined by patch semantic similarity, patch overlap, patch size, or uniform transition probabilities.

At the beginning of each trial, all observed paths were treated as unconsumed. The model randomly selected an initial primary patch from the set of patches containing at least one unconsumed path. It then visited the earliest remaining path associated with that patch. If the visited path belonged to multiple patches, all patches containing that path were simultaneously activated, whereas the selected patch was designated as the primary patch. The visited path was then marked as consumed and could not be visited again. Once a patch contained no remaining paths, it was removed from the set of candidate primary patches. At each subsequent step, the next primary patch was sampled from the transition probability distribution associated with the preceding primary patch, after excluding patches with no remaining paths and renormalizing the probabilities over the remaining candidates. The number of remaining paths determined whether a patch remained eligible for selection or co-activation, but did not otherwise modify its base transition probability.

The four random-walk models differed only in the construction of the patch-to-patch transition-weight matrix.

In the *UNIFORM random walk*, all eligible target patches were assigned equal base transition weights. After patches with no remaining paths were excluded, the weights of the remaining patches were renormalized. This model provided a control for patch transition without any structural or semantic preference.

In the *PATCH SIZE random walk*, the transition weight from any source patch *i* to target patch *j* was proportional to the total number of paths contained in the target patch as 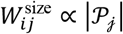, where *P*_*j*_ denotes the complete set of paths belonging to patch *j*. This model controlled for the possibility that larger patches were more likely to be visited simply because they contained more paths.

In the *PATCH OVERLAP random walk*, the directed transition weight from patch *i* to patch *j* was defined as 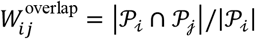. Because the denominator depended on the source patch, this transition matrix was generally asymmetric. Path identity was defined jointly by its starting hub node, ending hub node, and trial membership. Consequently, paths with the same hub-node endpoints but occurring in different trials were treated as distinct paths. This model tested whether overlap among the path sets of different patches was sufficient to explain the observed transition dynamics.

In the *PATCH SEMANTIC random walk*, the transition weight between two patches was determined by the mean semantic similarity between all hub nodes contained in the two patches. Let ℋ_*i*_ and ℋ_*j*_ denote the sets of hub nodes associated with patches *i* and*j*, respectively. Their semantic transition weight was defined as 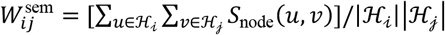, where *S*_node_(*u, v*) denotes the cosine similarity between the semantic embeddings of hub nodes *u* and *v*. The transition weights in each row were normalized to sum to one after ineligible patches were excluded. This model tested whether patch transitions could be explained solely by the complete semantic relationships between patches.

None of the random walk models calculated and used the patch value 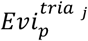 or the MVT-based stay and leave values, *V*_stay_ and *V*_leave_.

##### No S model

The No S model tested the contribution of the third-order structural value to model prediction. To preserve comparability with the full model, the original patch definitions and patch overlap structure were retained, but the third-order value *S* and the additional benefits attributable to the co-activated patches V(auto S, t) were removed from patch-value calculations. Patch value therefore depended only on the second-order path-value component as 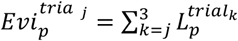. As in the full model, paths were processed according to their observed temporal order. When a path belonged to multiple patches, those patches were co-activated, but this co-activation did not generate an additional high-order structure value contribution. The model retained the foraging-related stay or leave decision based on *L*. The expected immediate path value available from the current patch contributed to *V*_stay_, whereas the mean path value obtained over previous steps contributed to *V*_leave_. The model left the current patch when *V*_stay_ < *V*_leave_, and was forced to leave when the current patch contained no remaining paths. After leaving, it entered the candidate patch with the greatest remaining current and future *L*-based evidence.

##### No L model

The No L model tested the predictive contribution of second-order path value. All path-specific *L* values were set to zero, while the original patch definitions, patch overlap structure, memberships, third-order structural value, and foraging-related decision process were retained. Although paths continued to be processed in their observed temporal order, they no longer contributed second-order value to patch-entry evidence or to the stay-leave comparison. Stay-leave decision was therefore determined only by the remaining S-related components, including the third-order value and co-activation related value.

##### Equal L model

The Equal L model tested whether the performance of the full model depended on path-specific differences in second-order structure value. This model retained the *L* component but removed the degree-dependent variation among paths within the same trial.

##### Depletion history model

The Depletion history model tested whether cumulative patch-activation history and the resulting value depletion were sufficient to explain the observed patch transition patterns in the absence of structure value. This model removed both the third-order structure value *S* and the second-order structure value *L*, as well as other path-specific value differences. Patch selection instead depended only on the number of times each patch had previously been activated. All patches were assigned an initial value of 1. The value of patch *p* at path *t* was defined as 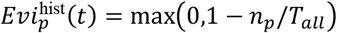, where *n* (*t*) denotes the cumulative number of previous activations of patch *p*, and *T*_all_ = *T*_1_ + *T*_2_ + *T*_3_ denotes the total number of paths across the three trials. Each activation therefore reduced the patch value by a fixed amount of (1/*T*_all_).

The cumulative activation count included both primary activation and automatic activation caused by patch overlap. If a path activated multiple overlapping patches, the activation histories of all corresponding patches were incremented. Depletion history was carried across trials, patches frequently activated in earlier trials consequently had lower entry values in later trials. Whenever a new patch was selected, the model entered the eligible patch with the highest current history-dependent value. Because all paths provided the same constant immediate reward and no path-specific value differences were retained, *V*_stay_ and *V*_leave_ were equal whenever the current patch still contained available paths, which makes the decision of leaving from the current patch stochastic. Thus, the Depletion history model primarily specified which patch was entered after leaving, rather than providing a structured prediction of when the current patch should be left.

---

**Supplementary Table 1:**
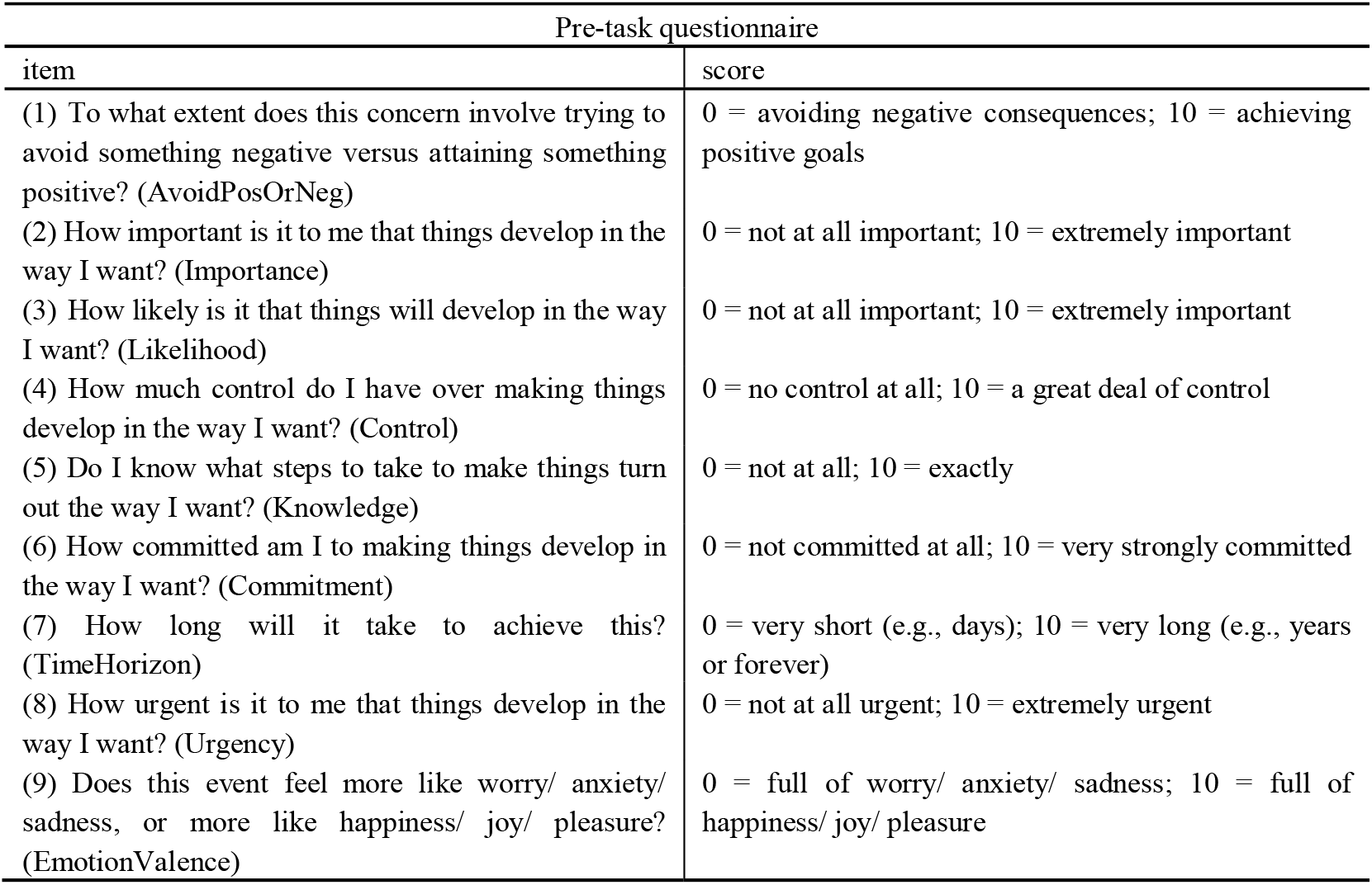
pre-task questionnaire.

| Pre-task questionnaire |  |
| --- | --- |
| item | score |
| (1) To what extent does this concern involve trying to avoid something negative versus attaining something positive? (AvoidPosOrNeg) | 0 = avoiding negative consequences; 10 = achieving positive goals |
| (2) How important is it to me that things develop in the way I want? (Importance) | 0 = not at all important; 10 = extremely important |
| (3) How likely is it that things will develop in the way I want? (Likelihood) | 0 = not at all important; 10 = extremely important |
| (4) How much control do I have over making things develop in the way I want? (Control) | 0 = no control at all; 10 = a great deal of control |
| (5) Do I know what steps to take to make things turn out the way I want? (Knowledge) | 0 = not at all; 10 = exactly |
| (6) How committed am I to making things develop in the way I want? (Commitment) | 0 = not committed at all; 10 = very strongly committed |
| (7) How long will it take to achieve this? (TimeHorizon) | 0 = very short (e.g., days); 10 = very long (e.g., years or forever) |
| (8) How urgent is it to me that things develop in the way I want? (Urgency) | 0 = not at all urgent; 10 = extremely urgent |
| (9) Does this event feel more like worry/ anxiety/ sadness, or more like happiness/ joy/ pleasure? (EmotionValence) | 0 = full of worry/ anxiety/ sadness; 10 = full of happiness/ joy/ pleasure |

**Supplementary Table 2:** the correlations between pre-test indicators and model predicted transition rates. Likelihood and Importance showed significant positive correlations between transition rates after FDR correction, though C1 that Likelihood belonged to was non-significant correlated with transition rates.

| Correlations between theme-ratings and transition rates |  |  |  |  |  |  |
| --- | --- | --- | --- | --- | --- | --- |
| Variable | n | Pearson_r | CI_low | CI_high | Pearson_p | Pearson_p_FDR |
| C1 | 71 | 0.216 | -0.018 | 0.428 | 0.07 | 0.14 |
| C2 | 71 | 0.306 | 0.079 | 0.504 | 0.009 | 0.038* |
| C3 | 71 | 0.06 | -0.176 | 0.289 | 0.622 | 0.622 |
| C4 | 71 | 0.134 | -0.102 | 0.357 | 0.264 | 0.351 |
| Likelihood | 71 | 0.338 | 0.114 | 0.529 | 0.004 | 0.018* |
| Control | 71 | 0.152 | -0.084 | 0.372 | 0.206 | 0.309 |
| Knowledge | 71 | 0.094 | -0.142 | 0.32 | 0.435 | 0.559 |
| Commitment | 71 | 0.178 | -0.058 | 0.395 | 0.138 | 0.309 |
| Importance | 71 | 0.338 | 0.113 | 0.529 | 0.004 | 0.018* |
| Urgency | 71 | 0.2 | -0.035 | 0.414 | 0.095 | 0.284 |
| TimeHorizon | 71 | 0.06 | -0.176 | 0.289 | 0.622 | 0.622 |
| AvoidPosOrNeg | 71 | 0.157 | -0.079 | 0.377 | 0.19 | 0.309 |
| EmotionValence | 71 | 0.076 | -0.16 | 0.304 | 0.526 | 0.592 |

**Supplementary Table 3:**
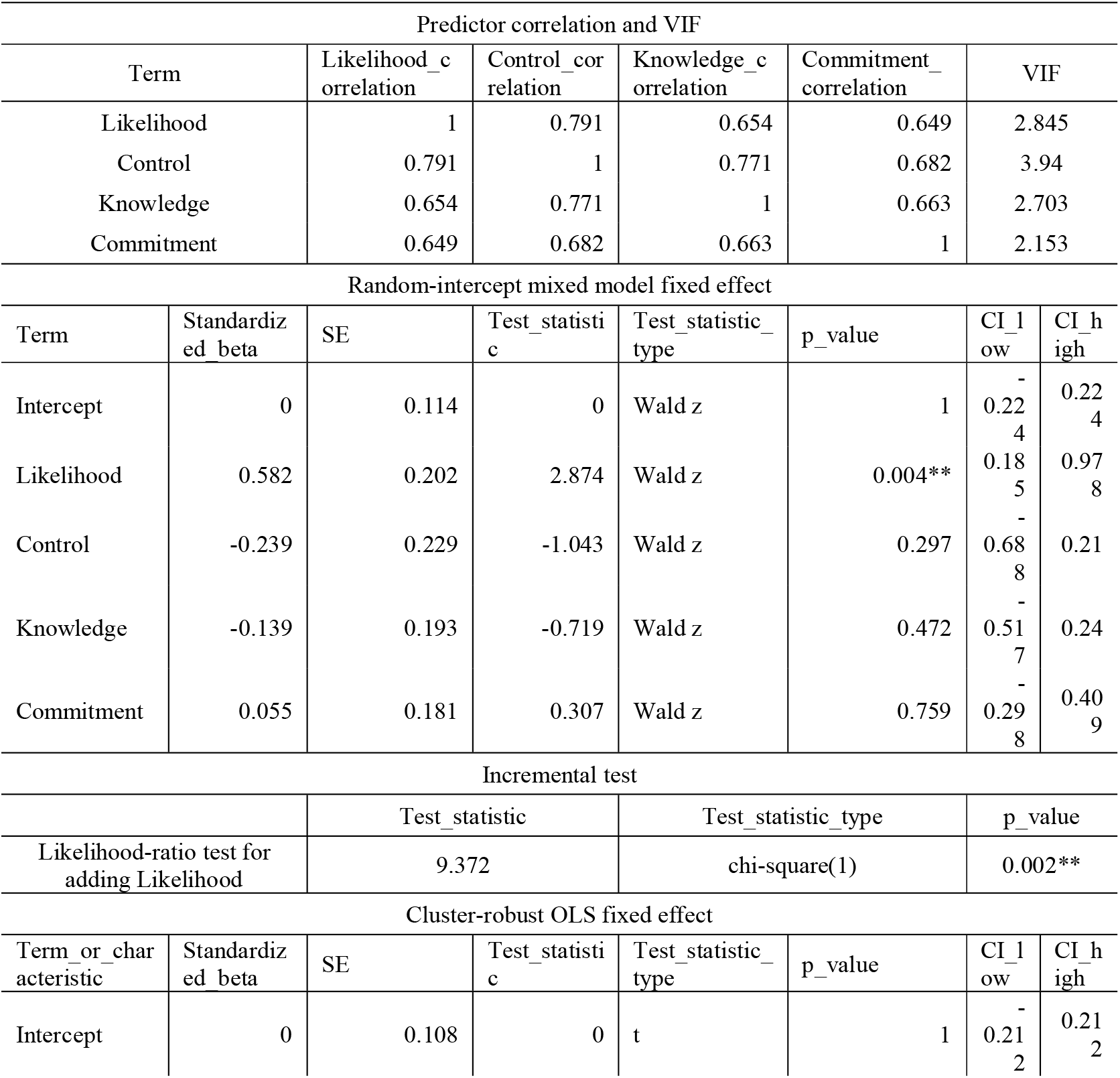
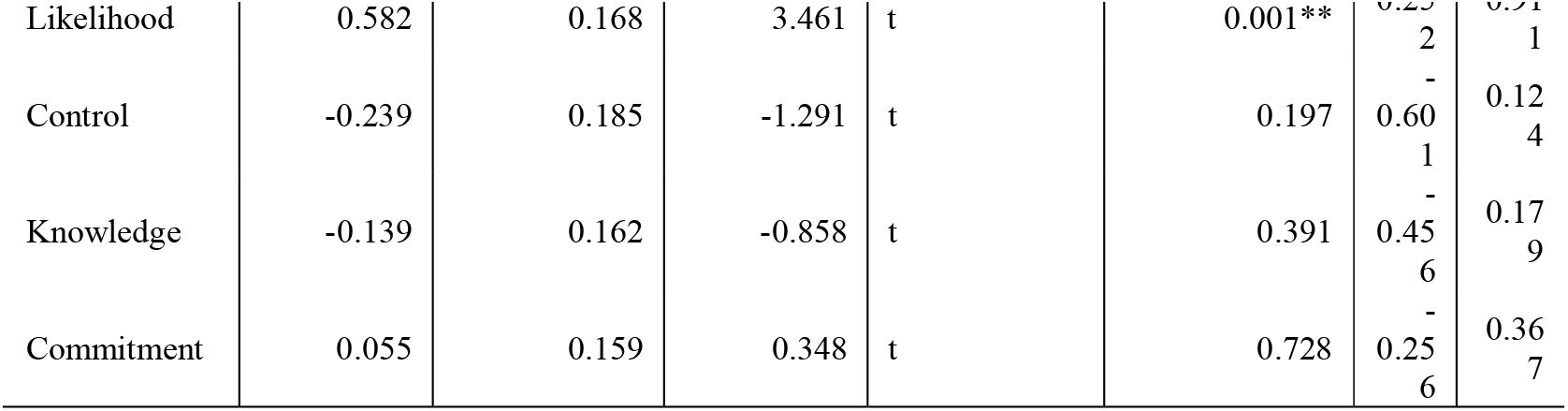
The mixed-effects model included a participant-specific random intercept to account for multiple themes contributed by the same participant. Predictor correlations ranged from 0.649 to 0.791, but all variance inflation factors were below 4. A sensitivity analysis using ordinary least-squares regression with standard errors clustered by participant therefore was conducted; this analysis produced the same substantive conclusion, with Likelihood remaining significant and the other Factor 1 indicators remaining nonsignificant. The likelihood-ratio test compared models with and without Likelihood.

**Supplementary Table 4:**
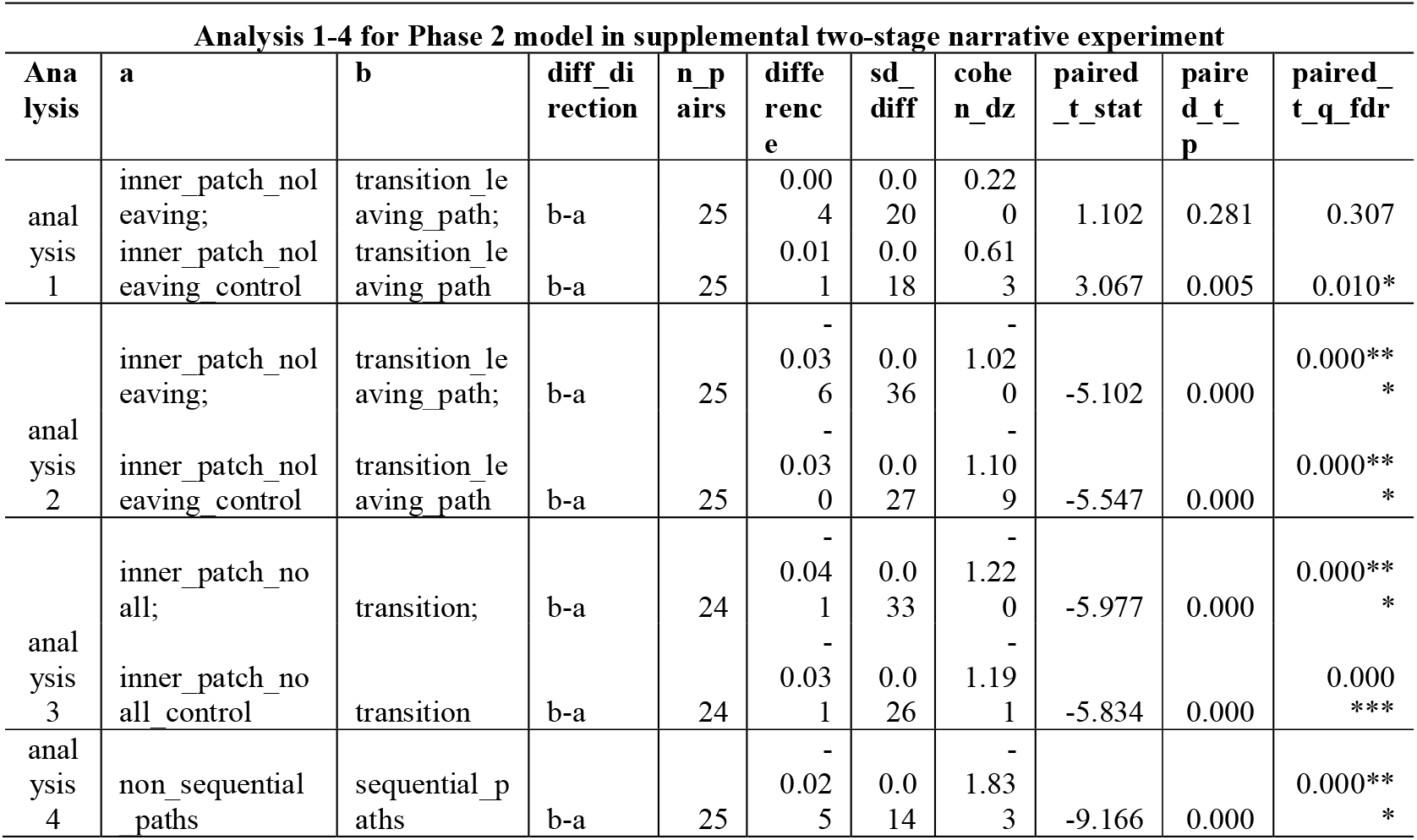
results of the four transition-related semantic analyses when candidate memory resources were added to the model simulation (Phase 2 model). Analyses 1–4 were significant and showed effect directions consistent with the results of Phase 1 model.

**Supplementary Table 5:**
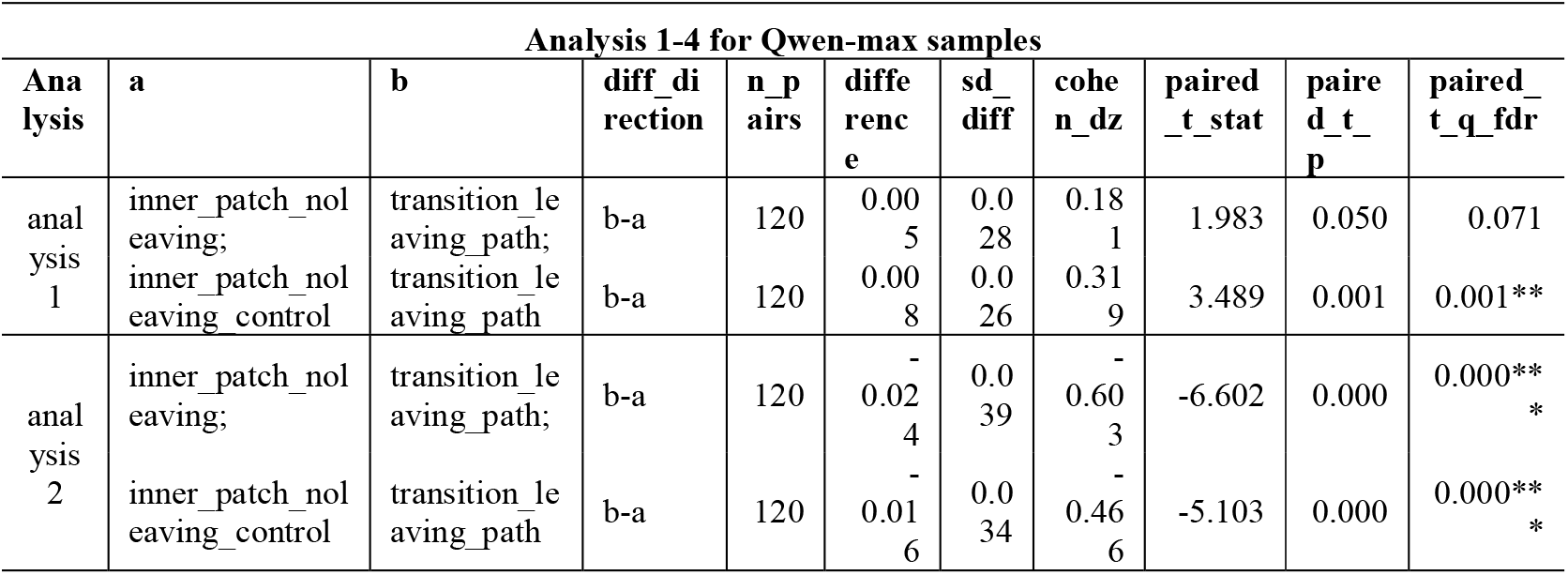
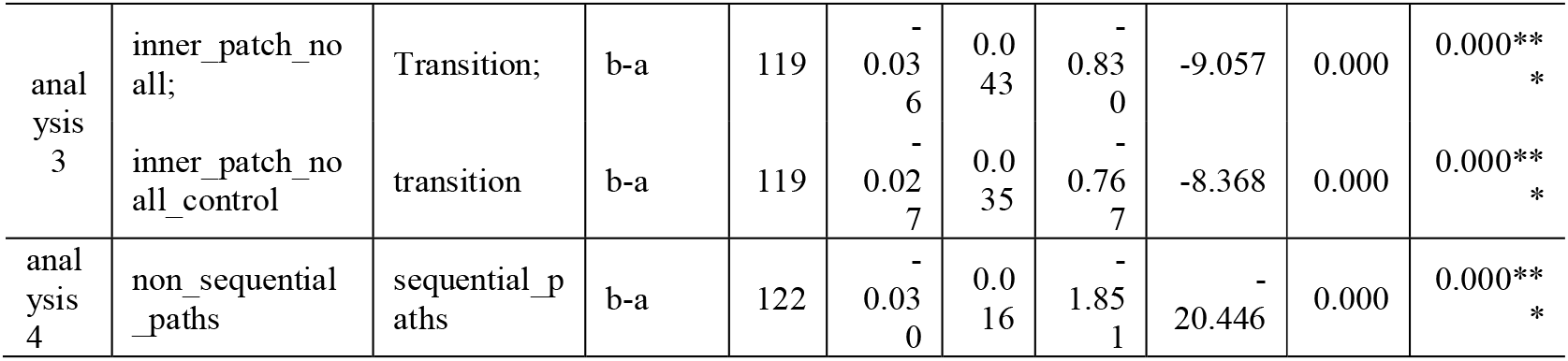
results of the four transition-related semantic analyses in Qwen-max data. Analyses 1–4 were significant and showed effect directions consistent with the human results.

**Supplementary Table 6:**
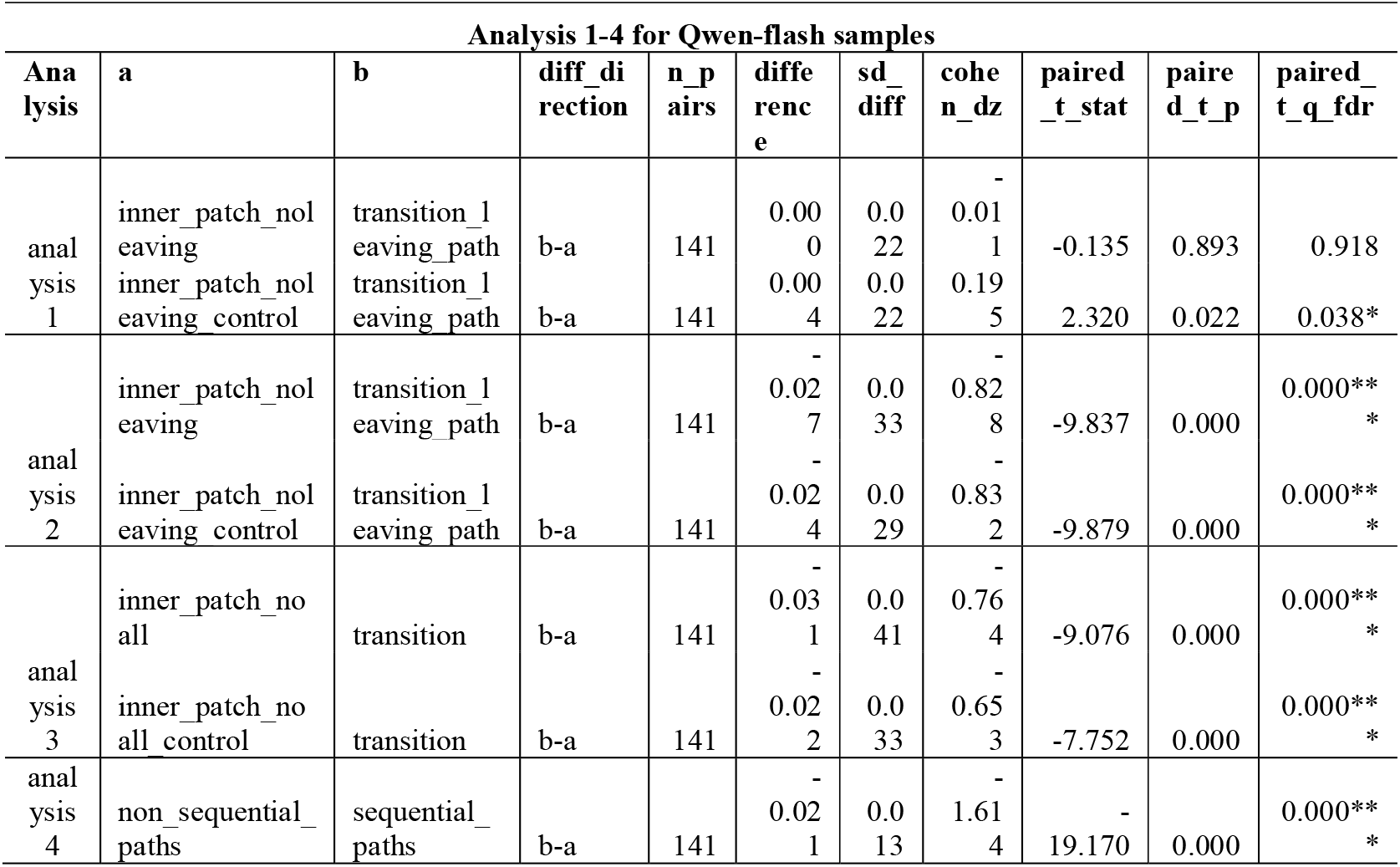
results of the four transition-related semantic analyses in Qwen-flash data. Analyses 1–4 were significant and showed effect directions consistent with the human results.

**Supplementary Table 7:**
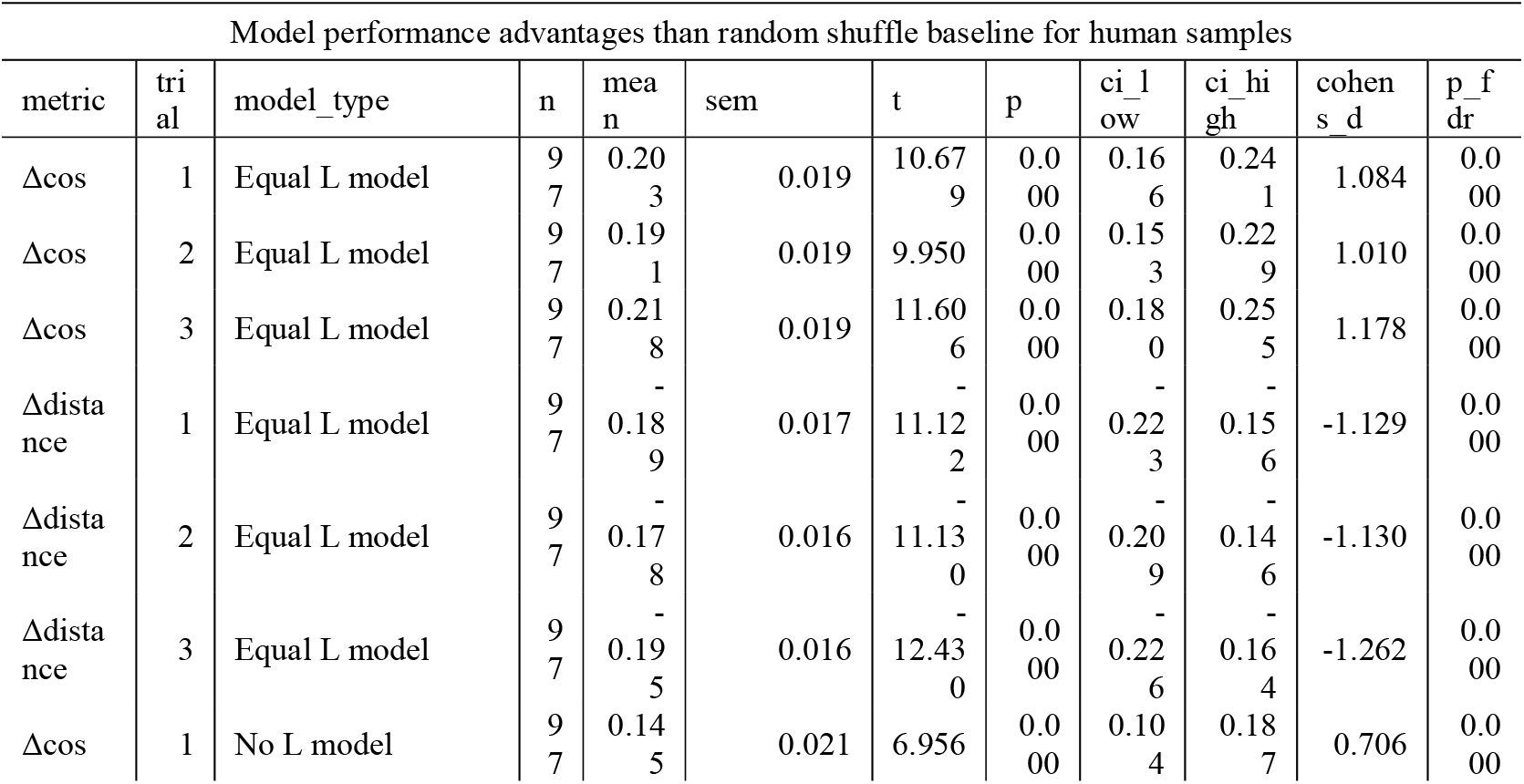
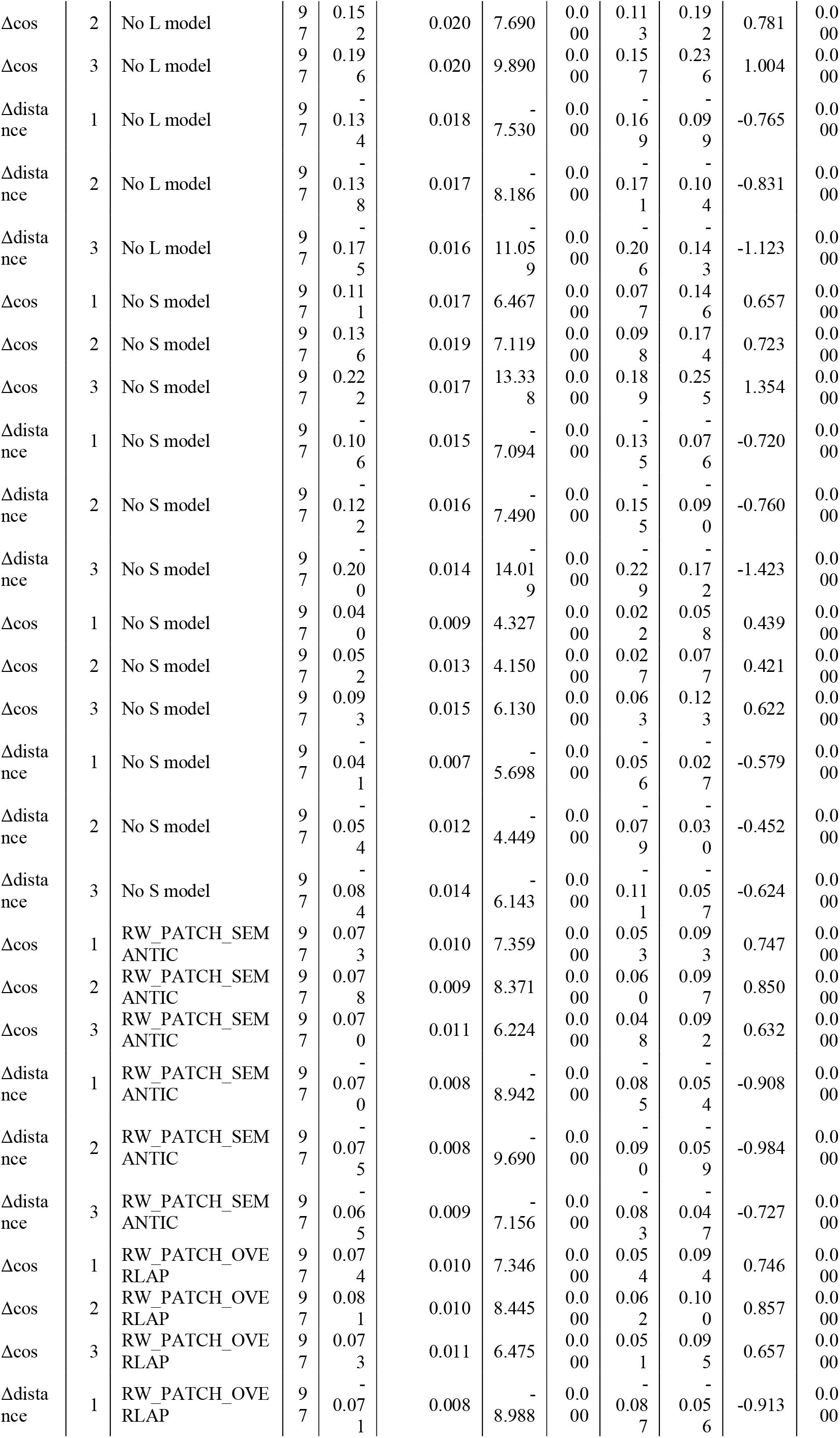
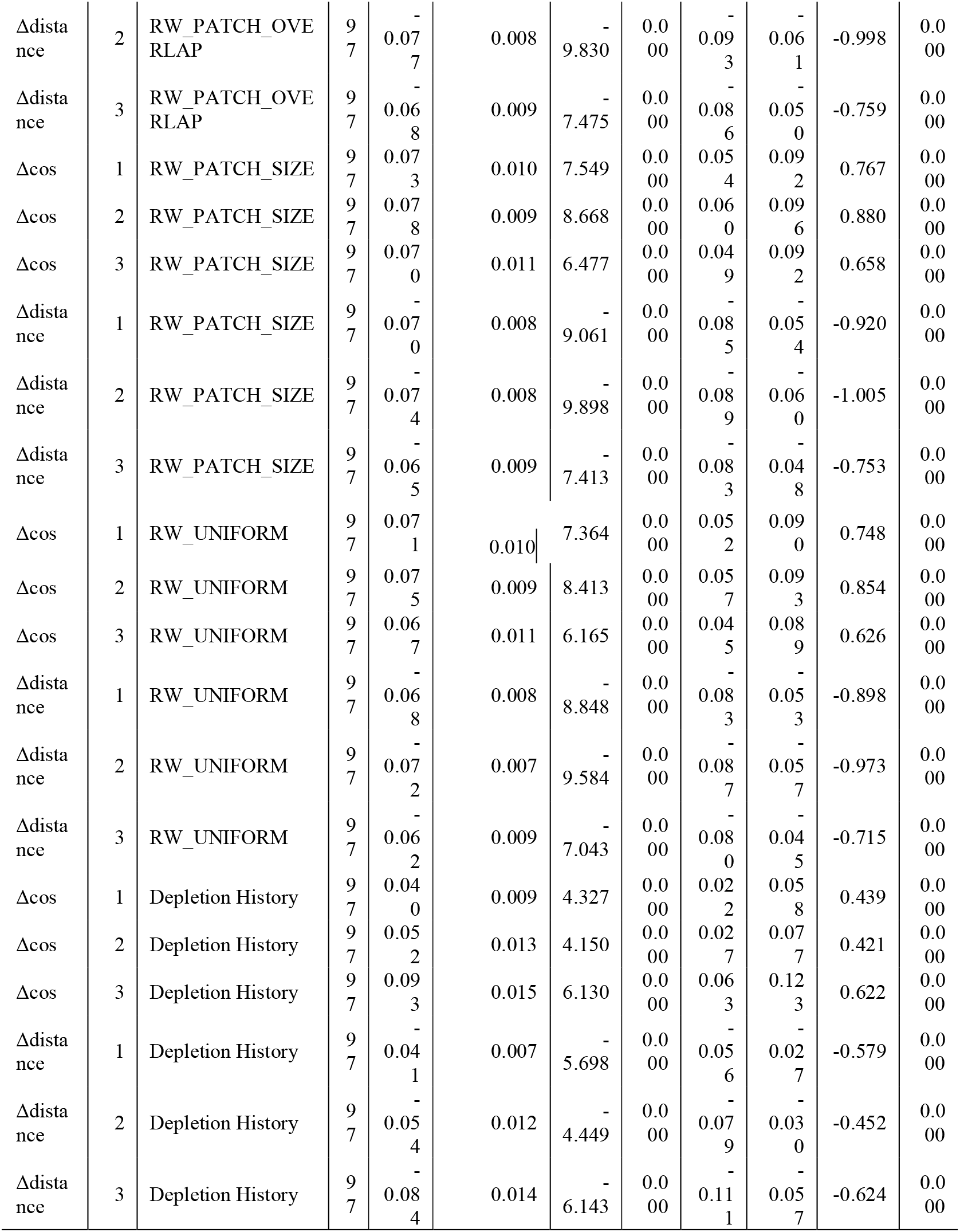
differences between control models and random shuffle baseline in human data. All control models outperformed than random shuffle baseline.

**Supplementary Table 8:**
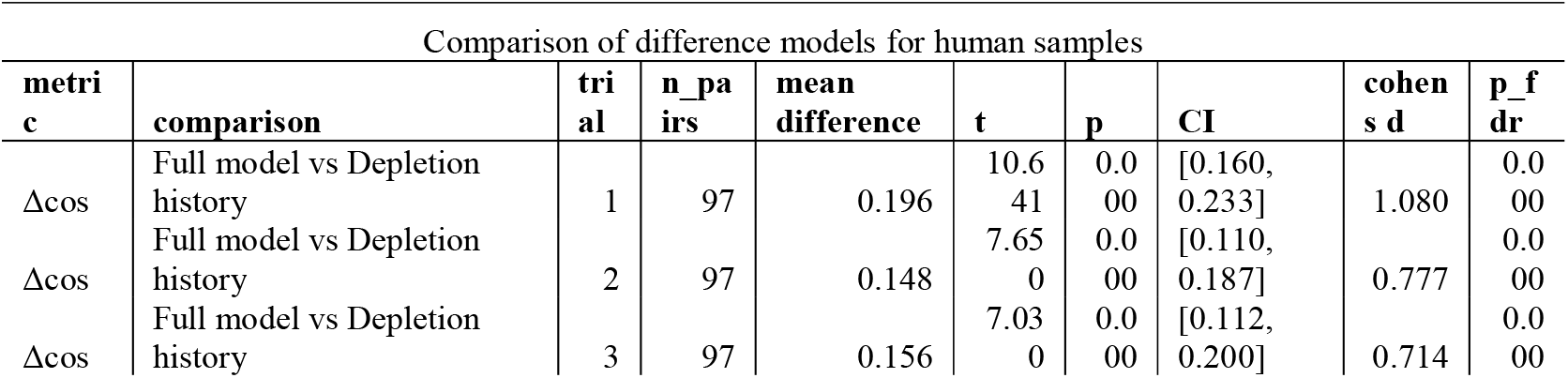
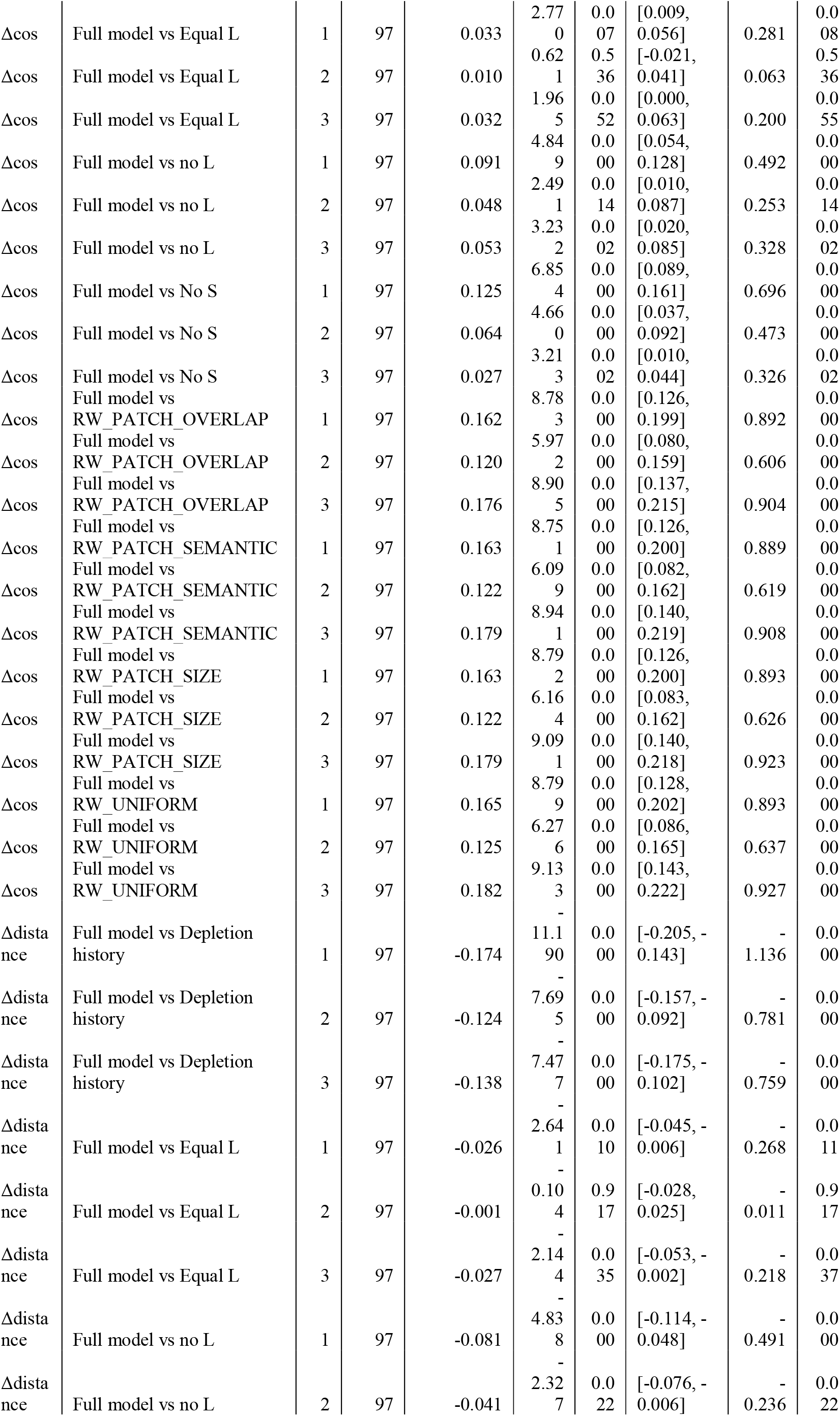
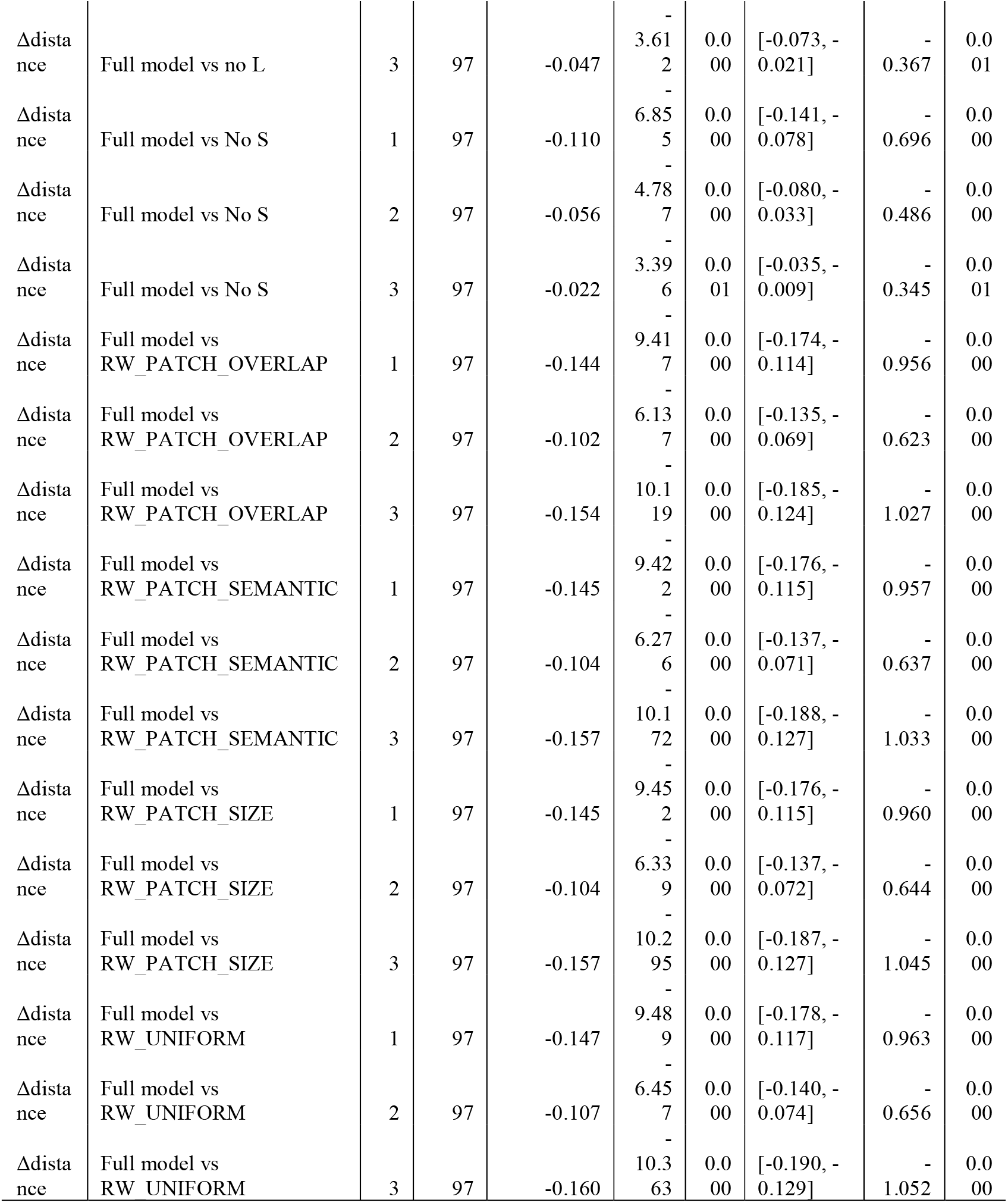
comparison between different models for human samples.

**Supplementary Table 9:**
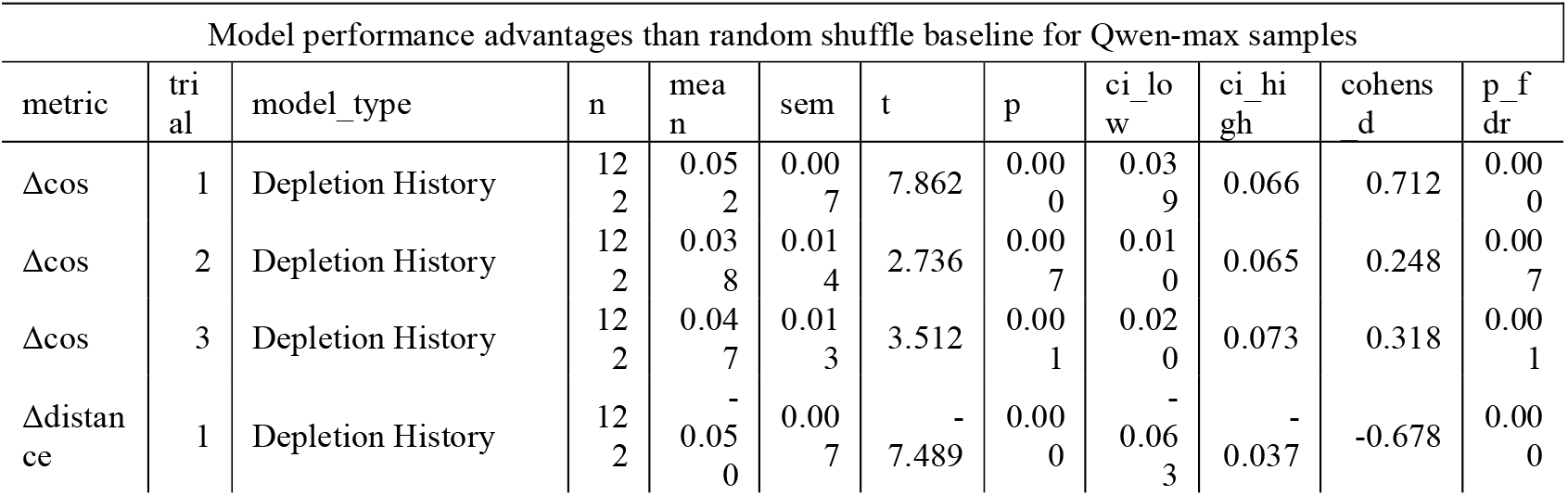
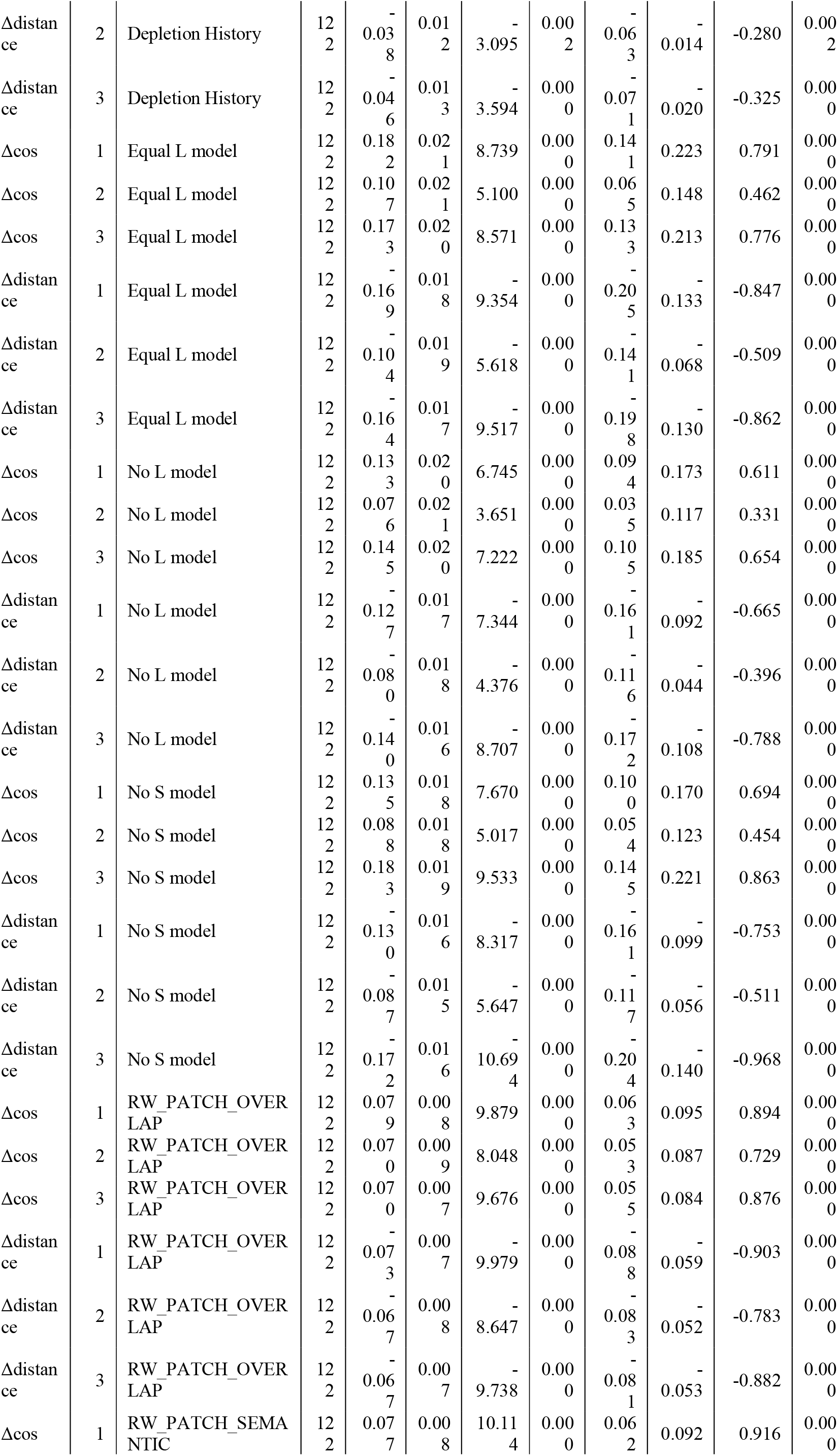
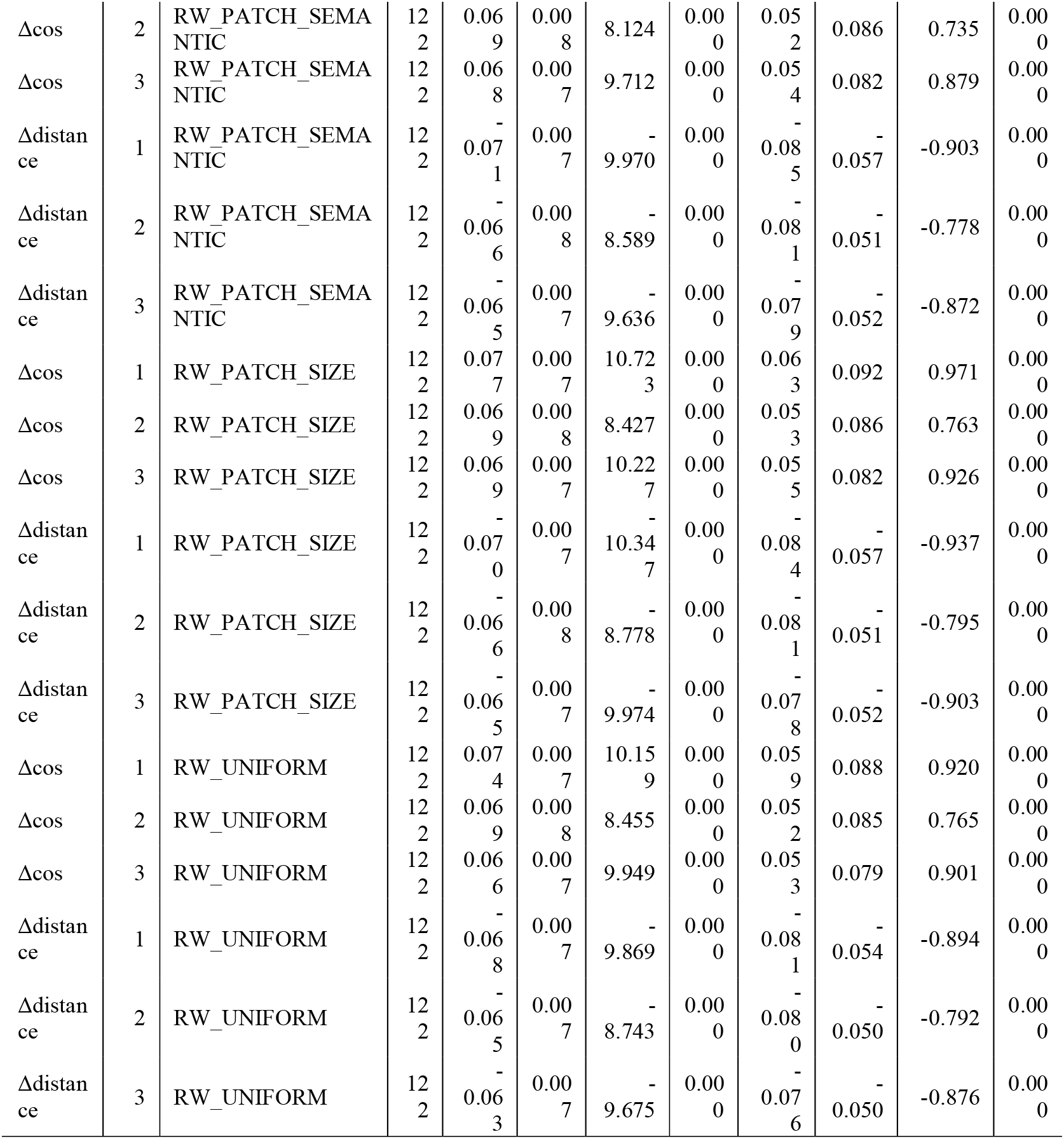
differences between control models and random shuffle baseline in Qwen-max data. All control models outperformed than random shuffle baseline.

**Supplementary Table 10:**
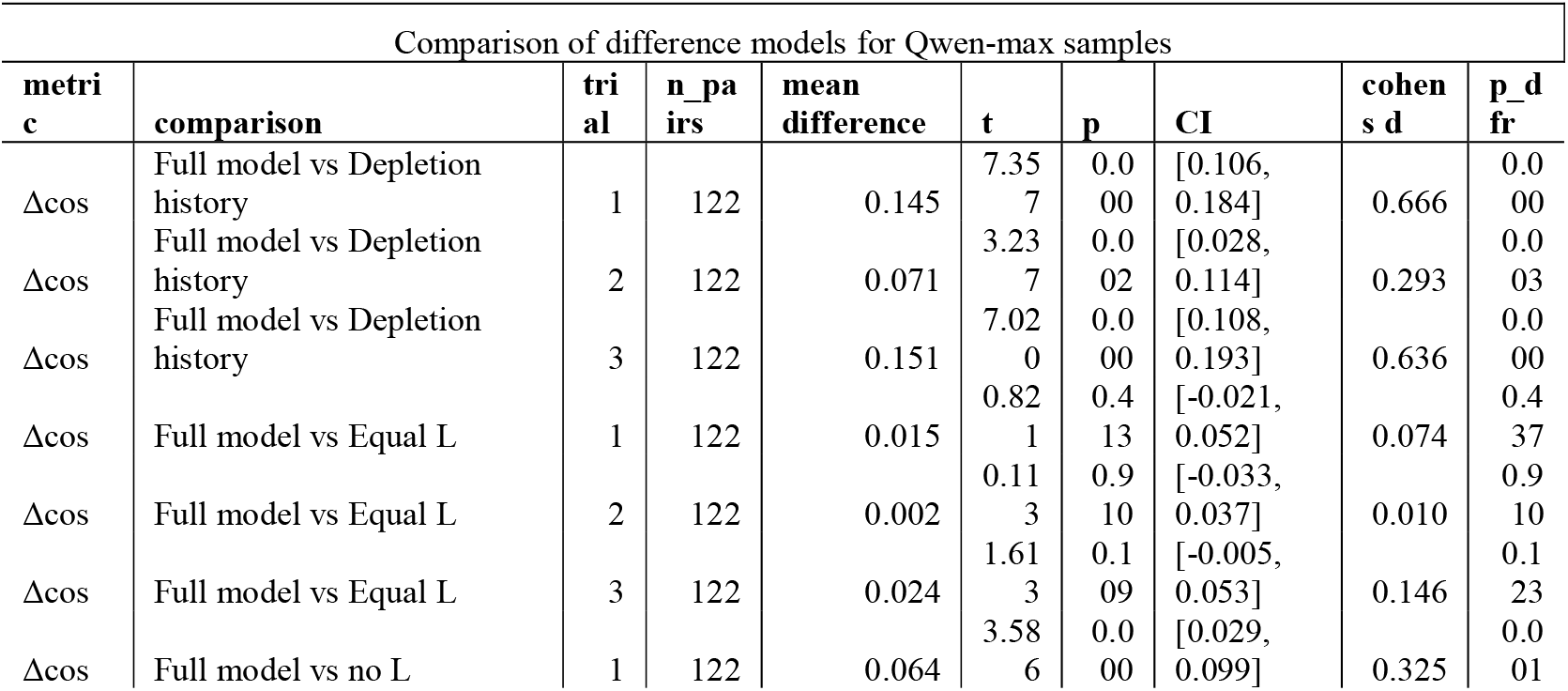
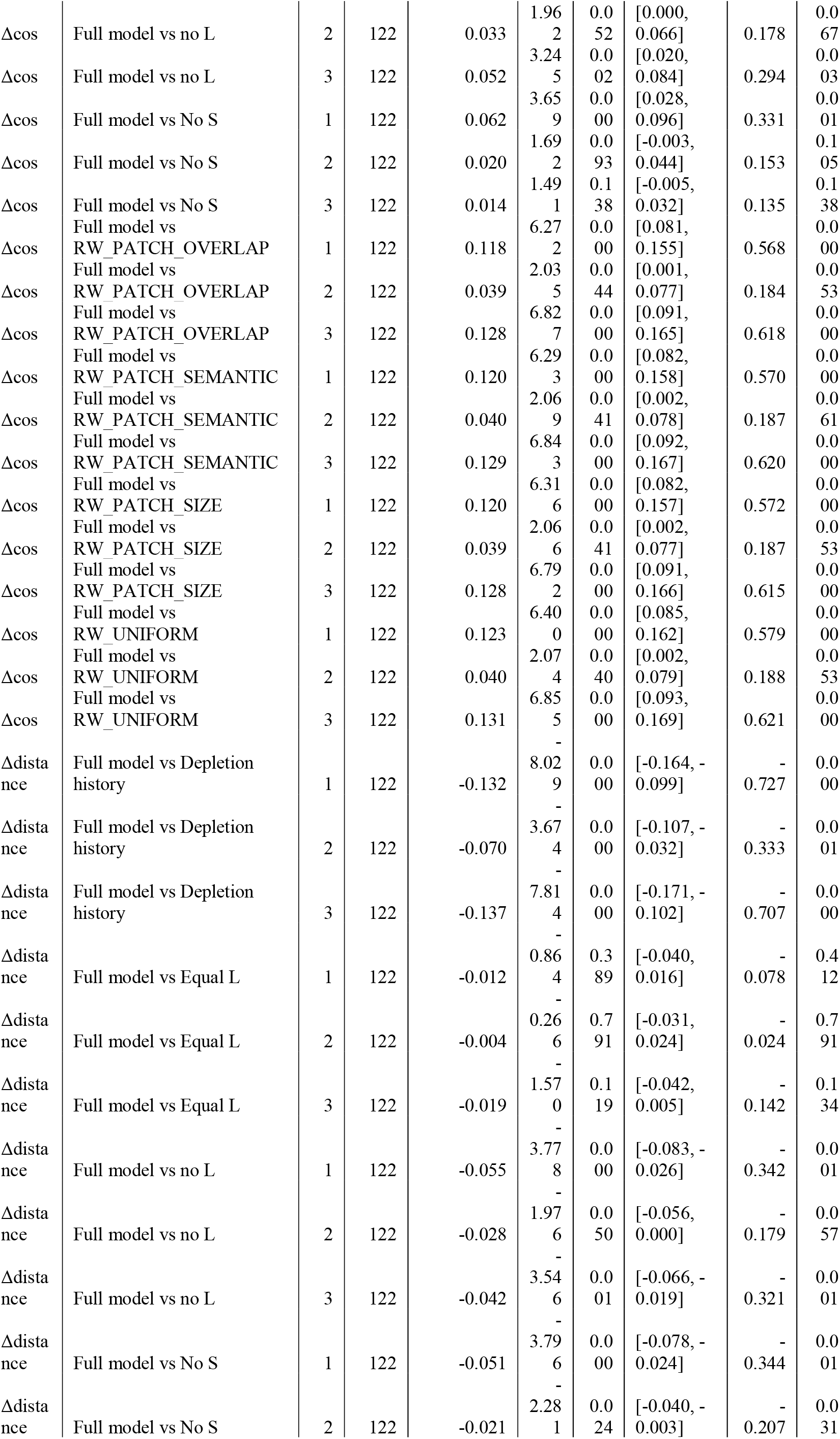
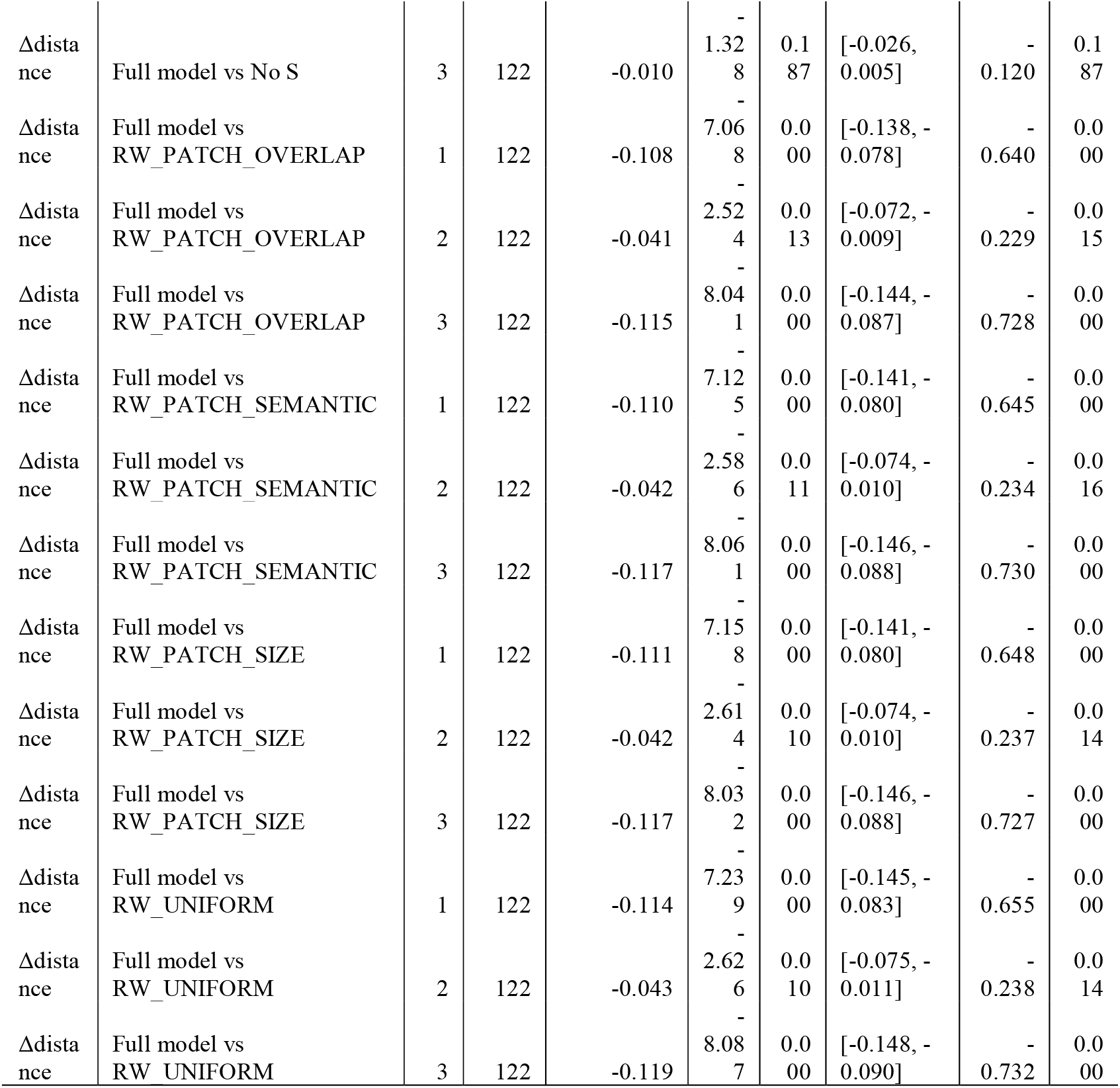
comparison between different models for Qwen-max samples.

**Supplementary Table 11:**
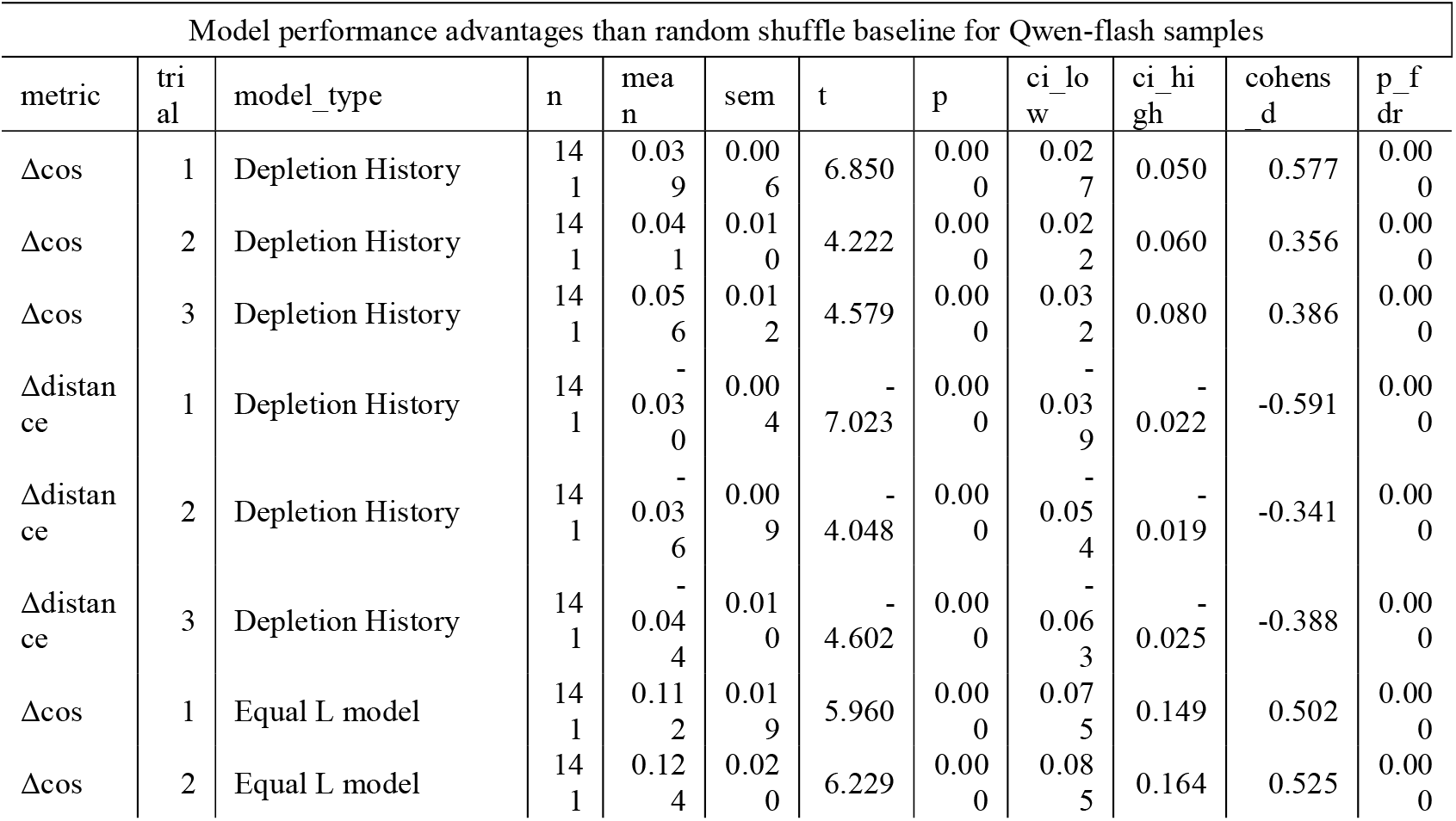
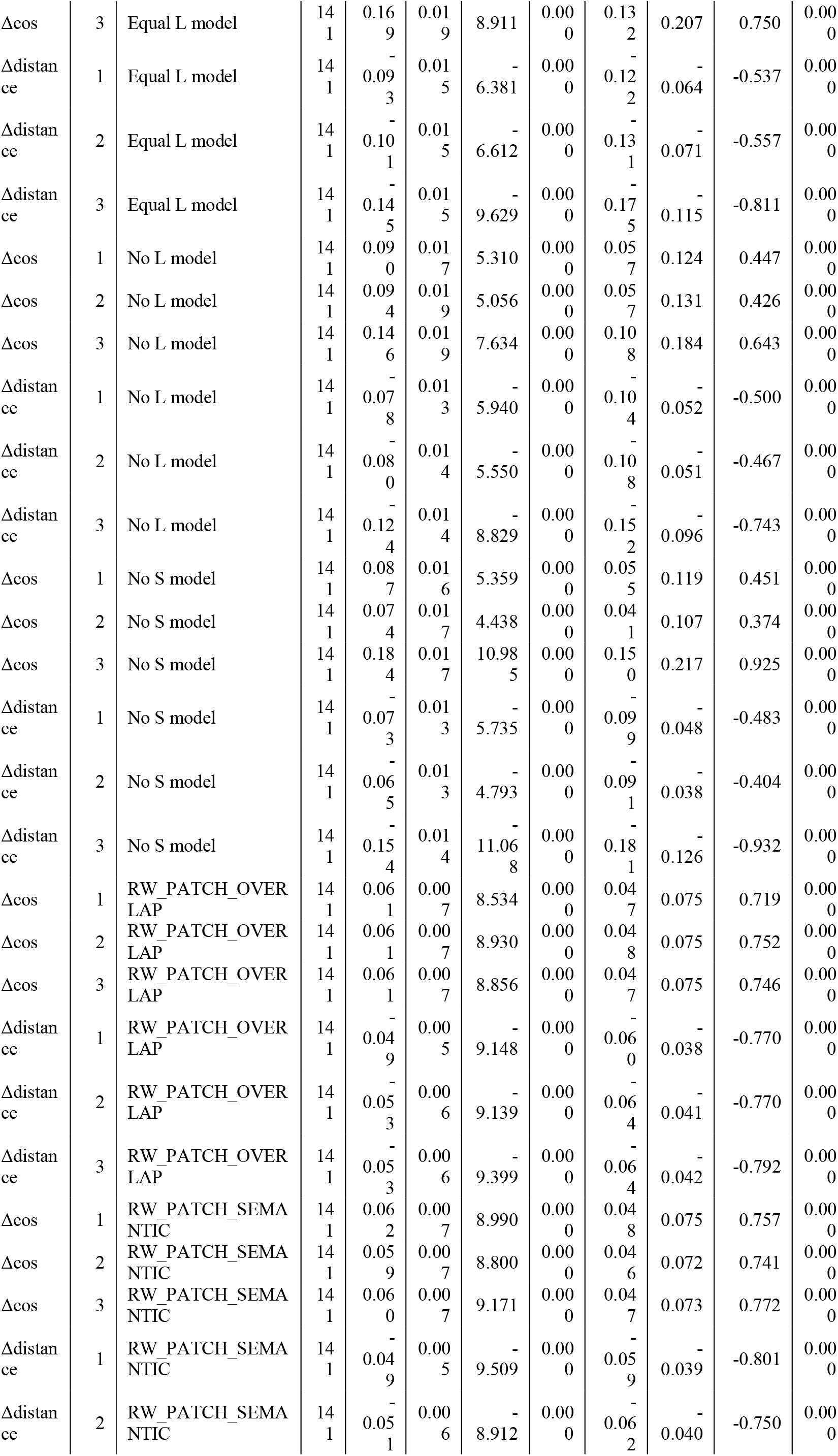
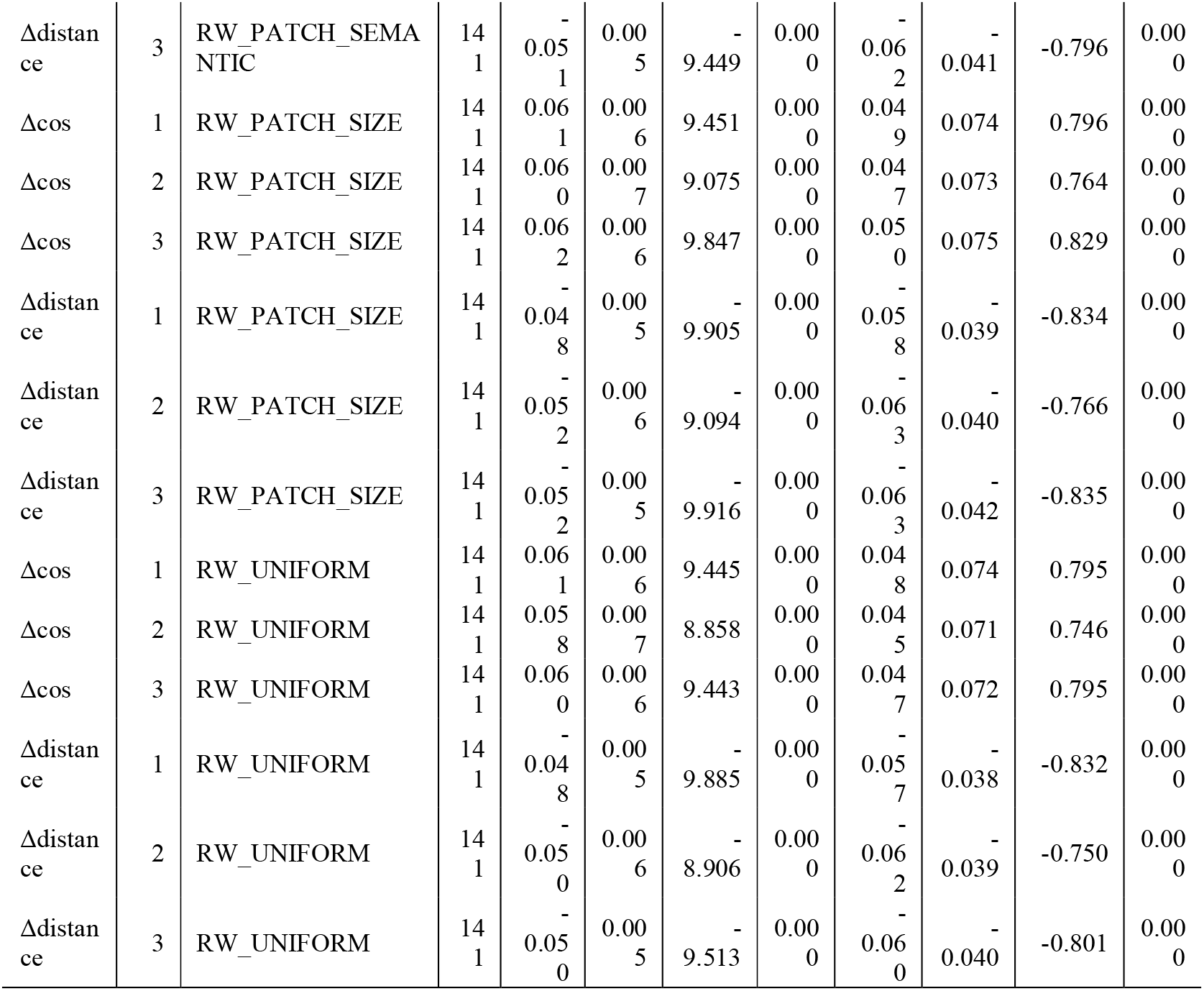
differences between control models and random shuffle baseline in Qwen-flash data. All control models outperformed than random shuffle baseline.

**Supplementary Table 12:**
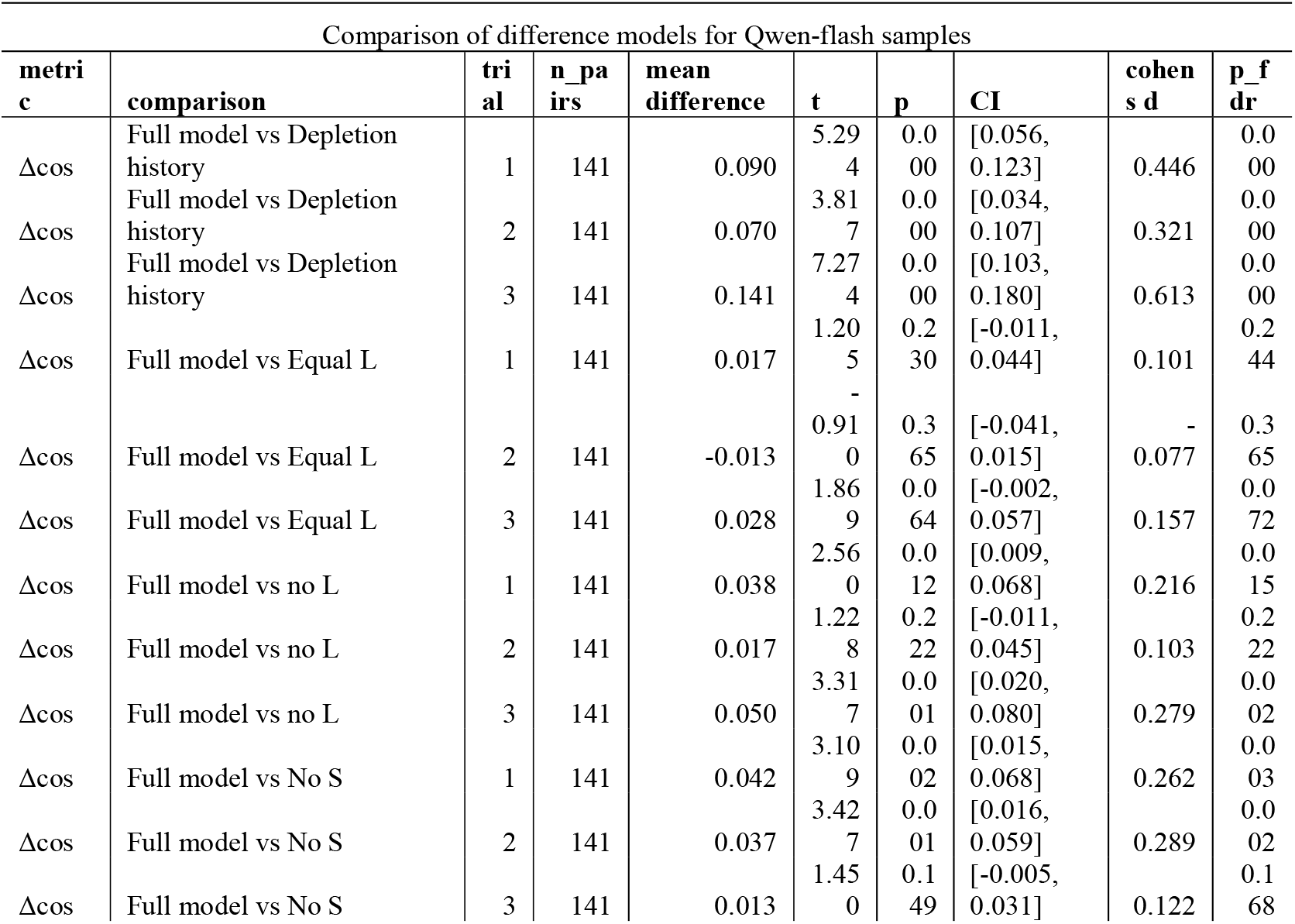
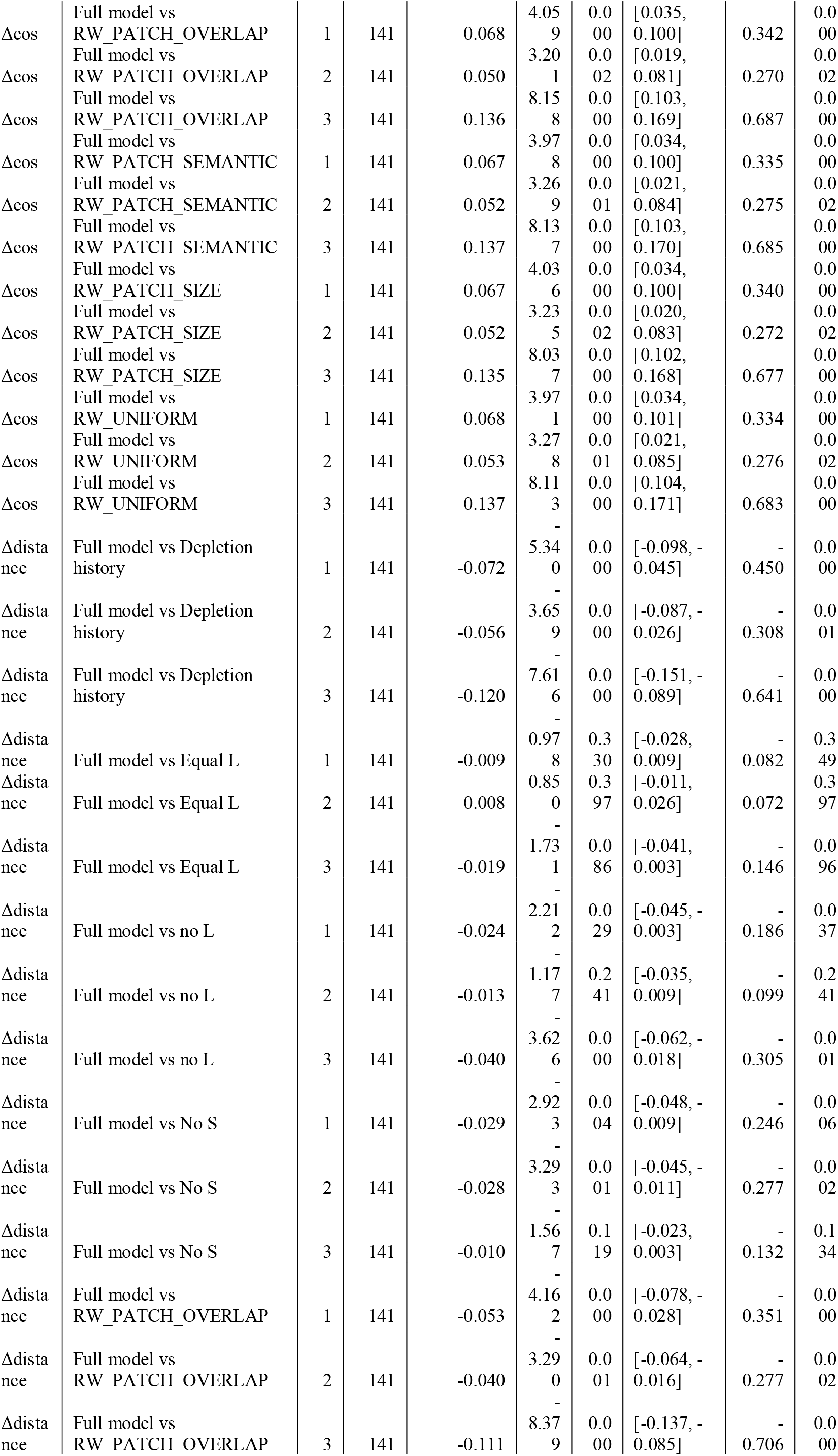
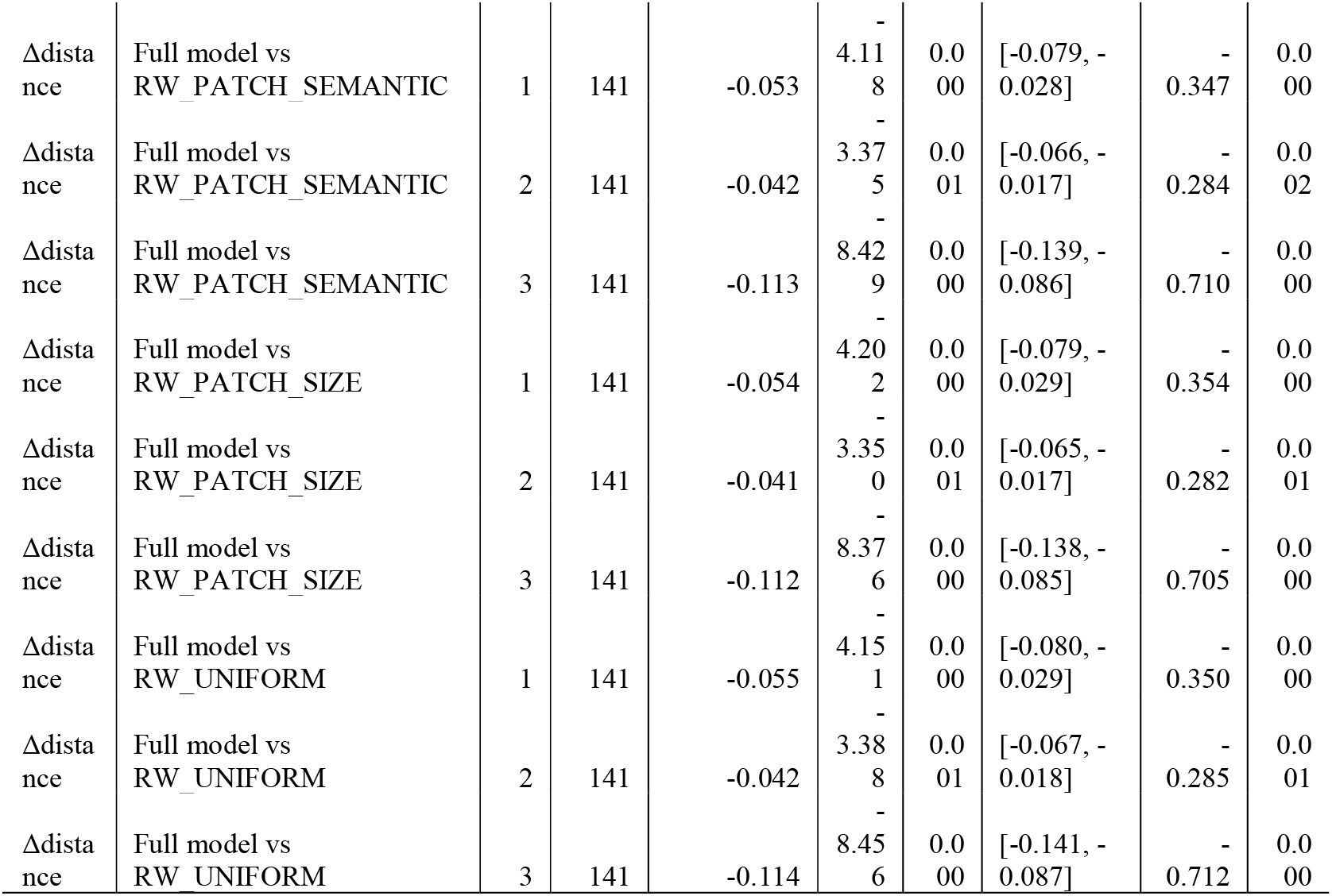
comparison between different models for Qwen-flash samples.

**Supplementary Figure S1.**
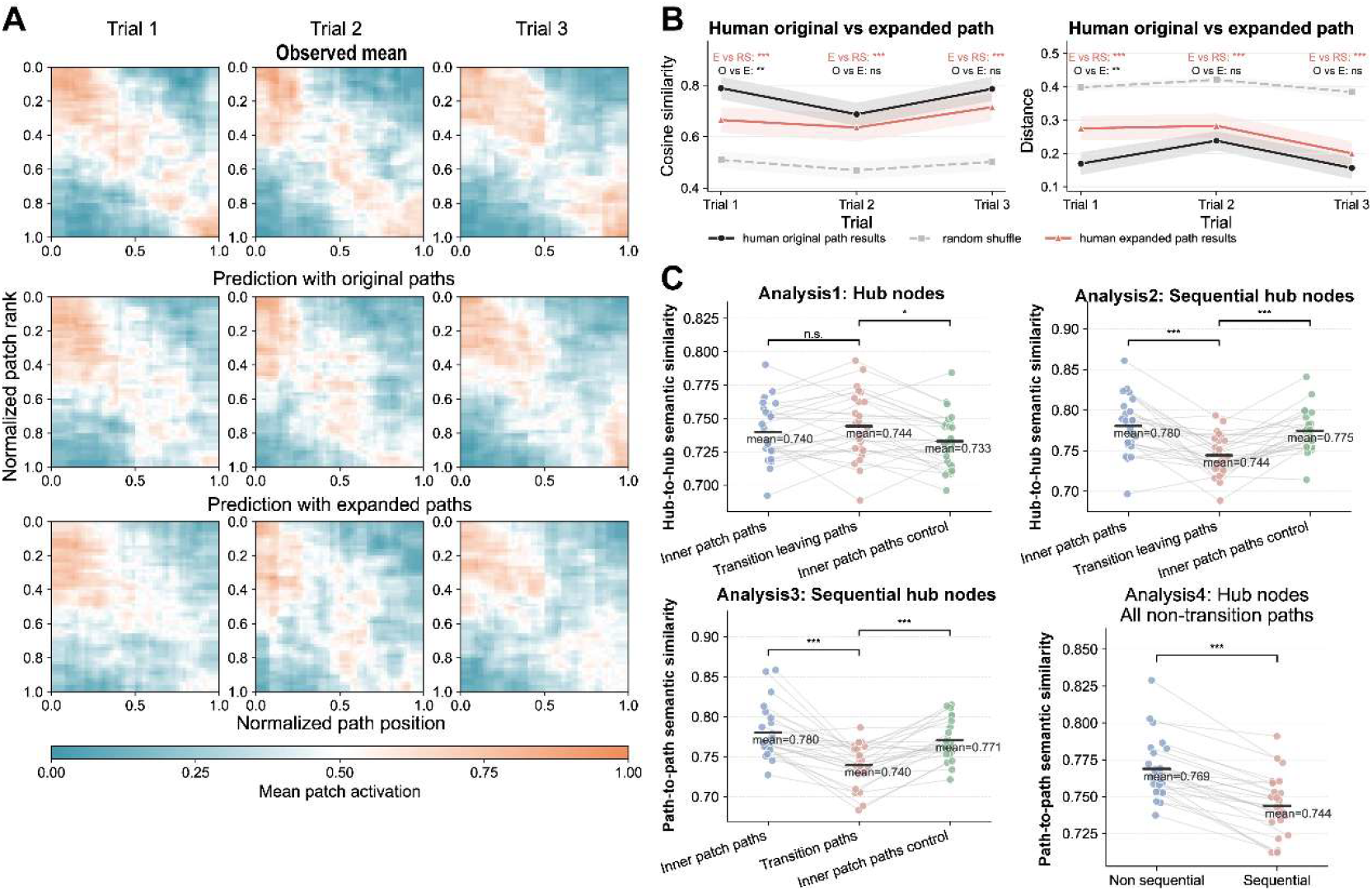
Model-predicted performance in the supplemental two-stage narrative experiment. (A) Normalized and resampled patch activation matrices. Compared with the Phase 1 model (middle), the Phase 2 model incorporating candidate memory resources (bottom) also captured the temporal organization of the human thought dynamics observed in Phase 1 (top), supporting the robustness of the MVT-based model’s predictive performance. (B) Model performance in the supplemental two-stage narrative experiment. Both the Phase 1 and Phase 2 models outperformed the random shuffle baseline, although the Phase 2 model performed significantly worse than the Phase 1 model in trial 1. (C) Results of Analyses 1–4 after expanded paths were incorporated. The model continued to show reduced semantic similarity at transitions in Analyses 2 and 3, greater non-temporal semantic change for transition-related paths than for inner-patch paths in Analysis 1, and lower cross-semantic similarity for sequential than for non-sequential paths in Analysis 4. The only exception was the nonsignificant difference between inner-patch paths and transition-leaving paths in Analysis 1. These results were broadly consistent with the main analyses. Detailed statistical results are provided in Supplementary Table 4. ***p < 0.001.

**Supplementary Figure S2.**
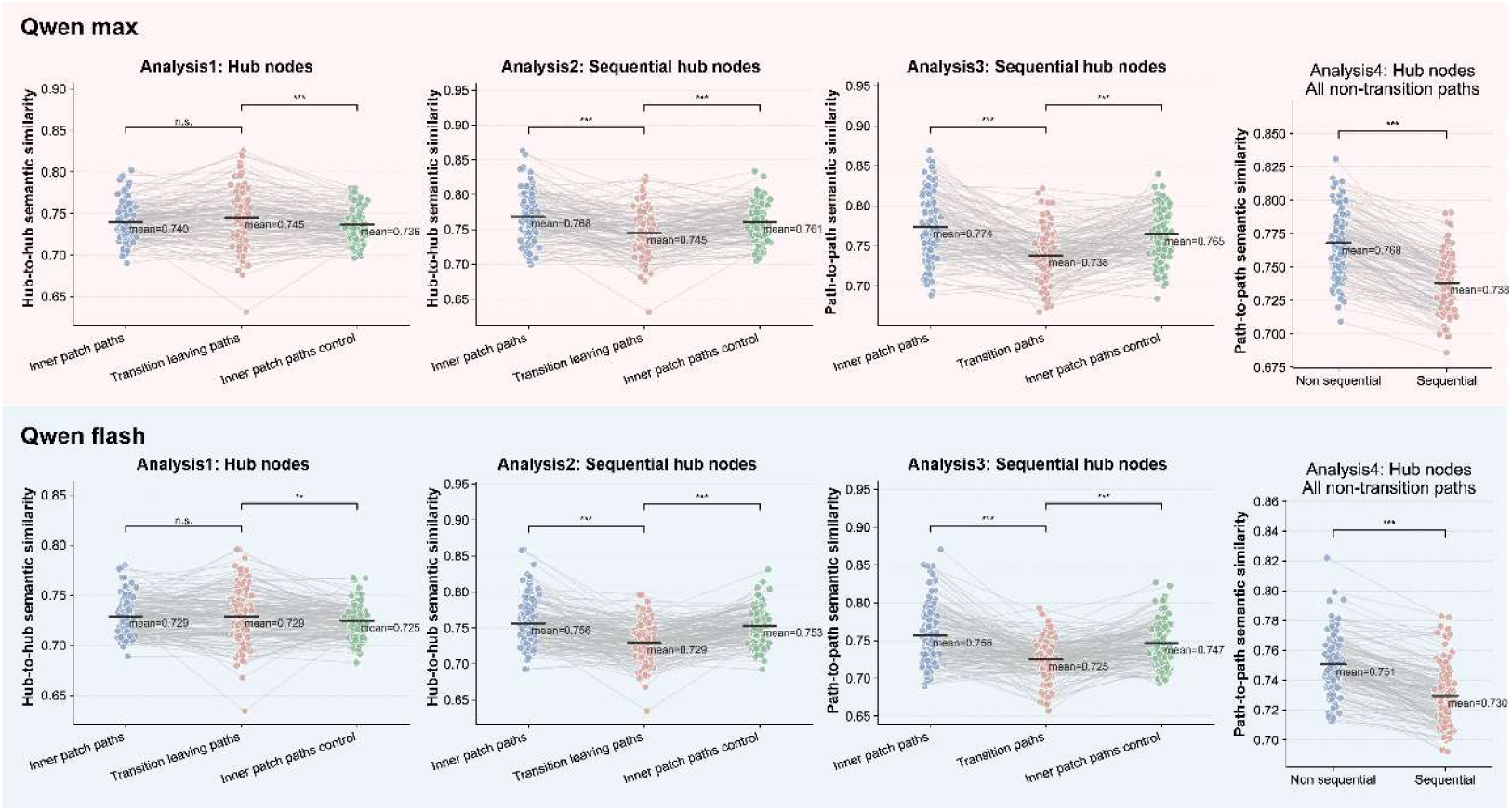
Results of transition-related semantic Analyses 1–4 for Qwen samples. Results are shown for Qwen-max (top) and Qwen-flash (bottom). The semantic-similarity differences between inner-patch and transition-related paths, as well as between sequential and non-sequential paths, were broadly consistent with those observed in the human samples. Detailed statistical results are provided in Supplementary Tables 5 and 6. ***p < 0.001.

**Supplementary Figure S3.**
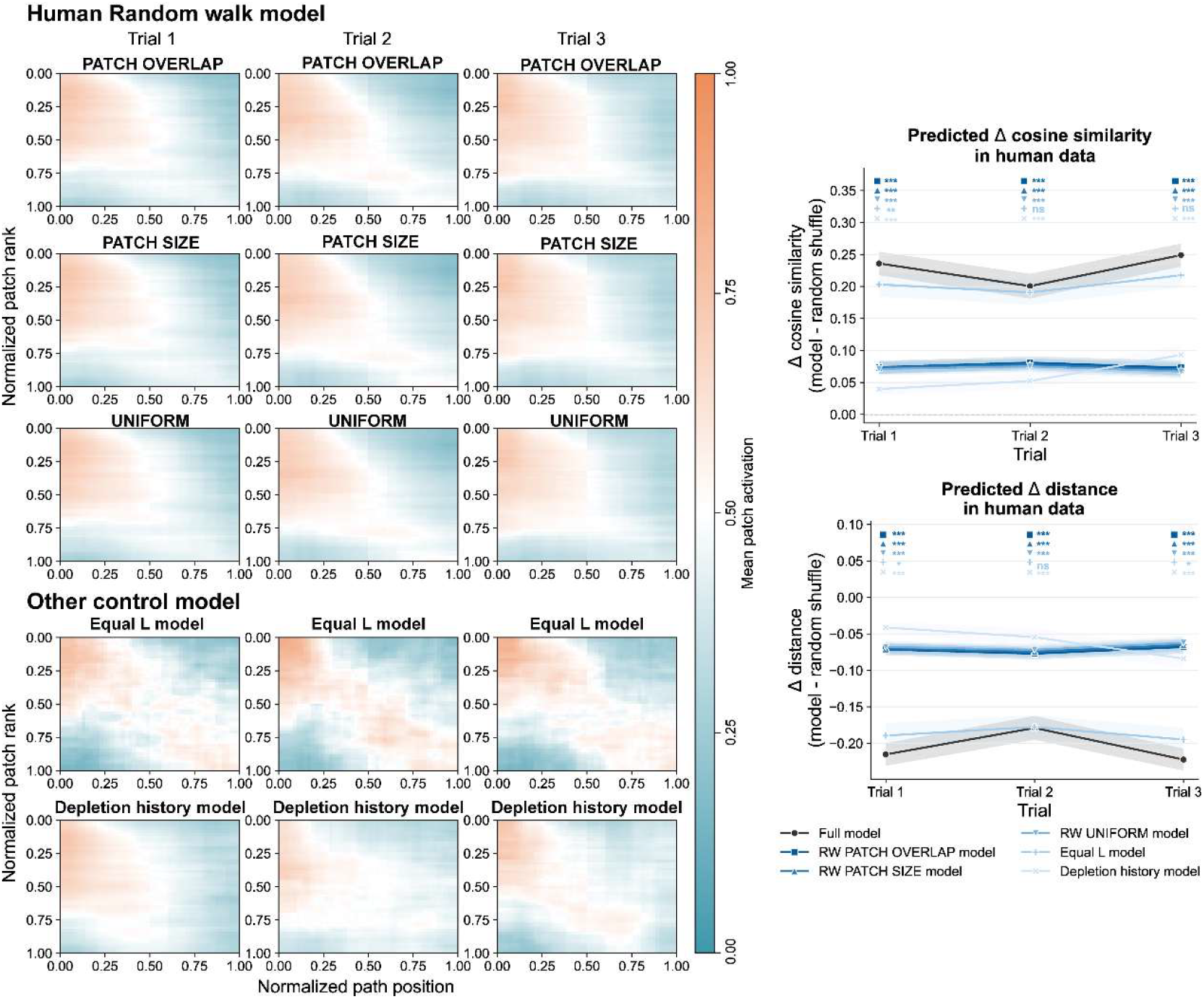
Predictive performance of additional control models for the human samples. Left: In addition to the patch semantic random walk reported in the main text, we tested random-walk models in which patch-to-patch transition probabilities were determined by patch overlap (PATCH OVERLAP), the number of paths contained in the target patch (PATCH SIZE), or a uniform probability distribution (UNIFORM). None of these random walk models fully captured the dynamics of the observed activation matrices, particularly during the later portions of the path sequences. In addition to the No S, No L, and Depletion history models reported in the main text, we also tested an Equal L model, in which the weights applied to L values were set to be equal. This model captured the dynamics of the observed activation matrices. Right: Performance of the control models shown on the left. Although all control models outperformed the random-shuffle baseline, all models except the Equal L model were significantly less predictive than the full model. The Equal L model showed performance close to that of the full model. Detailed statistical results are provided in Supplementary Tables 7 and 8. *p < 0.05, **p < 0.01, ***p < 0.001.

**Supplementary Figure S4.**
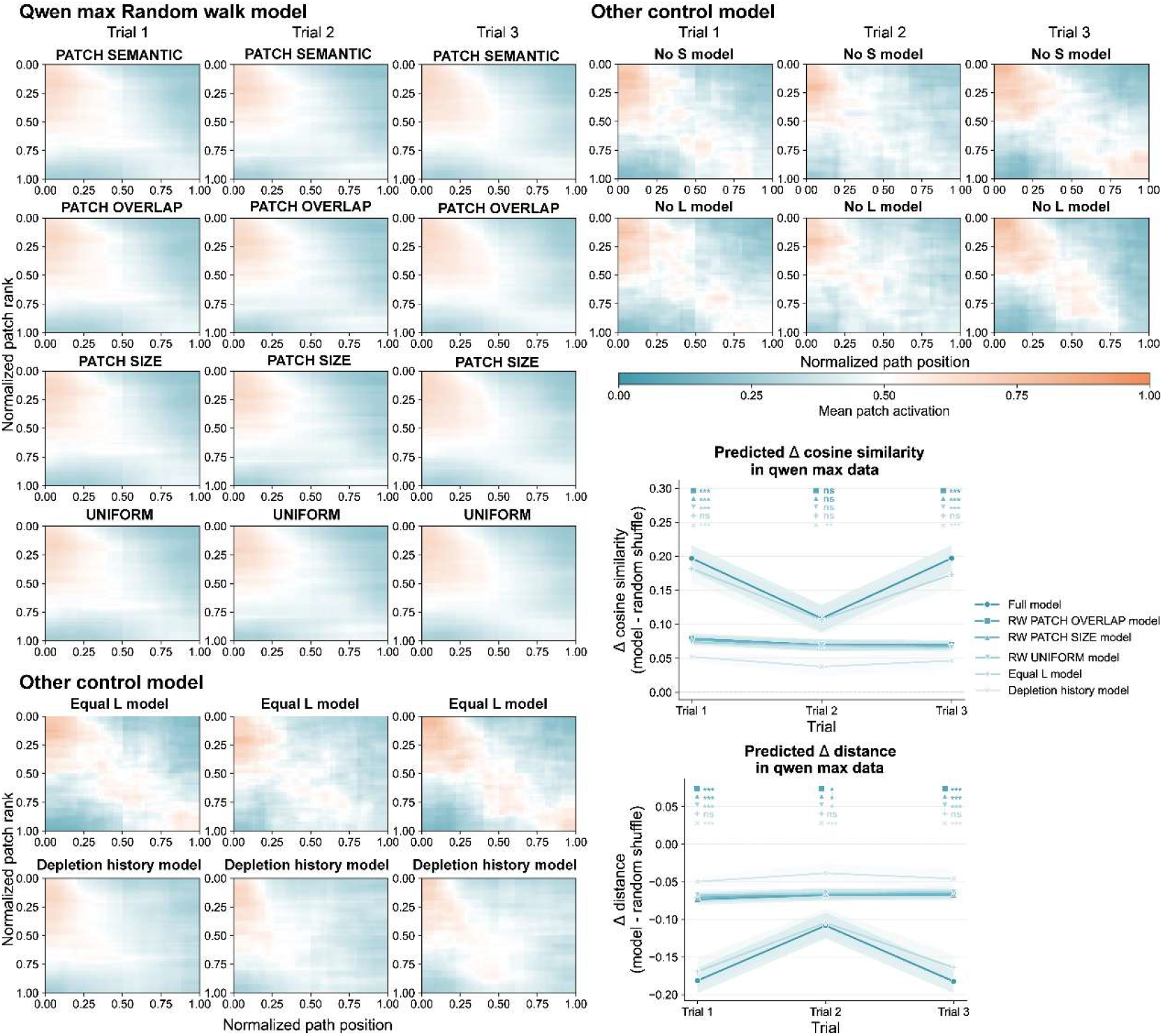
Predictive performance of all control models for Qwen-max samples. The PATCH SEMANTIC random walk model corresponds to the random walk model reported in the main text. In trial 2, all control models except the Depletion history model showed performance close to that of the full model for Qwen-max samples. Detailed statistical results are provided in Supplementary Tables 9 and 10. *p < 0.05, **p < 0.01, ***p < 0.001.

**Supplementary Figure S5.**
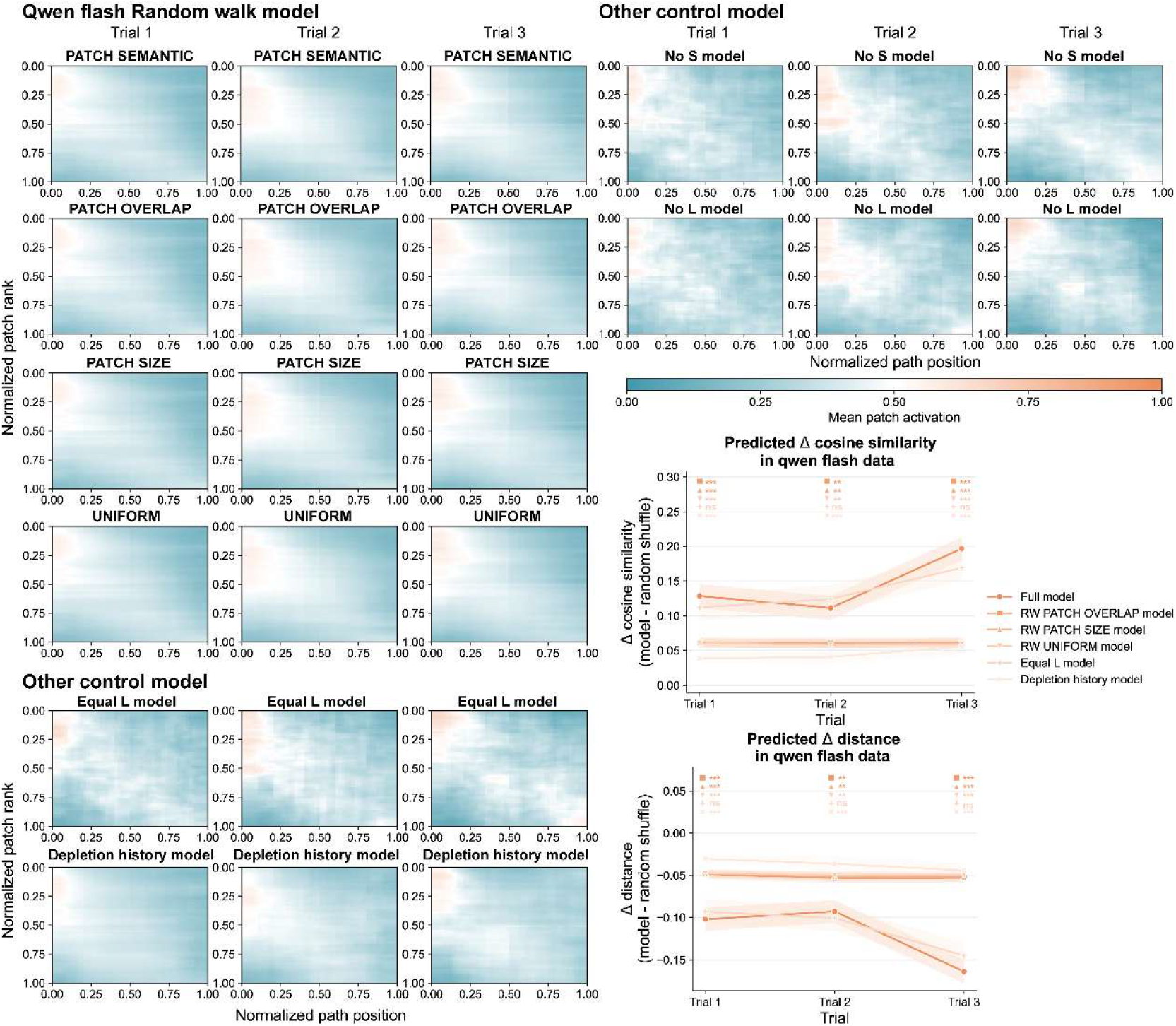
Predictive performance of all control models for Qwen-flash samples. In addition to the No S and No L models reported in the main text, which showed performance close to that of the full model in trials 2 and 3, the Equal L model did not differ significantly from the full model in any trial. Detailed statistical results are provided in Supplementary Tables 11 and 12. **p < 0.01, ***p < 0.001.

